# Protein generation with evolutionary diffusion: sequence is all you need

**DOI:** 10.1101/2023.09.11.556673

**Authors:** Sarah Alamdari, Nitya Thakkar, Rianne van den Berg, Neil Tenenholtz, Robert Strome, Alan M. Moses, Alex X. Lu, Nicolò Fusi, Ava P. Amini, Kevin K. Yang

## Abstract

Deep generative models are increasingly powerful tools for the *in silico* design of novel proteins. Recently, a family of generative models called diffusion models has demonstrated the ability to generate biologically plausible proteins that are dissimilar to any actual proteins seen in nature, enabling unprecedented capability and control in *de novo* protein design. However, current state-of-the-art diffusion models generate protein structures, which limits the scope of their training data and restricts generations to a small and biased subset of protein design space. Here, we introduce a general-purpose diffusion framework, EvoDiff, that combines evolutionary-scale data with the distinct conditioning capabilities of diffusion models for controllable protein generation in sequence space. EvoDiff generates high-fidelity, diverse, and structurally-plausible proteins that cover natural sequence and functional space. We show experimentally that EvoDiff generations express, fold, and exhibit expected secondary structure elements. Critically, EvoDiff can generate proteins inaccessible to structure-based models, such as those with disordered regions, while maintaining the ability to design scaffolds for functional structural motifs. We validate the universality of our sequence-based formulation by experimentally characterizing intrinsically-disordered mitochondrial targeting signals, metal-binding proteins, and protein binders designed using EvoDiff. We envision that EvoDiff will expand capabilities in protein engineering beyond the structure-function paradigm toward programmable, sequence-first design.

---

Evolution has yielded a diversity of functional proteins that precisely modulate cellular processes. Recent years have seen the emergence of deep generative models that aim to learn from this diversity to generate proteins that are both valid and novel, with the ultimate goal of then tailoring function to solve outstanding modern-day challenges, such as the rapid development of targeted therapeutics and vaccines or engineered enzymes for the degradation of industrial waste (**Fig. 1A**) (*1, 2*). Diffusion models provide a particularly powerful framework for generative modeling of novel proteins, as they generate high-diversity samples and can be conditioned given a wide variety of inputs or design objectives (*3–6*). Indeed, today’s most biologically-plausible instances of *in silico*-designed proteins come from diffusion models of protein *structure* (*7–15*).

**Figure 1:**
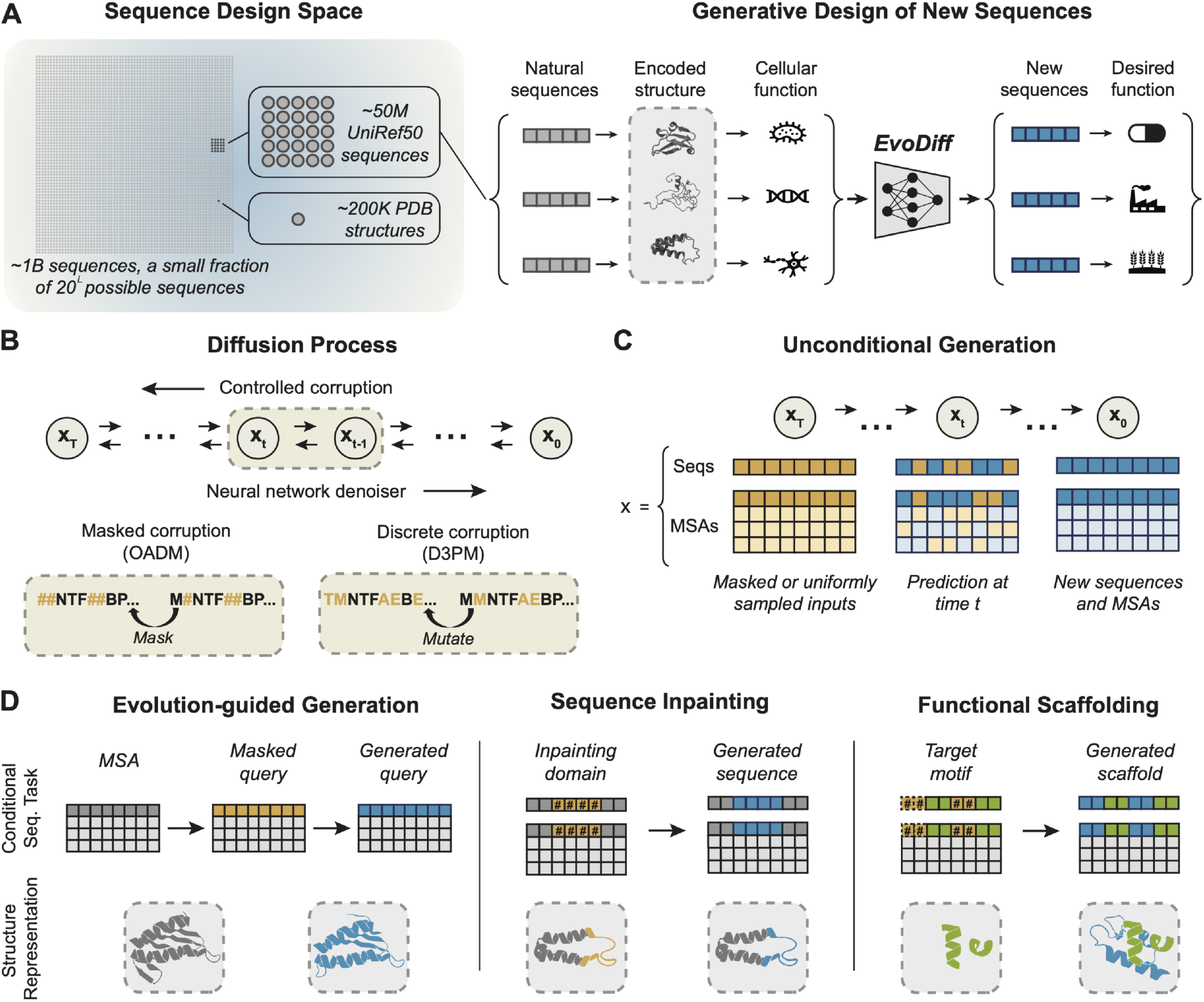
Protein sequence generation with evolutionary diffusion. **(A)** (Left) Evolution has sampled a tiny fraction of possible protein sequences. Experimental structures have been determined for even fewer proteins. (Right) EvoDiff is a generative discrete diffusion model trained on natural protein sequences. Sampling from EvoDiff yields new protein sequences that may perform desired functions. **(B)** Discrete diffusion models consist of controlled corruption and learned denoising processes. In the masked corruption process, input tokens are masked in an order-agnostic fashion (bottom, left). In the discrete corruption process, inputs are corrupted via a Markov process controlled by a transition matrix capturing amino acid mutation frequencies (bottom, right). **(C)** EvoDiff enables unconditional generation of protein sequences or MSAs. Starting from masked or uniformly sampled inputs *x*_*T*_, EvoDiff generates new sequences or MSAs by reversing the corruption process, iteratively denoising *x*_*t*_ into realistic sequences or MSAs *x*_*0*_. **(D)** Controllable protein design with EvoDiff, via conditioning on evolutionary information encoded in MSAs (left); inpainting functional domains from masked portions of a sequence (middle); or scaffolding structural motifs without explicit structural information (right). thermore, EvoDiff-OADM is the only model variant where performance scales with increased model size (**Table S1; Fig. S1**).

These models – including the current state-of-the-art approach RFdiffusion (*10*) – fit in the structure-based protein design paradigm of first generating a structure that fulfills desired constraints and then designing a sequence that will fold to that structure. However, sequence, not structure, is the universal design space for proteins. Every protein is completely defined by its amino-acid sequence. We discover proteins by finding their coding sequences in genomes, and proteins are synthesized as amino-acid sequences. Sequence then determines function through both an ensemble of structural conformations and the chemistry enabled by the amino acids themselves. However, not every protein folds into a static structure. In these cases, structure-based design is not viable because the function is not mediated by a static structure (*16–18*), with the most extreme examples being intrinsically disordered regions (IDRs) (*19*). Therefore, static structures characterized by X-ray crystallography are an incomplete distillation of the information captured in sequence space (*20–23*). Furthermore, structural data (ca. 200k solved structures in PDB) is scarce and unrepresentative of the full diversity of natural sequences (ca. billions of unique natural protein sequences; **Fig. 1A**), inherently limiting the capacity of any structure-based generative model to learn the full diversity of protein functional space.

We combine evolutionary-scale datasets with diffusion models to develop a powerful new generative modeling framework, which we term EvoDiff, for controllable protein design from sequence data alone (**Fig. 1**). Given the natural framing of proteins as sequences of discrete tokens over an amino acid language, we use a *discrete* diffusion framework in which a forward process iteratively corrupts a protein sequence by changing its amino acid identities, and a learned reverse process, parameterized by a neural network, predicts the changes made at each iteration (**Fig. 1B**). The reverse process can then be used to generate new protein sequences starting from random noise (**Fig. 1C**). Importantly, EvoDiff’s discrete diffusion formulation is mathematically distinct from continuous diffusion formulations previously used for protein structure design (*7–15*). Beyond evolutionary-scale datasets of single protein sequences, multiple sequence alignments (MSAs) inherently capture evolutionary relationships by revealing patterns of conservation and variation in the amino acid sequences of sets of related proteins. We thus additionally build discrete diffusion models trained on MSAs to leverage this additional layer of evolutionary information to generate new single sequences (**Fig. 1C-D**).

We evaluate our sequence and MSA models – EvoDiff-Seq and EvoDiff-MSA, respectively – across a range of generation tasks to demonstrate their power for controllable protein design (**Fig. 1D**). We first show that EvoDiff-Seq unconditionally generates high-quality, diverse proteins that capture the natural distribution of protein sequence, structural, and functional space. Unconditional sampling from EvoDiff can yield proteins that stably express and exhibit expected biophysical properties. Using EvoDiff-MSA, we achieve evolution-guided design of novel sequences conditioned on an alignment of evolutionarily-related, but distinct, proteins. Finally, by exploiting the conditioning capabilities of our diffusion-based modeling framework and its grounding in a universal design space, we demonstrate – through both *in silico* and wetlab experiments – that EvoDiff can reliably generate proteins with functional IDRs, directly overcoming a key limitation of structure-based generative models, and generate scaffolds for functional structural motifs without any explicit structural or binder information.

## Discrete diffusion models of protein sequence

EvoDiff is the first generative diffusion model for protein design trained on evolutionary-scale protein sequence data. We investigated two types of forward processes for diffusion over discrete data modalities (*24, 25*) to determine which would be most effective (**Fig. 1B**). In order-agnostic autoregressive diffusion (EvoDiff-OADM, see Methods) (*24*), one amino acid is converted to a special mask token at each step in the forward process (**Fig. 1B**). After *T* =*L* steps, where *L* is the length of the sequence, the entire sequence is masked. We additionally designed discrete denoising diffusion probabilistic models (EvoDiff-D3PM, see Methods) (*25*) for protein sequences. In EvoDiff-D3PM, the forward process corrupts sequences by sampling mutations according to a transition matrix, such that after *T* steps the sequence is indistinguishable from a uniform sample over the amino acids (**Fig. 1B**). In the reverse process for both, a neural network model is trained to undo the previous corruption. The trained model can then generate new sequences starting from sequences of masked tokens or of uniformly-sampled amino acids for EvoDiff-OADM or EvoDiff-D3PM, respectively (**Fig. 1C**).

To facilitate direct and quantitative model comparisons, we trained all EvoDiff sequence models on 42M sequences from UniRef50 (*26*) using a dilated convolutional neural network architecture introduced in the CARP protein masked language model (*27*). We trained 38M-parameter and 640M-parameter versions for each forward corruption scheme to test the effect of model size on model performance. As a first evaluation of our EvoDiff sequence models, we calculated each model’s test-set perplexity, which reflects its ability to capture the distribution of natural sequences and generalize to unseen sequences (see Methods). We observe that EvoDiff-OADM learns to reconstruct the test set more accurately than two tested EvoDiff-D3PM variants employing uniform and BLOSUM62-based transition matrices (**Table S1; Fig. S1**). Fur-

To explicitly leverage evolutionary information, we designed and trained EvoDiff MSA models using the MSA Transformer (*28*) architecture on the OpenFold dataset (*29*). To do so, we subsampled MSAs to a length of 512 residues per sequence and a depth of 64 sequences, either by randomly sampling the sequences (“Random”) or by greedily maximizing for sequence diversity (“Max”). Within each subsampling strategy, we then trained EvoDiff MSA models with the OADM and D3PM corruption schemes. OADM corruption results in the lowest validation set perplexities, indicating that OADM models are best able to generalize to new MSAs (**Table S2; Fig. S2**). To select a subsampling method, we compared the ability of each model to reconstruct validation set MSAs, finding that maximizing for sequence diversity yields improved performance no matter how the validation MSAs are subsampled (**Table S2**). We thus selected the OADM-Max model for downstream analysis, hereafter referring to it as EvoDiff-MSA.

### Structural plausibility of generated sequences

We next investigated whether EvoDiff could generate new protein sequences that were individually valid and structurally plausible. To assess this, we developed a workflow that evaluates the foldability and self-consistency of sequences generated by EvoDiff (**Fig. 2A**). We generated 1000 sequences from each EvoDiff sequence model with lengths drawn from the empirical distribution of lengths in the training set. We compared EvoDiff’s generations to sequences generated from a left-to-right autoregressive language model (LRAR) with the same architecture and training set as EvoDiff and to sequences generated from protein masked language models such as ESM-2 (*30*) (**Figs. 2B-C, S3, S4; Table S3**).

**Figure 2:**
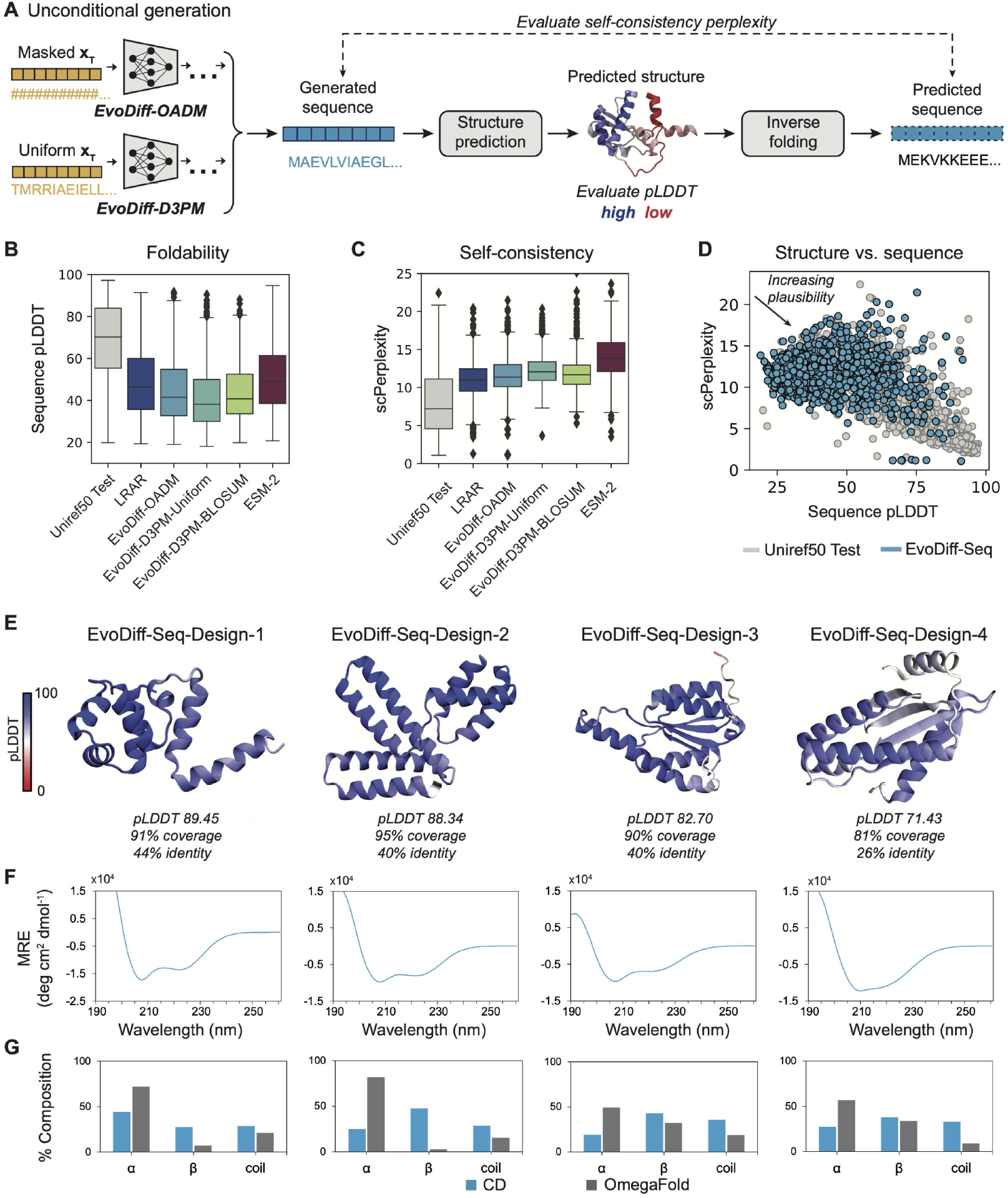
EvoDiff generates realistic and structurally-plausible protein sequences. **(A)** Workflow for evaluating the foldability and self-consistency of sequences generated by EvoDiff sequence models. **(B-C)** Distributions of foldability, measured by sequence pLDDT of predicted structures (B), and self-consistency, measured by scPerplexity (C), for sequences from the test set, EvoDiff models, and baselines (*n*=1000 sequences per model; box plots show median and interquartile range). **(D)** Sequence pLDDT versus scPerplexity for sequences from the test set (grey, *n*=1000) and the 640M-parameter OADM model EvoDiff-Seq (blue, *n*=1000). **(E)** Predicted structures and metrics for successfully expressed and characterized unconditional generations from EvoDiff-Seq, the 640M-parameter OADM model. OmegaFold predictions, colored by pLDDT, are shown, and the average pLDDT for each structure is reported. % coverage and % identity to the top BLAST hit is denoted below each design. **(F)** Circular dichroism (CD) spectra for designed sequences from (E). **(G)** Structural composition of each sequence design as inferred from CD spectra (blue) versus from OmegaFold (grey). AlphaFold predictions are included in **Fig. S6** for comparison.

We assessed the foldability of individual sequences by predicting their corresponding structures using OmegaFold (*31*) and computing the average predicted local distance difference test (pLDDT) across the whole structure (**Fig. 2B**). pLDDT reflects OmegaFold’s confidence in its structure prediction for each residue. In addition to the average pLDDT across a whole protein, we observe that pLDDT scores can vary significantly across a protein sequence (**Fig. S5 B**). It is important to note that while pLDDT scores above 70 are often considered to indicate high prediction confidence, low pLDDT scores can be consistent with intrinsically disordered regions (IDRs) of proteins (*32*), which are found in many natural proteins. As an additional metric of structural plausibility, we computed a self-consistency perplexity (scPerplexity) by redesigning each predicted structure with the inverse folding algorithm ESM-IF (*33*) and computing the perplexity against the original generated sequence (**Fig. 2A, C; Table S3**). Given that ESM-IF and EvoDiff were both trained on UniRef50 data, it is possible that sequences from EvoDiff’s validation set overlap with sequences in the ESM-IF train set; thus we performed the same self-consistency evaluations using ProteinMPNN (*34*), which is not trained on UniRef50, for inverse folding (**Table S3**).

While no generative model approaches the test set values for foldability and self-consistency, EvoDiff-OADM outperforms EvoDiff-D3PM and improves when increasing the model size (**Fig. 2B-D; Table S3**). We therefore selected the 640M-parameter EvoDiff-OADM model for downstream analysis and hereafter refer to it as EvoDiff-Seq. While a left-to-right autoregressive (LRAR) protein language model generates slightly more structurally-plausible sequences (**Table S3**), EvoDiff-Seq offers the advantage of direct, flexible conditional generation due to its order-agnostic decoding. Unconditional generation from masked language models produces less structurally-plausible sequences because of the mismatch between the training and generation tasks (**Table S3**). Analysis of representative examples of structurally plausible sequences sampled from EvoDiff-Seq across 4 different sequence lengths illustrates their structural plausibility and novelty from sequences in the training set (**Fig. S5**).

To assess the stability and quality of EvoDiff generations *in vitro*, we used EvoDiff-Seq to unconditionally generate a set of 1000 sequences across four different sequence lengths. We then nominated 25 candidates, based on structural plausibility (OmegaFold pLDDT > 70), for experimental characterization (**Supplementary Table 1**). Four designs (EvoDiff-Seq-Designs-1…4, **Fig. 2E**) expressed and had circular dichroism (CD) spectra consistent with mixed alpha helical and beta sheet secondary structure elements (**Fig. 2F-G**). Two of our successful designs (EvoDiff-Seq-Design-3 and EvoDiff-Seq-Design-4) have OmegaFold pLDDT < 85, but many unsuccessful designs have a pLDDT > 85. In one case (EvoDiff-Seq-Design-1), the OmegaFold and AlphaFold predictions even have significant disagreement (> 1Å) in backbone RMSD, a common metric used to filter for designability in structure-based methods. These observations suggest that structural metrics are not always strong predictors of *in vitro* expressibility. Together, these *in silico* and *in vitro* results demonstrate that EvoDiff generates protein sequences that are individually valid.

### Biological properties of generated sequence distributions

Having shown that EvoDiff’s generations are individually foldable and self-consistent, we next evaluated how well the *distribution* of designed protein sequences covered natural protein space. Ideally, generated sequences should capture the natural distribution of sequence, structural, and functional properties while still being diverse from each other and from natural sequences.

Previous work has shown that even without explicit supervision, protein language model embeddings contain information about both sequence and function as captured in GO annotations (*35, 36*). To evaluate coverage over the distribution of sequence and functional properties, we embedded each generated sequence using ProtT5 (*37*), a protein language model explicitly benchmarked for imputing GO annotations (*35*), and calculated the embedding space Fréchet distance between a set of generated sequences and the test set, where lower distance reflects better coverage. We refer to this metric as the Fréchet ProtT5 distance (FPD) and visualize these embeddings and the corresponding FPDs for sequences generated by EvoDiff-Seq and baseline models (**Figs. 3A, S7, S8; Table S1**). For RFdiffusion, we unconditionally generated 1000 structures with the same lengths as for EvoDiff-Seq and then used ESM-IF (*33*) to design their sequences. Both qualitatively and quantitatively, EvoDiff-Seq generates proteins that better recapitulate natural sequence and functional diversity than sampling from a state-of-the-art protein masked language model (ESM-2) or predicting sequences from structures generated by a state-of-the-art structure diffusion model (RFdiffusion) (**Fig. 3A**).

**Figure 3:**
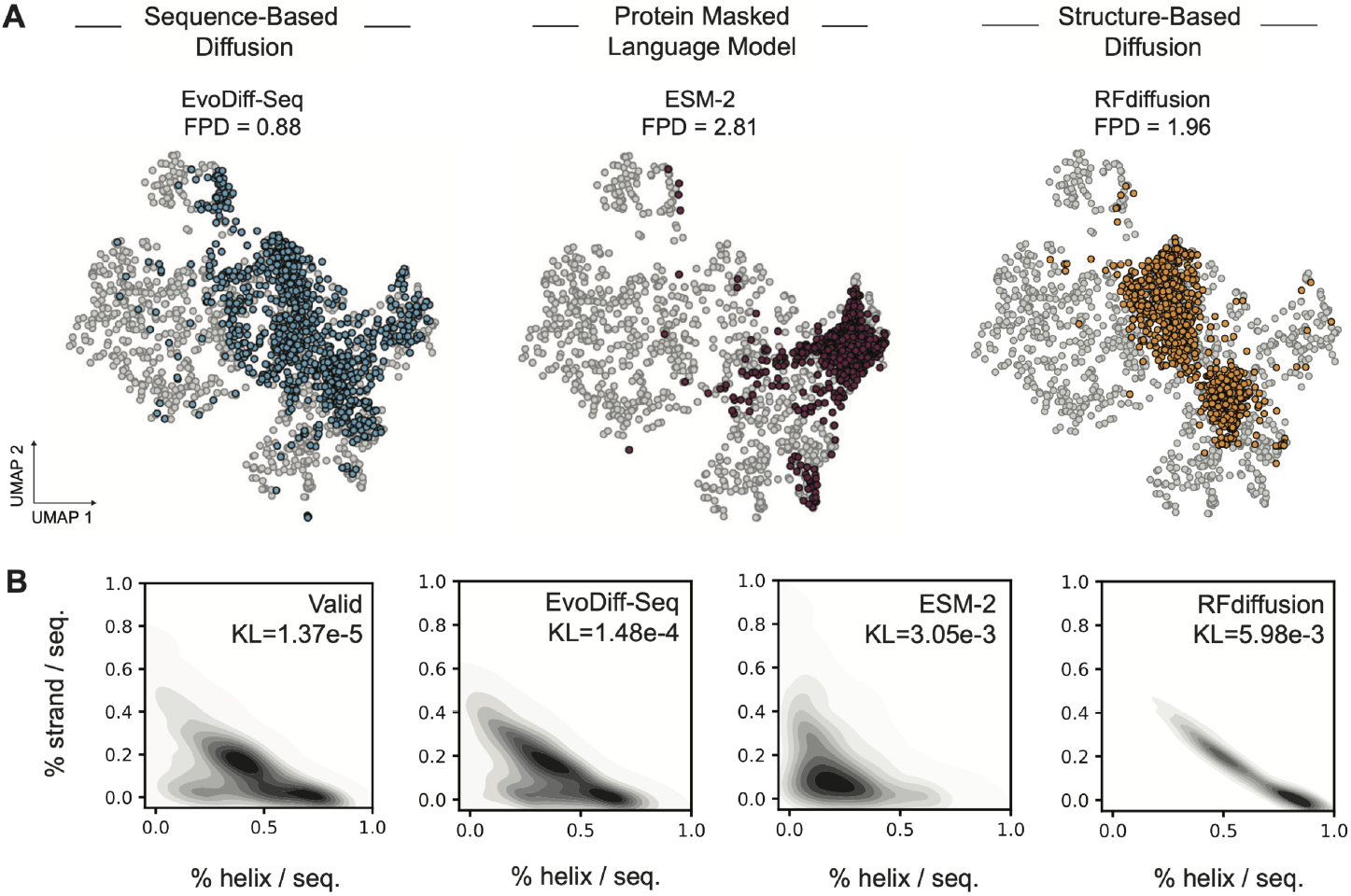
Generated protein sequences capture natural distributions of protein functional and structural features. **(A)** UMAP of ProtT5 embeddings, annotated with FPD, of natural sequences from the test set (grey, *n* =l000) and of generated sequences from EvoDiff-Seq (blue, *n* =l000) and ESM-2 (red, *n* =l000), and inferred sequences inverse-folded from structures from RFdiffusion (orange, *n*=l000). **(B)** Multivariate distributions of helix and strand structural features in generated sequences, based on DSSP 3-state predictions (*n* =l000 samples from each model or the validation set) and annotated with the Kullback-Leibler (KL) divergence relative to the test set.

To evaluate the distribution of structural properties in generated sequences, we computed 3-state secondary structures (*38*) for each residue in generated and natural sequences and quantitatively compared the resulting distributions of structural properties to the distribution for the test set (**Figs. 3B, S9**). EvoDiff-Seq generates proportions of strands and disordered regions that are much more similar to those in natural sequences, while ESM-2 and RFdiffusion both generate proteins enriched in helices (**Fig. 3B**). To ensure our models were not memorizing training data, we calculated the Hamming distance between each generated sequence and all training sequences of the same length and reported the minimum Hamming distance, representing the closest match of any generated sequence to any sequence in the train set (**Table S1**). On average, a sequence generated from EvoDiff-Seq has a Hamming distance of 0.83 from the most similar training distance of the same length. Together, these results demonstrate, via comparison to ESM-2 and RFdiffusion, that EvoDiff’s diffusion objective and evolutionary-scale training data are both necessary to generate novel sequences that cover protein sequence, functional, and structural space.

### Conditional sequence generation for controllable design

EvoDiff’s OADM diffusion framework induces a natural method for conditional sequence generation by fixing some subsequences and inpainting the remainder. Because the model is trained to generate proteins with an arbitrary decoding order, this is easily accomplished by simply masking and decoding the desired portions. We applied EvoDiff’s power for controllable protein design across three scenarios: conditioning on evolutionary information encoded in MSAs, inpainting functional domains, and scaffolding functional structural motifs (**Fig. 1D**).

### Evolution-guided protein generation with EvoDiff-MSA

First, we tested the ability of EvoDiff-MSA to generate query sequences conditioned on the remainder of an MSA, thus generating new members of a protein family without needing to train family-specific generative models. We masked the query sequences from 250 randomly-chosen MSAs from the validation set and newly generated these sequences using EvoDiff-MSA. We then evaluated the quality of the resulting conditionally-generated query sequences via our foldability and self-consistency pipeline (**Fig. 4A**). We find that EvoDiff-MSA generates more foldable and self-consistent sequences than sampling from ESM-MSA (*28*) or using Potts models (*39*) trained on individual MSAs (**Figs. 4B-C, S10; Table S4**). To evaluate sample diversity, we computed the aligned residue-wise sequence similarity between the generated query sequence and the most similar sequence in the original MSA. In contrast to sampling from a Potts model, generating from EvoDiff-MSA yields sequences that exhibit strikingly low similarity to those in the original MSA (**Figs. 4D; Table S4**) while still retaining structural integrity relative to the original query sequences (**Figs. 4E-F; Table S4**) To showcase these properties, we visual ize OmegaFold-predicted structures and evaluation metrics for a sample of high pLDDT, low scPerplexity conditionally-generated query sequences that exhibit low sequence similarity to anything in the conditioning MSA (**Fig. 4G**). These results indicate that EvoDiff-MSA can conditionally generate novel, structurally plausible members of a protein family given guidance from evolutionary information and without further finetuning.

**Figure 4:**
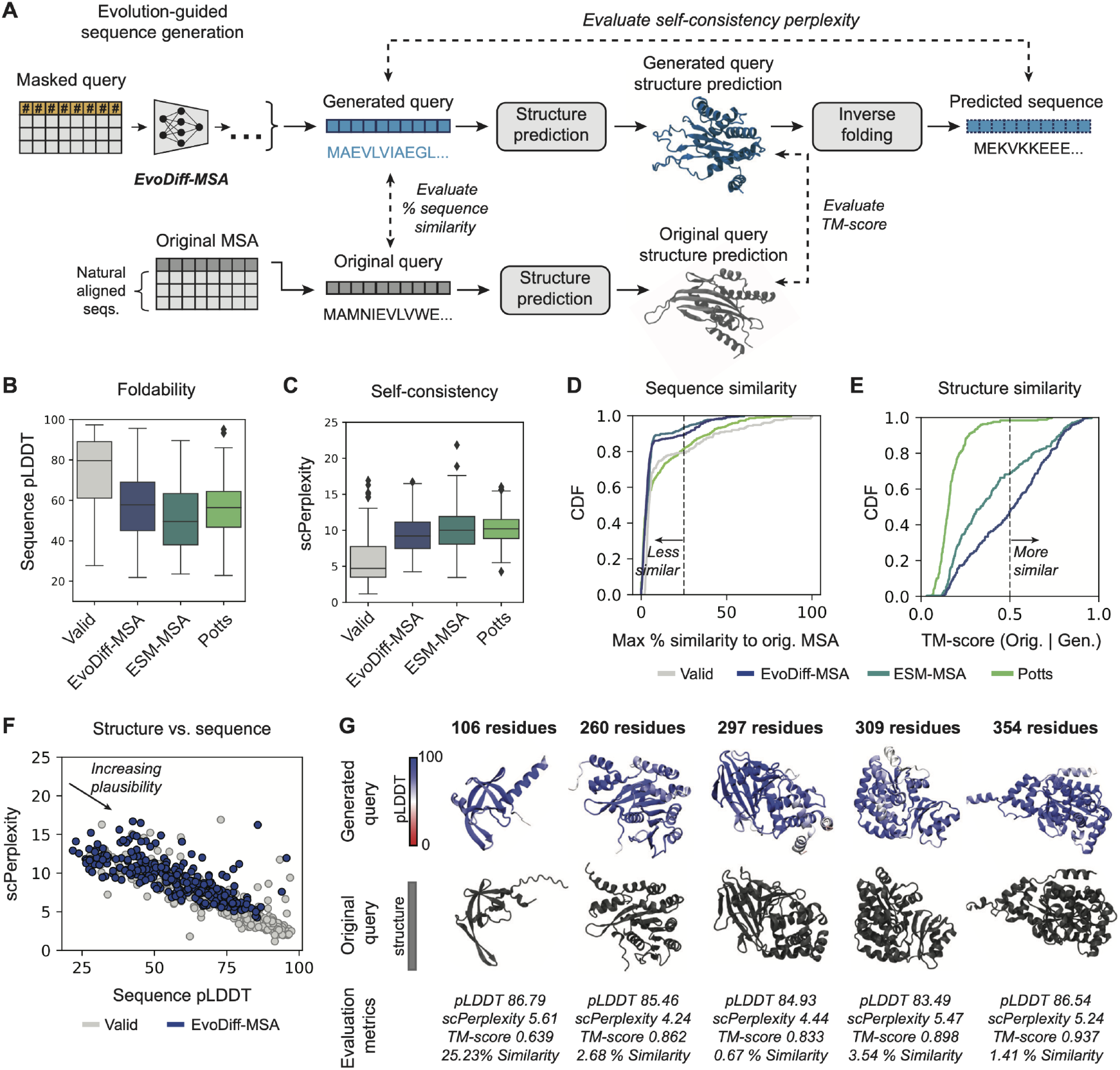
EvoDiff-MSA enables evolution-guided sequence generation. **(A)** A new sequence is generated from EvoDiff-MSA via diffusion over only the query component. Generations are evaluated for diversity and self-consistency and for the quality and consistency of their predicted structures. **(B-E)** Distributions of pLDDT (B), scPerplexity (C), sequence similarity (D; dashed line at 25%), and TM-score (E; dashed line at 0.5) for sequences from the validation set, EvoDiff-MSA, ESM-MSA, and a Potts model (*n=*250 sequences per model; box plots show median and interquartile range). **(F)** Sequence pLDDT versus scPerplexity for sequences from the validation set (grey, *n*=250) and EvoDiff-MSA (blue, *n*=250). **(G)** Predicted structures and metrics for structurally plausible generations from EvoDiff-MSA.

### Generating intrinsically disordered regions

Because it generates directly in sequence space, we hypothesized that EvoDiff could natively generate intrinsically disordered regions (IDRs). IDRs are regions within a protein that lack secondary or tertiary structure; up to 30% of eukaryotic proteins contain at least one IDR, and IDRs make up over 30% of the residues in eukaryotic proteomes (*19*). IDRs carry out important and diverse functional roles in the cell directly facilitated by their lack of structure, such as subcellular trafficking (*40, 41*), protein-protein interactions (*42, 43*), and signaling (*44*). Altered abundance and mutations in IDRs have been implicated in human disease, including neurodegeneration and cancer (*45–47*). Despite their prevalence and critical roles in function and disease, IDRs do not fit neatly in the structure-function paradigm and remain outside the capabilities of structure-based protein design methods.

Having observed that unconditional generation using EvoDiff-Seq produced a similar fraction of residues predicted to lack secondary structure as that in natural sequences (**Fig. 3B**), we used inpainting with EvoDiff-Seq and EvoDiff-MSA to intentionally generate disordered regions via conditioning on their surrounding structured regions (**Fig. 5A**). To accomplish this, we leveraged a previously curated dataset of computationally predicted IDRs covering the human proteome (*48*). We selected this dataset because it also curates orthologs for these proteins, enabling construction of MSAs (*48*). After using EvoDiff to generate putative IDRs via inpainting, we then predicted disorder scores for each residue in the generated and natural sequences using DR-BERT (*49*) (**Figs. 5A, S11**). Over 100 generations, we observe that IDR regions inpainted by EvoDiff-Seq and EvoDiff-MSA result in distributions of disorder scores similar to those for natural sequences, across both the IDR and the surrounding structured regions (**Figs. 5B, S12**). Generations from EvoDiff-MSA exhibit strong correlation in predicted disorder scores to those of true IDRs (**Fig. S12**). Although putative IDRs generated by EvoDiff-Seq are less similar to their original IDR than those from EvoDiff-MSA (**Fig. 5C**), both models generated disordered regions that preserve disorder scores over the entire protein sequence and still exhibit low sequence similarity to the original IDR (**Fig. S13**).

**Figure 5:**
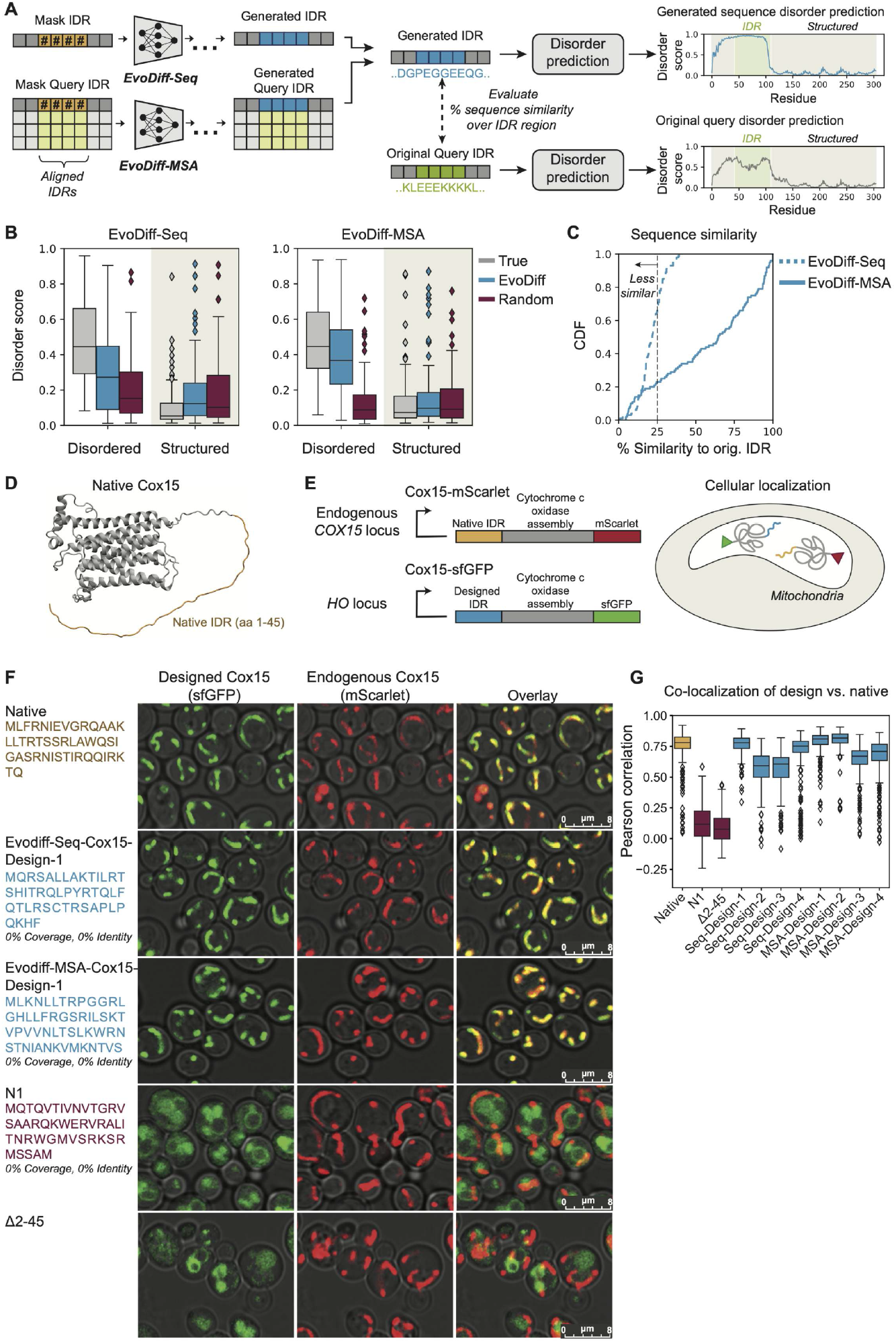
EvoDiff generates intrinsically disordered regions. **(A)** A new IDR sequence is generated from EvoDiff-Seq or EvoDiff-MSA by inpainting disordered residues in the query sequence. DR-BERT is then used to predict disorder scores for the original and regenerated sequences. **(B)** Distributions of disorder scores over disordered and structured regions for sequences with true (grey), inpainted (blue), and randomly-sampled (red) IDRs (*n*=100 sequences per condition; box plots show median and interquartile range). **(C)** Distribution of sequence similarity relative to the original IDR for generated IDRs from EvoDiff-Seq (blue, dashed) and EvoDiff-MSA (blue, solid) (*n*=100; dashed line at 25%). **(D)** Structure of the yeast Cox15 protein, with the disordered N-terminal mitochondrial targeting signal highlighted in yellow. EvoDiff is used to inpaint this disordered region. **(E)** Microscopy assay in yeast to assess cellular localization of Cox15 proteins with EvoDiff-designed targeting signals. Mitochondrial localization is measured by quantifying the fluorescence overlap between endogenous Cox15 (mScarlet) and designed Cox15 (sfGFP). **(F)** Fluorescence microscopy imaging of yeast cells. Columns from left to right show: the N-terminal IDR sequence used for a given Cox15-sfGFP design (rows); signal from the designed Cox15 (sfGFP, colored in green); signal from the endogenous Cox15 (mScarlet, colored in red); overlaid images. All images show brightfield in grey, to visualize cell boundaries. Images for all 8 designs are included in **Fig. S15** and **S16**. Scale bars = 8 *μ*m. **(G)** Per-cell pixel-wise Pearson correlation coefficients of signal intensities between the mScarlet channel and the sfGFP channel over *n*=183-313 cells segmented by YeastSpotter for each design. Correlations for a biological replicate are shown in **Fig. S17**.

We next sought to validate experimentally that inpainted IDRs maintain cellular function. Many IDRs are targeting signals that drive precise subcellular localization critical to the protein’s underlying function. Notably, the vast majority of mitochondrial proteins contain N-terminal disordered mitochondrial targeting signals that are cleaved after import into the mitochondria (*40, 41, 50–52*). We hypothesized that, given the mature protein as context, EvoDiff can design N-terminal IDRs that drive mitochondrial localization in cells. To test this hypothesis, we masked out the N-terminal IDR (amino acid positions 1-45, inclusive) of the yeast Heme A synthase Cox15 and inpainted 200 IDR candidates of varying lengths from EvoDiff-Seq and EvoDiff-MSA (Rand subsampling) (**Fig. 5D**). We predicted disorder with DR-BERT and mitochondrial targeting propensity with MitoFates (*53*) for each candidate (**Fig. S14A-B**) and selected the top eight generated sequences (**Fig. S14C-D**), four from each of EvoDiff-Seq and EvoDiff-MSA, for experimental characterization (**Supplementary Table 1**).

We replaced the native Cox15 N-terminal IDR with each of the generated IDRs and assessed the cellular localization of the resultant proteins using a microscopy screen in yeast cells. Briefly, we tagged the generated constructs with super-folding GFP (sfGFP) and recombined them into a yeast strain expressing endogenous Cox15 tagged with mScarlet, such that overlap between sfGFP (green) and mScarlet (red) fluorescence would indicate co-localization of the designed and endogenous Cox15 and thus demonstrate that the generated IDRs successfully induce mitochondrial transport (**Fig. 5E**). As negative controls, we used both a knockout of the native Cox15 N-terminal IDR (∆2-45) as well as the N1 N-terminal sequence, which was designed in a previous IDR study to match the bulk biophysical properties of the Cox15 mitochondrial targeting signal but lacks mitochondrial targeting activity, as predicted by MitoFates and demonstrated experimentally (*54*). Consistent with our hypothesis, all 8 EvoDiff-IDR-Cox15 variants exhibit a subcellular localization pattern consistent with the localization of the native Cox15, suggesting that EvoDiff can design IDRs that function in their cellular context (**Fig. 5F-G, S15-S17**).

### Scaffolding functional motifs with sequence information alone

Thus far, the primary application of deep generative models of protein structure in protein engineering is their ability to scaffold binding and catalytic motifs: given the 3D coordinates of a functional motif, these models can often generate a structural scaffold that holds the motif in precisely the 3D geometry needed for function (*10, 14, 55*). Given that the fixed functional motif includes the residue identities for the motif, we investigated whether a structural model is actually necessary for motif scaffolding.

We used conditional generation with EvoDiff to generate scaffolds for a diverse set of 17 motif-scaffolding problems (*10*) by fixing the functional motif, supplying only the motif’s amino-acid sequence as conditioning information, and then decoding the remainder of the sequence (**Fig. 1D**). The problems include simple “inpainting”, viral epitopes, receptor traps, small molecule binding sites, protein-binding interfaces, and enzyme active sites. Many of the motifs are not contiguous in sequence space. We compared the performance of EvoDiff, which uses only sequence information, to the state-of-the-art structure model RFdiffusion, and facilitated direct comparisons by using OmegaFold to predict structures for our generated sequences as well as for sequences inverse-folded from RFdiffusion structures. Notably, we use the same EvoDiff models for both unconditional and conditional generation, while the version of RFdiffusion used for scaffolding is finetuned from that used for unconditional generation.

We evaluated the ability of each of EvoDiff-Seq, EvoDiff-MSA, and RFdiffusion to generate successful scaffolds (**Fig. 6A-B**), where we define a scaffold to be successful if the predicted motif coordinates have less than 1Å RMSD from the desired motif coordinates and if the scaffolded sequence has a pLDDT > 70. Despite operating entirely in sequence space, EvoDiff-Seq and EvoDiff-MSA generate successful scaffolds for 8 and 13 of the 17 problems, respectively (**Table S5, S6**). EvoDiff-MSA has a higher success rate than EvoDiff-Seq for 10 problems and a higher success rate than RFdiffusion for 6 problems. EvoDiff-Seq has a higher success rate than RFdiffusion for 2 problems and a higher success rate than EvoDiff-MSA for 3 problems. There are two scaffolding problems (1YCR, 3IXT) where EvoDiff-MSA is outperformed by both EvoDiff-Seq and RFdiffusion (**Table S5, S6**). These are both examples where, for scaffolding, an MSA containing fewer than 64 protein sequences was input to EvoDiff-MSA, which did not see MSAs with fewer than 64 sequences during training.

**Figure 6:**
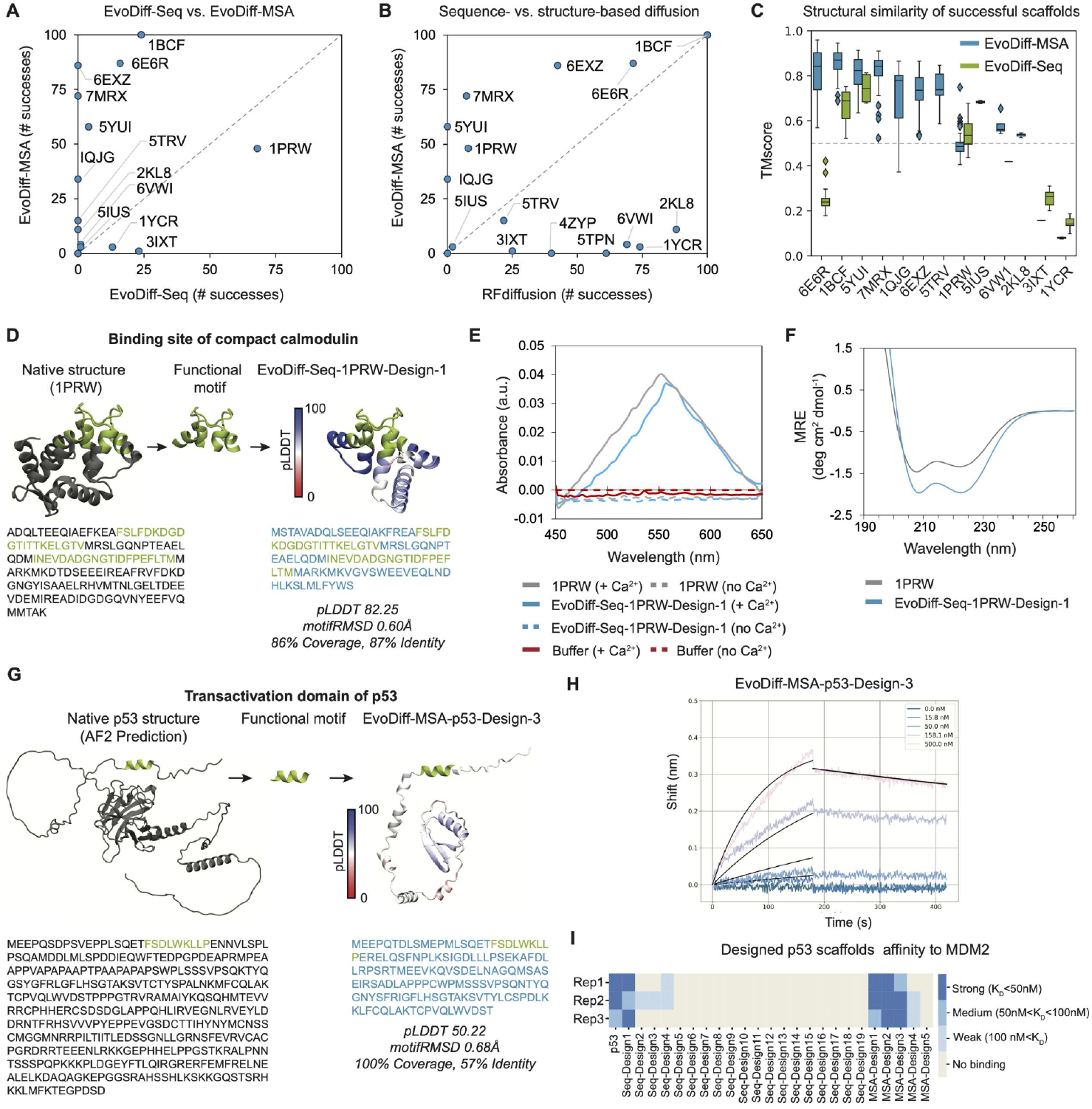
EvoDiff scaffolds functional motifs without explicit structural information. **(A)** Number and identity of successful scaffolds from *n*=100 trials for EvoDiff-Seq (x-axis) versus EvoDiff-MSA (y-axis) across scaffolding problems in which at least one method succeeds. **(B)** Performance comparison of sequence-based scaffolding via EvoDiff-MSA (y-axis) versus structure-based scaffolding via RFdiffusion (x-axis) across scaffolding problems in which at least one method succeeds (*n*=100 trials per problem). **(C)** Distributions of TM-scores of successfully generated scaffolds from EvoDiff models relative to the true structures (dashed line at 0.5; box plots show median and interquartile range). **(D)** EvoDiff-Seq successfully scaffolds the binding site of compact calmodulin. The generated sequence, OmegaFold-predicted structure (colored by pLDDT), and computed metrics for EvoDiff-Seq-1PRW-Design1 are shown. Scaffolding motif is shown in green, and the structure and sequence of calmodulin (1PRW) are shown in dark grey. **(E)** Background-corrected absorbance spectra of the native 1PRW (grey), EvoDiff-Seq-1PRW-Design-1 (blue), and control buffers (red) with (solid line) and without (dashed lines) Ca^2+^. **(F)** CD spectra of native 1PRW (grey) and EvoDiff-Seq-1PRW-Design-1 (blue). **(G)** EvoDiff successfully scaffolds the transactivation domain of p53. Generated sequence, OmegaFold-predicted structure (colored by pLDDT), and computed metrics for EvoDiff-MSA-p53-Design-3, are shown. The transactivation domain is shown in green, and the AlphaFold-predicted structure and sequence of p53 are shown in dark grey. **(H**) BLI measurements for EvoDiff-MSA-p53-Design-3 measured over 5 concentrations. **(I)** *In vitro* binding affinities of all EvoDiff designs against the full-length human MDM2. Measurements for all designs were performed in triplicate (**Fig. S20-S24**).

Interestingly, there is almost no correlation between the problem-specific success rates of EvoDiff and RFdiffusion, and there are very few problems for which both methods have high success rates, showing that EvoDiff may have orthogonal strengths to RFdiffusion (**Fig. 6A-B**). Due to its conditioning on evolutionary information, EvoDiff-MSA generates scaffolds that are more structurally similar to the native scaffold than EvoDiff-Seq (**Fig. 6C**). To ensure that EvoDiff is not finding trivial solutions, we show that it outperforms both random generation and the single-order LRAR model (which decodes unconditionally up to and after a motif) (**Table S5**). ESM-MSA performs similarly to EvoDiff-MSA on this task, as the motif scaffolding task is well-aligned with its training task, and it is trained on approximately 200x more MSAs than EvoDiff-MSA (**Table S6**). We illustrate examples of successful scaffolds sampled from EvoDiff and note both the qualitative and quantitative quality of generated proteins and predicted structures across a range of functional motifs (**Fig. S18**).

To validate EvoDiff’s ability to design functional scaffolds around structural motifs, we generated scaffolds for two distinct targets – calcium (Ca^2+^) and MDM2 – by conditioning on the binding motif sequences from calmodulin (1PRW) and p53 (1YCR), respectively. We do this without providing any explicit information about the binding target. First, we generated 100 scaffolds each from EvoDiff-Seq, EvoDiff-MSA (Max subsampling), and EvoDiff-MSA (Rand subsampling) for the discontinuous 1PRW binding domain (**Fig. 6D**). Then, we nominated 13 designs, based on *in-silico* metrics (pLDDT > 70 and motifRMSD < 1Å), to express and evaluate experimentally (**Supplementary Table 1**). 12 of these designs expressed successfully, and one of these designs (EvoDiff-Seq-1PRW-Design-1) demonstrated a spectroscopic shift in the presence of Ca^2+^ (**Fig. 6F, S19**), indicative of calcium binding (*56*). We show the predicted structure of the successful binder in **Fig. 6E**. Binding experiments were repeated independently in triplicate (**Fig. S19**).

Next, we scaffolded the transactivation domain of p53 (**Fig. 6G**), which binds MDM2. Given the importance of p53 as a tumor suppressor in cancer, competitive inhibition of the p53-MDM2 interaction via a synthetic scaffold could promote tumor-suppressing activity of native p53 (*57*). Unlike in structure-based methods (*10*), we did not include information about the binder during inference; additionally, we note that p53 contains an intrinsically disordered N-terminal domain (*58*) (**Fig. 6G**). Starting with 900 EvoDiff-Seq and 400 EvoDiff-MSA generations, we nominated 24 total designs (**Supplementary Table 1**). For EvoDiff-Seq we nominated 19 designs with OmegaFold pLDDT > 70 and motifRMSD < 1Å. However, no EvoDiff-MSA generations exceeded the pLDDT threshold, so we nominated 5 candidates from EvoDiff-MSA using only motifRMSD. We screened all 24 designs *in vitro* for binding to full-length human MDM2 using biolayer interferometry (BLI) (**Fig. S20-S24**). All designs expressed in a cell-free system and four designs (EvoDiff-Seq-p53-Design1 and Evodiff-MSA-p53-Design-1…3) demonstrated strong binding to full-length MDM2, categorized as *K*_*D*_ < 50 nM observed over at least two replicates. The strongest binders, EvoDiff-MSA-p53-Design-2 and EvoDiff-MSA-p53-Design-3, bind with an average *K*_*D*_ of 25.9 nM and 40.2 nM while having 85% and 53% sequence similarity, respectively, to any natural protein sequence (**Fig. 6H, S22; Supplementary Table 1**). In total, 8 designs demonstrated evidence of binding at varying strengths, as measured across three replicates (**Fig. 6I, S21, S22**). These *in silico* and *in vitro* results demonstrate that EvoDiff can design functional scaffolds around structural motifs via conditional generation in sequence space alone.

## Discussion

We present EvoDiff, a diffusion modeling framework capable of generating high-fidelity, diverse, and novel proteins with the option of conditioning according to sequence constraints.

We validate EvoDiff’s generative capabilities both *in silico* and in the lab. Because it operates in the universal protein design space, EvoDiff can unconditionally sample diverse structurally-plausible proteins that express and exhibit stable secondary structure, generate functional intrinsically disordered regions, and scaffold structural motifs using only sequence information, challenging a paradigm in structure-based protein design.

EvoDiff is the first suite of deep learning models to demonstrate the power of diffusion generative modeling over evolutionary-scale protein sequence space. Unlike previous attempts to train diffusion models on protein structures (*7–15*) and/or sequences (*59–63*), EvoDiff is trained on a large, diverse sample of all natural sequences, rather than on smaller protein structure datasets or sequence data from a specific protein family. Previous protein generative models trained on global sequence space have been either left-to-right autoregressive (LRAR) models (*64–67*) or masked language models (MLMs) (*27, 28, 30, 37, 68*). EvoDiff’s OADM training task generalizes the LRAR and MLM training tasks. Specifically, the OADM setup generalizes LRAR by considering all possible decoding orders, while the MLM training task is equivalent to training on one step of the OADM diffusion process.

This generalized mathematical formulation yields empirical benefits, as EvoDiff-Seq produces sequences that better cover protein functional and structural space than sampling from state-of-the-art protein MLMs (**Fig. 3**). While an LRAR model learned to fit the evolutionary sequence distribution better (**Table S1**), the fixed decoding order of traditional left-to-right autoregression restricts conditional generation. EvoDiff directly addresses this barrier by enabling different forms of conditioning, including evolution-guided generation (**Fig. 4**) as well as inpainting and scaffolding (**Figs. 5-6**). We report the first demonstrations of these programmable generation capabilities from deep generative models of protein sequence alone.

Future work may expand these capabilities to enable conditioning via guidance, in which generated sequences can be iteratively refined to fit desired properties (*69, 70*). While we observe that OADM generally outperforms D3PM in unconditional generation, likely because the OADM denoising task is easier to learn than that of D3PM, conditioning via guidance intuitively fits into the EvoDiff-D3PM framework because the identity of each residue in a sequence can be edited at every decoding step. OADM and existing conditional LRAR models, such as ProGen (*64*), both fix the identity of each amino acid once it is decoded, limiting the effectiveness of guidance. Guidance-based conditioning of EvoDiff-D3PM should enable the generation of new protein sequences specifying functional objectives, such as those specified by sequence-function classifiers.

Because EvoDiff only requires sequence data, it can readily be extended for diverse down-stream applications, including those not reachable from a traditional structure-based paradigm. As a first example, we have demonstrated EvoDiff’s ability to generate functional IDRs – overcoming a prototypical failure mode of structure-based predictive and generative models – via inpainting without fine-tuning. Fine-tuning EvoDiff on application-specific datasets, such as those from display libraries or large-scale screens, may unlock new biological, therapeutic, or scientific design opportunities that would be otherwise inaccessible due to the cost of obtaining structures for large sequence datasets. Experimental data for structures is much sparser compared to sequences, and while structures for many sequences can be predicted using AlphaFold and similar algorithms, these methods do not work well on point mutants and can be overconfident on spurious proteins (*71, 72*).

While we demonstrated some coarse-grained strategies for conditioning generation through scaffolding and inpainting, to achieve even more fine-grained control over protein function, with future development EvoDiff may be conditioned on text, chemical information, or other modalities. For example, text-based conditioning (*73*) could be used to ensure that generated proteins are soluble, readily expressed, and non-immunogenic. Future use cases for this vision of controllable protein sequence design include programmable modulation of nucleic acids via conditionally-designed transcription factors or endonucleases, improved therapeutic windows via biologics optimized for *in vivo* delivery and trafficking, as well as newly-enabled catalysis via zero-shot tuning of enzyme substrate specificity.

In summary, we present an open-source and experimentally-validated suite of discrete diffusion models that provide a foundation for sequence-based protein engineering and design. EvoDiff models can be directly deployed for unconditional, evolution-guided, and conditional generation of protein sequences and may be extended for guided design based on structure or function. We envision that EvoDiff will enable new abilities in controllable protein design by reading and writing function directly in the language of proteins.

## Acknowledgements

The authors thank Christian Dallago for helpful discussions about the methods, applications, and evaluations; Jonathan Carlson for valuable feedback on the manuscript; Kevin E. Wu for providing sequences generated from FoldingDiff; Joseph Watson and David Juergens for providing sequences generated from RFDiffusion; and Alessandro Sordoni, Hannes Schultz, Rémi Piché-Taillefer, Remi Tachet des Combes, and Sean Whitzell for assistance on using Microsoft’s compute resources ; Philip Rosenfield for his immense effort in managing the project; Colby T. Ford for providing valuable input on the GitHub repository; and Hannah Wayment-Steele for identifying disordered protein annotations.

## Author contributions

Conceptualization: S.A., N.T., N.F., A.P.A., K.K.Y.; Methodology: S.A., N.T., R.vdB., A.P.A., K.K.Y.; Software Programming: S.A., N.T., K.K.Y.; Experimental Design: S.A., R.S., A.M.M, A.X.L., A.P.A., K.K.Y.; Investigation: S.A., N.T., R.S., A.X.L., A.P.A., K.K.Y.; Validation: S.A., N.T., R.S., A.X.L., A.P.A., K.K.Y.; Formal analysis: S.A., N.T., A.X.L., A.P.A., K.K.Y.; Resources Provision: N.F., K.K.Y.; Data Curation: S.A., N.T., A.X.L., K.K.Y.; Visualization: S.A., R.S., A.P.A., K.K.Y; Writing - Original Draft: S.A., A.P.A., K.K.Y.; Writing - Review & Editing: S.A., N.T., R.vdB., N.Ten., R.S., A.M.M., A.X.L., N.F., A.P.A., K.K.Y.; Supervision: A.M.M., N.F., A.P.A., K.K.Y.

## Resource availability

Code is available at https://github.com/microsoft/evodiff. Model weights, generated sequences, and computed metrics are available at https://zenodo.org/record/8332830.

## Methods

### Diffusion models

Diffusion models are a class of generative models that learn to generate data from noise. They consist of a forward corruption process and a learned reverse denoising process. The forward process is a Markov chain of diffusion steps *q*(*x*_*t*_|*x*_*t*−1_) that corrupts an input (*x*_0_) over *T* timesteps such that *x*_*T*_ is indistinguishable from random noise. The learned reverse denoising process *p*_*θ*_(*x*_*t*−1_|*x*_*t*_) is parameterized by a model such as a neural network and generates new data from noise. Discrete diffusion models have previously been developed over binary random variables (*3*), developed over categorical random variables with uniform transition matrices (*74, 75*), linked to autoregressive models (*24*), and optimized for use with transition matrices (*25*).

This work presents models from two different discrete diffusion frameworks – order-agnostic autoregressive diffusion models (OADMs) and discrete denoising diffusion probabilistic models (D3PMs) – on protein sequences and multiple sequence alignments (MSAs).

### Discrete Denoising Diffusion Probabilistic Models (D3PMs)

Discrete denoising diffusion probabilistic models (D3PMs) operate by defining a transition matrix *Q* such that, over *T* timesteps, discrete inputs (i.e. protein amino-acid sequences for EvoDiff) are iteratively corrupted via a controlled Markov process until they constitute samples from a uniform stationary distribution at time *T*. This section describes the D3PM process and loss for a single categorical variable *x* in one-hot format. The forward corruption process is described by:

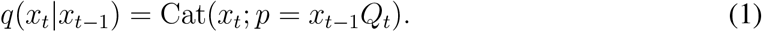

This allows for efficient training via efficient computation of *q*(*x*_*t*_|*x*_0_) and *q*(*x*_*t*−1_|*x*_*t*_). The D3PM approach can emulate a masked modeling process by choosing a transition matrix with an absorbing state (e.g., [`MASK`]; (*25*)). However, in this work, the D3PM formulation is only used for discrete corruption because masking corruption via OADM generally outperforms absorbing-state D3PM (*24*). EvoDiff includes two discrete corruption schemes: one based on a uniform transition matrix (D3PM-Uniform) and one based on a biologically-informed transition matrix (D3PM-BLOSUM).

EvoDiff-D3PM models are trained via a hybrid loss function

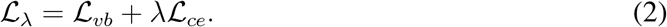

This loss combines a variational lower bound ℒ_*vb*_ on the negative log likelihood

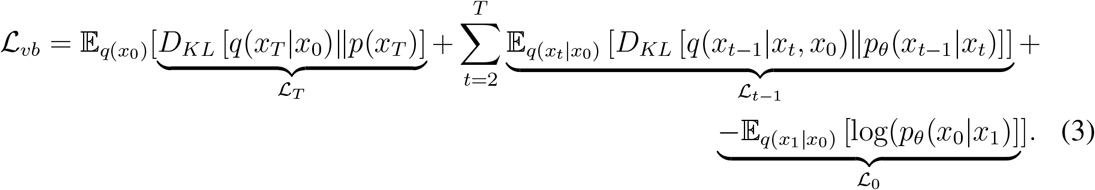

and a cross-entropy loss ℒ_*ce*_ on *p*_*θ*_(*x*_0_|*x*_*t*_). Investigation of the impact of *λ* on model performance revealed minimal improvement to sample generation quality when *λ* > 0, consistent with the findings of the original D3PM paper (*25*). Thus *λ*=0 and *T* =500 were used in all D3PM experiments.

ℒ_*vb*_ has three terms. ℒ_*T*_ measures whether the corruption reaches the stationary distribution *p*(*x*_*T*_) at time *T* and does not depend on *θ*. Next consider the remaining two terms ℒ_*t*−1_ and ℒ_0_, which depend on *θ*. Following the original D3PM paper, 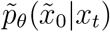 is directly predicted by the neural network. To compute the loss at timesteps 0 < *t* < *T*, the terms *q*(*x*_*t*−1_|*x*_*t*_, *x*_0_) and *p*_*θ*_(*x*_*t*−1_|*x*_*t*_) must be computed from *x*_*t*_, *x*_0_, and 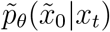 using Markov properties

Defining 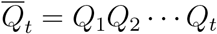:

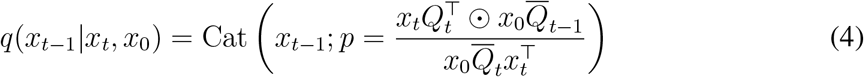

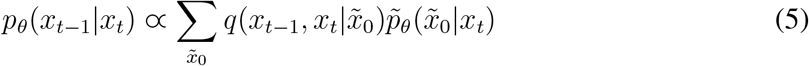

where ⊙ represents an element-wise product. For Equation 5 rules of conditional probability and Markov properties are used to define 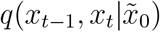 in terms of *x*_*t*_ and 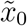:

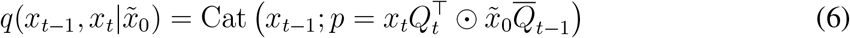

Putting everything together, at each step of training a corruption timestep is sampled according to *t* ∼ 𝒰 (1, …, *T* − 1). *x*_*t*_ is then sampled via 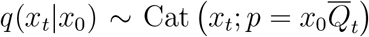 for every residue in the input protein, and the neural network predicts 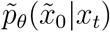. Note that, while the corruption and loss are computed independently over each residue, the neural network predicts 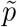 in the context of the entire sequence. If *t* = 1, only the loss ℒ_0_ is used, reflecting a standard negative log likelihood. Otherwise, Equations 4 and 5 are used to compute the loss ℒ_*t*−1_.

Sampling from a trained model begins with the noised x_*T*_, where each residue is randomly sampled from a uniform distribution over amino acids. *x*_*t*−1_ is then iteratively sampled via *p*_*θ*_(*x*_*t*−1_|*x*_*t*_) as described in Equation 5. For all models, generated sequences are sampled to match the distribution of sequence lengths in the training set, going up to 2048 residues as the maximum length.

### EvoDiff-D3PM-Uniform

Many strategies exists to schedule corruption in D3PMs. EvoDiff-D3PM-Uniform employs the simplest case – a uniform corruption scheme. Specifically, EvoDiff-D3PM-Uniform models implement a doubly stochastic, uniform transition matrix *Q*_*t*_ with a corruption schedule (*T* − *t* + 1)^−1^ from Sohl-Dickstein et al. (*3*), so that information is linearly corrupted between *x*_*t*_ and *x*_0_ for all *t* < *T*.

### EvoDiff-D3PM-BLOSUM

EvoDiff-D3PM-BLOSUM implements a transition matrix derived from BLOSUM62 matrices of amino acid substitution frequencies (*76*). BLOSUM matrices are derived from observed alignments across highly conserved regions of protein families and thus provide the relative frequencies of amino acids and their substitution probabilities.

Rows that represent uniform transition probabilities for non-standard amino acid codes (J, O, U) and for < `GAP` > tokens in the MSA input case are included in addition to standard amino acids. BLOSUM substitution frequencies are converted to a matrix of transition probabilities by performing a softmax over the frequencies and then normalizing over rows and columns via the Sinkhorn-Knopp algorithm to obtain a doubly stochastic matrix. In this scheme, the gradual corruption of a single sequence to random noise is simulated in a way that prioritizes conserved evolutionary relationships of amino acid mutations. A *β*-schedule was implemented to taper the number of mutations over time for timesteps up to *T* =500, specifically via an empirical schedule that corrupts half the sequence content by half of *T* (*t*=250) (Fig. S25). This schedule was chosen to approximate the linear rate of mutations observed over 500 timesteps in the uniform transition matrix case, shown in Fig. S25b.

### Order-Agnostic Autoregressive Diffusion Models (OADMs)

Order-agnostic autoregressive diffusion models (OADMs) generalize absorbing-state D3PM and left-to-right autoregressive models (LRARs) (*24*). This section describes the OADM process and loss for a sequence *x* of *L* categorical variables. In the case of EvoDiff, *L* is the sequence length.

LRARs factorize a high-dimensional joint distribution *p*(*x*) into the product of *L* univariate distributions using the probability chain rule:

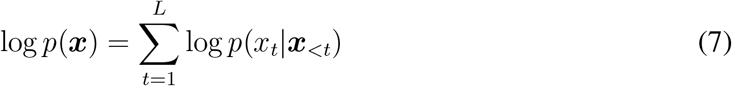

where *x*_<*t*_ = *x*_1_, *x*_2_, … *x*_*t*−1_. LRARs are typically parametrized using a triangular dependency structure, such as causal masking in a transformer or CNN, in order to allow parallelized computation of all the conditional distributions in the likelihood during training. LRARs learn to generate sequences in a pre-specified left-to-right decoding order, which may be non-obvious for modalities such as proteins and does not allow conditioning on arbitrary fixed subsequences.

LRARs can be expanded into a diffusion framework via two subtle changes. Following the exposition in Hoogeboom *et al*., (*24*), the first change is to allow order-agnostic decoding. In an order-agnostic autoregressive model, a decoding order *σ* is first sampled uniformly from all possible decoding orders *S*_*L*_. At time step *t* in the forward process, *x*_*σ*(*L*−*t*)_ is masked. The log-likelihood for an order-agnostic autoregressive model is derived using Jensen’s inequality:

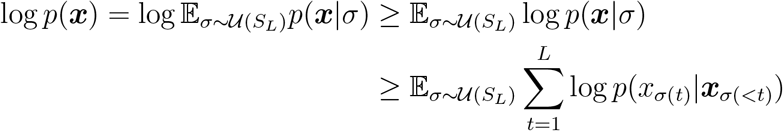

The next change involves an objective that optimizes over arbitrary decoding orders one timestep at a time in the style of modern diffusion models, without requiring a neural network that enforces a triangular or causal dependency structure. This is accomplished by replacing the summation over *t* by an expectation that is appropriately re-weighted.

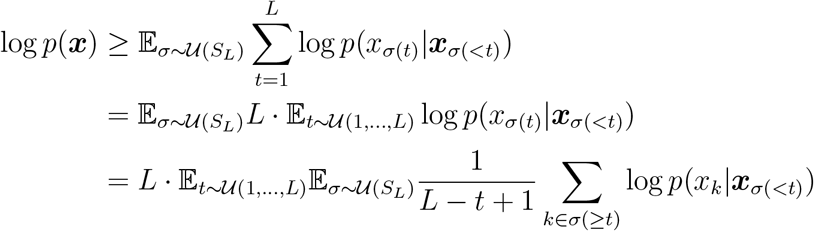

The overall expected log likelihood log *p*(*x*) can be thought of according to a series of likelihoods, each captured in the loss at step *t*, ℒ_*t*_:

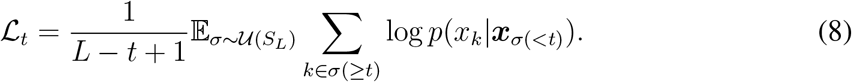

Thus, the overall expected log likelihood is lower bounded as:

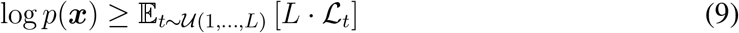

A neural network can be efficiently trained to learn the reverse process *p*_*θ*_(*x*_*σ*(*t*)_|*x*_*σ*(<*t*)_) by randomly masking a set of *t* tokens at each iteration and minimizing the reweighted loss, allowing the model to learn from predictions of all masked positions at each timestep. By learning one model over all possible decoding orders, OADM allows for conditioning by fixing arbitrary subsequences at generation time. Sequences were generated unconditionally from OADM models by beginning with an all-mask sequence as input, randomly sampling a decoding order, and sampling each token from the predicted probability distribution.

### Left-to-right autoregressive and masked language models are diffusion models

The connection between autoregressive models and diffusion models has been described previously (*24, 25*). Left-to-right autoregressive (LRAR) diffusion models implement a masked modeling process that is akin to a process which iteratively and deterministically masks all tokens to the right of the sampled token *x*_*t*_, where the current diffusion timestep *t* is equivalent to the number of tokens masked over the entire sequence length, with all tokens masked at the final timestep *T* = *L*.

Likewise, masked language models (MLMs) are equivalent to only learning one step *t* of OADM:

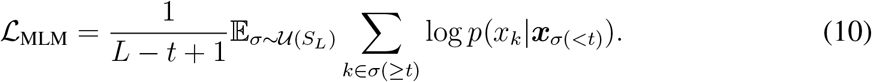

Thus, the OADM setup generalizes LRAR models by considering all possible decoding orders rather than left-to-right decoding, while the MLM learning task is equivalent to only training on one step of the OADM diffusion process.

### Datasets

Sequence-only EvoDiff models were trained on UniRef50 (*26*) which contains approximately 45 million protein sequences. The UniRef50 release and train/validation/testing splits from CARP (*27*) were used to facilitate comparisons between models. Sequences longer than 1024 residues were randomly subsampled to 1024 residues. This split includes the random assignment of 82,929 sequences to the validation set and 209,426 sequences to the test set. Multiple sequence alignment (MSA) EvoDiff models were trained on OpenFold (*29*), which contains 401,381 MSAs for 140,000 unique Protein Data Bank (PDB) chains and 16,000,000 UniClust30 clusters. To construct the MSAs used to train EvoDiff, insertions, denoted by lowercase characters, were removed to restore the alignments, as the queries do not contain gap characters. Next, MSAs that contained sequences with more than 512 consecutive < `GAP` > tokens as well as MSAs that contained fewer than 64 sequences per alignment were filtered out. This filtering resulted in 382,296 total MSAs, which were then randomly split into 372,296 training and 10,000 validation MSAs.

### MSA subsampling for training EvoDiff-MSA models

To optimize for memory constraints during training, MSAs were subsampled to 64 sequences and a maximum sequence length of 512. MSAs shorter than 512 sequences were padded to a sequence length of 512, but MSAs containing fewer than 64 sequences were excluded from training. For MSAs with more than 64 sequences, two subsampling schemes were implemented: random (“Rand.”) and MaxHamming (“Max”). The random subsampling scheme (“Rand.”) randomly samples 64 sequences from the MSA, making sure that the reference/query sequence (i.e. the first sequence) is always included. The “Max” subsampling scheme greedily selects for sequence diversity in the 64 sequence subset by iteratively selecting the sequence that maximizes the minimum Hamming distance to the sequences already selected. The Hamming distance measures the distance between two sequences, denoted by the number of amino acids that differ between aligned sequences. Subsampling to maximize the Hamming distance enabled input of an MSA rich with evolutionarily diverse sequences to EvoDiff-MSA models.

### Modeling, architecture, and training details

For sequences, the EvoDiff denoising model adopts a ByteNet-style CNN architecture (*77*) previously shown to perform similarly to transformers for protein sequence masked language modeling tasks (*27*). All models are implemented in PyTorch (*78*). In EvoDiff-OADM models, the diffusion timestep is implicitly encoded in the number of masked positions. EvoDiff-D3PM models use a 1D sinusoidal encoding (*79*) to denote the timestep for each input. All sequence models were trained with the Adam optimizer (*80*), a learning rate of 1e-4 with linear warmup over 16,000 steps, and dynamic batching that adaptively selects the number of samples to maximize GPU utilization. EvoDiff’s small sequence models implement a ByteNet-style architecture with ca. 38M parameters. Large models were scaled to a ByteNet architecture of ca. 640M parameters by increasing the model dimension *d* from 1020 to 1280, increasing the encoder hidden dimension from *d/*2 to *d*, and increasing the number of layers from 16 to 56.

38M parameter models were trained on 8 32GB NVIDIA V100 GPUs; 640M parameter models were trained on 32 (2×16) 32GB NVIDIA V100 GPUs. The maximum number of tokens per GPU in each batch was reduced from 40,000 to 6,000 to accommodate training the larger 640M parameter models. 38M parameter models were trained for approximately 2 weeks and saw ca. 3e14 tokens over 700,000 training steps. 640M parameter models were trained to the extent permitted by available compute resources to maximize model performance. Models saw between ca. 1e10 and 1e17 tokens over ca. 400,000-2,000,000 training steps. The D3PM-BLOSUM model ceased to improve after approximately 12 days of training. The D3PM-Uniform and OADM models were trained for 23 days without reaching convergence.

For MSAs, the EvoDiff denoising model adopts a 100M parameter MSA Transformer architecture (*28*). As with the single sequence models, EvoDiff-MSA-OADM models implicitly encode the diffusion timestep; EvoDiff-MSA-D3PM models include an additional sinusoidal timestep embedding. All MSA models were trained with the Adam optimizer with a learning rate of 1e-4 and linear warmup over 15,000 steps. EvoDiff MSA models were trained on 16 32GB NVIDIA V100 GPUs for 10 days and saw ca. 3e9 tokens over 55,000 training steps.

### Baseline models

To enable direct comparison, the left-to-right autoregressive (LRAR) and CARP baselines were trained with the same CNN architectures on the same dataset as EvoDiff sequence models. For LRAR, the convolution modules have a causal mask to prevent information leakage. For additional MLM baselines, sequences were sampled from the protein MLMs ESM-1b (*68*) and ESM-2 (*30*), which were trained on different releases of UniRef50. ESM-1b and ESM-2 both generated many “unknown” amino acids (`X`); performance was improved results by manually setting the logits for `X` to − `inf`. Sequences were sampled from MLMs by treating the MLM as an OADMs and beginning from an all-mask state. For the structure-based diffusion baselines, sequences were obtained from FoldingDiff (*8*) and RFdiffusion (*10*) by first unconditionally generating structures and then using ESM-IF (*33*) to design their sequences. For MSA baselines, new query sequences were generated from ESM-MSA (*28*) by treating it as an OADM and sampling from an all-mask starting query sequence. CCMgen (*39*) with default parameters was used to train and generate from Potts models of validation MSAs from OpenFold.

### Computation of test-set perplexities

Perplexity was calculated by uniformly sampling a timestep for each test sequence, corrupting the sequence according to each diffusion model, predicting the sequence *x*_0_ at *t* = 0 by passing inputs once through each trained model, and then computing the perplexity. For D3PM models, the perplexity is:

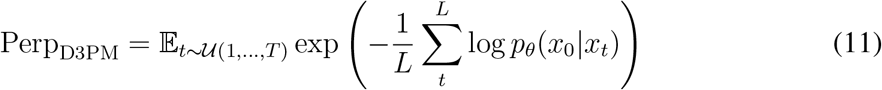

For OADMs, the perplexity is:

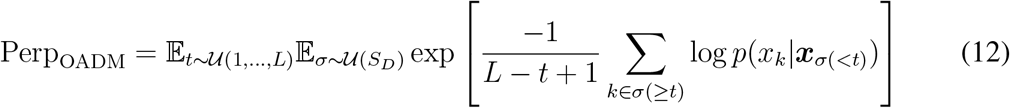

To enable model comparison, perplexities for MLMs (CARP, ESM-1b, ESM-2) were computed as if they are OADMs.

And for LRAR models, the perplexity is:

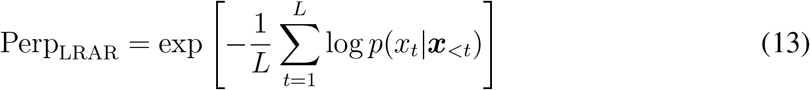

Calculated D3PM perplexities were on average higher as *t* → *T* and lower as *t* → 1, and masked perplexities were similarly higher for a greater number of masked tokens per sequence, i.e., as *t* → *L*_*masked*_ (**Fig. S1, S2**). Lower perplexities indicated improved performance and generalization capacity.

### Evaluation of structural plausibility

The structural plausibility pipeline (**Fig. 2A**) evaluates both the foldability and self-consistency of a given sequence. Foldability was evaluated by averaging the per-residue confidence score, reported as pLDDT by OmegaFold, across the entire sequence. Sequence self-consistency, denoted scPerplexity, describes how likely the generated sequence is to correspond to the predicted structure. Self-consistency was measured by taking structures predicted for a sequence from OmegaFold, running them through ESM-IF, and calculating the perplexity between the ESM-IF predicted-sequence and the original generated sequence.

The novelty of generated sequences was evaluated relative to training data seen by the model, by computing the Hamming distance between each generated sequence and every trainingset sequence of the same sequence length. The Hamming distance reported here is the distance between two sequences of equal length, normalized by sequence length. A distance of 1 indicates that there are no shared residues between two sequences, while a distance of 0 indicates that all the residues are shared between two sequences. The minimum of these Hamming distances, representing the closest sequence seen by the model during training, was reported for each sequence.

### BLAST protein homology search

Statistics on homology to natural sequences for all experimentally tested designs are provided in Supplementary Table 1. Each sequence was evaluated using the `blastp` tool in BLAST (*81*), comparing each protein sequence against BLAST’s non-redundant protein database, which includes GenBankCDS translations, RefSeq, PDB, Swissprot, PIR, and PRF. The % query coverage and % sequence identity to the top hit identified are reported. In some cases, there are no homologous sequences identified; these are denoted as having “no blast hits”.

### Computation of functional and structural features

To evaluate sequence coverage, ProtT5 embeddings were computed for each of 1,000 generated protein sequences and 10,000 sequences sampled from the test set using the Tools from Protein Prediction for Interpretation of Hallucinated Proteins (PPIHP) package (*82*). The resulting distributions of sequence embeddings (i.e., representing the corresponding distributions of sequences) were compared via the Fréchet ProtT5 distance (FPD),

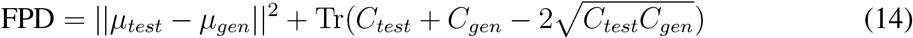

where, given the embedding space feature vectors for the test and generated distributions, *μ* is the feature-wise mean for each set of sequences, *C* is the respective covariance matrix, and Tr refers to the trace linear algebra operation, defined as the sum of the elements along the main diagonal of a square matrix. FPD quantifies the distance between the two distributions of ProtT5 embeddings and is a computationally tractable approximation of the Wasserstein distance between them. Embeddings were visualized in 2D via uniform manifold approximation and projection (UMAP), fit to the test data and with `n neighbors`=25. The number of neighbors hyperparameter was selected to favor local similarities in place of global ones, in order to appropriately visualize the corresponding differences in embedding space and FPD measured for each model.

Structural features of generated sequences were evaluated via the ProtTrans (*37*) CNN predictor model to assign a 3-state secondary structure definition from DSSP (helix, strand, or other) to each residue in a protein. The fraction of predicted ‘helix’, ‘strand’, or ‘other’ was computed (the three values sum to 1 per sequence). The resulting multivariate distributions of secondary structure features (computed over 1000 generated or natural sequences) were visualized via kernel density estimation. The KL divergence between the mean values across the 3-state predictions for the generated and test sets was used to quantitatively evaluate the distribution of secondary-structures assigned for each model.

### Evolution-guided generation with EvoDiff-MSA

Starting with either a random or Max-Hamming subsampled MSA, new query sequences were generated by sampling from an all-mask starting query sequence. The generated query sequence was evaluated relative to the corresponding original query sequence using the same tools and workflow described in *Evaluation of structural plausibility*. Each generated sequence was additionally evaluated for similarity relative to its reference MSA, which is comprised of a query sequence and alignment sequences. The % similarity of each generated sequence relative to its parent MSA was computed as the maximum % similarity over all sequences in the original MSA. Specifically, for a pair of sequences, the % similarity was computed by calculating the number of shared residue identities (accounting for both amino-acid identity and position index in the sequence), and for a given generated sequence the maximum value of these % similarities was determined. Across generated sequences both the CDF and mean of maximum % similarity were reported. Generated sequences were additionally evaluated for structural similarity relative to their original query sequences. Structures were predicted for each of the generated query sequences and the original query sequences using OmegaFold. Structural similarity was measured via the template modeling score (TM-score) (*83*) for the two predicted structures following structural alignment:

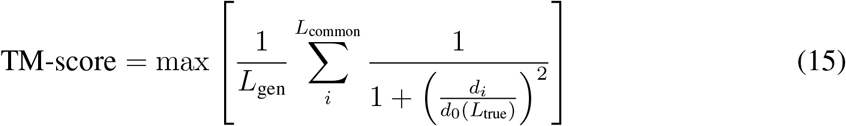

where *L*_gen_ is the length of the generated query sequence; *L*_common_ is the number of shared residues; *d*_*i*_ is the distance between the *i*^*th*^ pair of residues; *L*_true_ is the length of the true query sequence; and 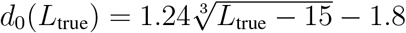 is a distance scale for normalization.

### Overview of protein expression, purification, and functional assays

Designs were expressed in mammalian (Chinese Hamster Ovary, CHO), bacterial (*Escherichia coli, E. coli*), yeast (*Saccharomyces cerevisiae, S. cerevisiae*), or cell-free systems; a mapping of design to expression system is provided in Supplementary Table 1. Unconditional generations were expressed, purified, and assayed by WuXi Biologics. Designed N-terminal IDRs for Cox15 were tested via cell-based assays in *S. cerevisiae*. Designs scaffolding the calmodulin Ca^2+^-binding motif were expressed, purified, and assayed by WuXi Biologics. Designs scaffolding the p53 transactivation domain were expressed and assayed by Adaptyv Bio. All experimental details are provided in the corresponding Methods sections.

### Mammalian cell expression and downstream purification of unconditional generations and calmodulin scaffolds

A set of unconditional generations and scaffolds of the calmodulin Ca^2+^-binding motif were expressed by WuXi Biologics in CHO cell expression systems (Supplementary Table 1). The WuXi Biologics Proprietary CHO K1 cell line was adapted from CHO-K1 cells (ATCC) to grow as suspension culture in WuXi’s proprietary culture and feeding media (WuXi Biologics). Briefly, proteins were initially tested by WuXi’s Ultra 96 platform, which allows for the expression of a greater number of proteins at a lower scale. Designs that expressed successfully were then produced by WuXi’s WuXian transient platform in CHO K1 cells (WuXi Biologics) scaled up to 0.2-1.0L scale; purification followed successful expression at scale. When expressed successfully in CHO cells, protein samples used in downstream assays were derived from scaled expression in CHO, though certain designs also expressed in *E. coli* (Supplementary Table 1). Brief details on scaled expression in CHO cells and subsequent purification are provided below.

DNA sequences of target proteins were codon optimized and inserted into a proprietary backbone plasmid (WuXi Biologics). Synthesized constructs contained a CMV promoter inframe to express and secrete target proteins into the cell culture media. Constructed vectors were transformed into chemically competent *E. coli* TOP10 cells following manufacturer’s instruction, and plasmids were isolated from successful colonies and then sequence confirmed. DNA was transfected into CHO K1 host cells, and transfected cultures were incubated by shaking at 150 rpm for 7 days.

Cell culture supernatants were harvested by primary centrifugation at 10,000 x g for 40 min, followed by sterile filtration. Supernatants were purified by affinity capture chromatography with Ni-Excel resin (Cytiva), with step imidazole elution, conducted on the AKTA Pure 25 column chromatography system (Cytiva). Following affinity capture chromatography, samples were further purified via size exclusion chromatography on Superdex 75 pg resin (Cytiva) in PBS. Relevant fractions were pooled, and final products were subject to quality control testing, including via SDS-PAGE, HPLC (Vanquish Duo UHPLC; ThermoFisher Scientific) and LC-MS (6545XT AdvanceBio LC/Q-TOF; Agilent). Final products were stored at -80C.

### *E. coli* expression of unconditional generations

For *E. coli* expression of unconditional generations (EvoDiff-Seq-Design-2; EvoDiff-Seq-Design-3), assembly with NcoI and XhoI was used to insert constructs (Genewiz) into the pET28a vector containing 6xHis tags. Plasmids were transformed into *E. coli* BL21(DE3). Large-scale cultures were grown at 5L (EvoDiff-Seq-Design-2; EvoDiff-Seq-Design-3) scale in auto-induction medium (*84*) for 2.5hrs at 37C pre-induction and 20hrs at 20C for induction. Though EvoDiff-Seq-Design-1 also successfully expressed in *E. coli* (Supplementary Table 1), its scaled-up expression occurred in CHO, followed by downstream purification as described above.

Following growth, cultures were harvested via centrifugation at 8,000 x g for 20 min at 4C. Pellets were suspended at 5 mL / 1 g pellet in lysis buffer (for EvoDiff-Seq-Design-2, EvoDiff-Seq-Design-3: 20 mM Tris, 300 mM NaCl, 20 mM Imidazole, 1 mM TCEP, 10% v/v glycerol, 0.1% v/v Triton X-100, with EDTA-free protease inhibitor cocktail, pH 8.0; for 1PRW and EvoDiff-Seq-1PRW-Design-5: 50 mM Tris, 150 mM NaCl, 1 mM TCEP, 10% v/v glycerol, with EDTA-free protease inhibitor cocktail, pH 8.0) and incubated at 4C for 1hr with stirring. High-pressure homogenization was used to lyse the cells (3x homogenization at 1000 bar), and the material was pelleted via centrifugation at 40,000 x g for 60 min at 4C. For EvoDiff-Seq-Design-2 and EvoDiff-Seq-Design-3, pellets were dissolved in a second lysis buffer (5 mL / 1 g pellet; 20 mM Tris, 300 mM NaCl, 20 mM Imidazole, 1 mM TCEP, 3 M Urea, 10% v/v glycerol, 0.1% v/v Triton X-100, pH 8.0), incubated at 4C for 1hr with stirring, then homogenized (2x homogenization at 1000 bar). Following centrifugation, the clarified supernatant was collected.

### Purification of *E. coli*-expressed unconditional generations

Following centrifugal lysate clarification, proteins were first purified by affinity chromatography at 4C. Clarified supernatant was equilibrated into equilibration buffer (EvoDiff-Seq-Design-2, EvoDiff-Seq-Design-3: 20 mM Tris/HCl, 300 mM NaCl, 20 mM Imidazole, 0.5 mM TCEP, 3 M Urea, 10% v/v glycerol, pH 8.0), loaded onto Ni-NTA 3 mL affinity chromatography matrix (Ni-NTA Superflow, Qiagen), washed once with equilibration buffer, washed twice with Ni-NTA wash buffer (20 mM Tris/HCl, 300 mM NaCl, 20 mM Imidazole, 0.5 mM TCEP, 10% v/v glycerol, pH 8.0), and eluted with Ni-NTA elution buffer (20 mM Tris/HCl, 300 mM NaCl, 250 mM Imidazole, 0.5 mM TCEP, 10% v/v glycerol, pH 8.0).

Following affinity chromatography, purification of *E. coli*-expressed unconditional generations proceeded as follows. Throughout, intermediate samples were verified by SDS-PAGE. The purity and molecular weight of final samples were confirmed by liquid chromatography-mass spectrometry (LC-MS; Agilent).

For EvoDiff-Seq-Design-2 and EvoDiff-Seq-Design-3, following affinity chromatography, samples were further purified by Q-HP anion exchange chromatography (HiTrap Q HP anion exchange chromatography column, Cytiva) at 4C. EvoDiff-Seq-Design-2 and EvoDiff-Seq-Design-3 samples were diluted 6x in dilution buffer (20 mM Tris/HCl, 0.5 mM TCEP, 3 M Urea, 10% v/v glycerol, pH 8.0), equilibriated, loaded onto the column, washed with equilibriation buffer (20 mM Tris/HCl, 50 mM NaCl, 0.5 mM TCEP, 3 M Urea, 10% v/v glycerol, pH 8.0), and eluted (20 mM Tris/HCl, 1 M NaCl, 0.5 mM TCEP, 3 M Urea, 10% v/v glycerol, pH 8.0) using a concentration gradient of 0-60%, 60-100% for elution.

For EvoDiff-Seq-Design-2, following Q-HP purification, monomeric fractions were pooled, and samples were buffer exchanged overnight at 4C into a final buffer (50 mM Tris/HCl, 300 mM NaCl, 0.5 mM TCEP, pH 7.5) and stored.

For EvoDiff-Seq-Design-3, following Q-HP purification, the protein was further purified at scale-up via reverse-phase (RP) chromatography. For RP chromatography, pooled protein from Q HP purification was adjusted to pH of 4.0 with acetic acid in a final solution with 5% v/v acetonitrile. The sample was loaded onto a Sepax Bio-C8 column (Sepax Technologies), washed with a solution of 0.1% v/v TFA, 5% v/v acetonitrile in ddH2O, and then eluted (0.1% v/v TFA in acetonitrile) using a concentration gradient of 5-100%, with sample eluting at a concentration of ca. 50% elution buffer. The final sample was frozen in liquid nitrogen and freeze-dried in a vacuum dryer.

### Structural characterization by UV circular dichroism

Circular dichroism (CD) was used to characterize higher-order structures in EvoDiff-designed proteins. All CD data collection was performed with the Chirascan V100 CD spectrometer (Applied Photophysics), and analysis was conducted via the Chirascan software (Applied Photophysics). Briefly, secondary structure characteristics were determined via measurement of the far-UV CD spectra. Prior to measurement, protein samples were concentrated and then diluted with ultrapure water to a final concentration of 0.1 mg/mL. Measurements were acquired in the far-UV spectra ranging from 190 to 260nm; all reported measurements were acquired within the linear range of the instrument. Analysis of secondary structural composition was determined using the CDNN software with a net setting using 33 basespectra. CD experiments were conducted by WuXi Biologics.

### Generation of intrinsically disordered regions (IDRs)

IDR generation and analysis leveraged a publicly available dataset of 15,996 human IDRs and their orthologs (*48*). This dataset was generated by running SPOT-Disorder v1 (*85*) on the human proteome and applying the predicted IDR positions to an MSA of likely-similar-function orthologs (determined using an evolutionary distance heuristic), curated from the larger set of orthologs contained in the OMA database (*86*). The resulting dataset only contained IDRs, not full protein sequences, and thus IDR sequences were mapped back to the MSAs of full protein sequences in OMA in order to provide context about the sequence regions surrounding the IDRs.

For input to EvoDiff models, the full sequence of an IDR-containing human protein was treated as the query sequence, and a corresponding MSA was constructed by subsampling 63 other sequences from all the query’s orthologs. All sequences were subsampled to 512 residues in length, with the following criteria maintained. Subsampling criteria were that the subsampled query sequence contain at least 1 IDR, and that the total IDR region was less than half the total length of the subsampled sequence (*L*_IDR_ ≤ 256). For IDR generation from EvoDiff-Seq, the query sequence with the IDR region masked was provided as the only input to EvoDiff-Seq, which then generated new residues for the masked region (i.e., the region corresponding to the true IDR). For IDR generation from EvoDiff-MSA, the query sequence with the IDR region masked, aligned to the rest of the MSA, was provided as input to EvoDiff-MSA, which then generated new residues for the masked region.

The resulting generations, containing putative IDRs, were input to DR-BERT, a protein language model fine-tuned for disorder prediction (*49*), to obtain per-residue disorder scores ranging from 0-1 (less to more disordered). A single-sequence IDR predictor (DR-BERT) was used in place of MSA-based IDR scoring methods, because of an observed bias towards higher disorder scores with MSA-based methods – e.g., random uniform sampling of residues in the masked query positions still resulted in a prediction of disorder given the presence of the orthologs in the alignment. Disorder scores for true IDRs, generated IDRs, scrambled IDRs, and randomly generated IDRs were computed to evaluate the performance of DR-BERT predictions. The randomly-sampled baseline was constructed by randomly sampling amino acids over an IDR region; the scrambled baseline was constructed by shuffling the existing amino acids over an IDR region into a scrambled permutation. In all cases (true IDRs, generated IDRs, scrambled and random baselines), the entire protein sequence was input to DR-BERT for scoring. Since DR-BERT is for single-sequences, for putative IDRs generated by EvoDiff-MSA, the entire query sequence was inputted into DR-BERT, with < `GAP` > tokens eliminated, to obtain per-residue disorder scores. Lastly, a direct comparison between the original IDR and the generated putative IDR was conducted by calculating the % sequence similarity between the fraction of shared residues between the two IDR regions.

### Design of N-terminal IDRs for Cox15

The yeast Heme A synthase Cox15 was used as a target for the design of synthetic N-terminal IDRs. A total of 600 inpainted designs for the disordered N-terminal IDR were generated using EvoDiff-Seq and EvoDiff-MSA (Rand subsampling). The first 200 designs were generated by masking the N-terminal IDR (amino acid positions 1 to 45, inclusive) and randomly decoding at each position in the sequence with EvoDiff-Seq. Another 200 sequences were generated by iteratively re-masking and generating at each position of the IDR in order from left to right. Both design strategies are equally represented in the final tested candidates. The last 200 designs were generated using EvoDiff-MSA, using the alignment created for Cox15 from Lu et al. (*48*). The yeast Cox15 sequence was set as the query sequence, and an alignment of 63 sequences was randomly subsampled. For each generation, a start token within the Cox15 query was randomly sampled; this results in varying N-terminal inpainting lengths for each independent generation. The masked N-terminal region was then decoded in random order. To nominate 8 designs for experimental validation, all 600 designs were scored for and then ranked by the probability of disorder by DR-BERT and the likelihood of mitochondrial localization by MitoFates (**Fig. S14**). For each of EvoDiff-Seq and EvoDiff-MSA, the top four designs with the largest edit distance to the native IDR N-terminal sequence of Cox15 were selected for experimental validation (**Supplementary Table 1**).

### Yeast experimental methods for design of synthetic IDRs

Synthetic IDRs designed to be mitochondrial targeting signals for the yeast protein Cox15 were tested *in vivo* via a subcellular localization assay, readout by confocal microscopy. A tagged Cox15-mScarlet serves to show wild-type localization of the native protein and provides an internal control for direct comparison to the synthetic IDRs tested in this study. This control strain was constructed by tagging the endogenous locus of Cox15 at the C-terminus with mScarlet followed by a selectable resistance marker (KANMX4) via homologous recombination in *S. cerevisiae* S288C derivative (BY4741: MATa his3∆1 leu2∆0 met15∆0 ura3∆0) (*87*), using the standard LiAc method (*88*), and isolated by marker selection on yeast synthetic minimal media (SD). The corresponding strain with the mScarlet-tagged Cox15 full-length ORF served as the parental strain for all subsequent strains in this study; the mScarlet-tagged Cox15 full-length ORF remained intact in all experimental strains.

The experimental Cox15 strains, containing wild type or synthetic IDR sequences, were made by inserting a second copy of the Cox15 ORF tagged with SuperFold-GFP (sfGFP) (*89*) at the HO locus of the Cox15-mScarlet control strain via recombinant homologous transformation as described above. Synthetic amino acid sequences were obtained via computational design, and corresponding DNA sequences were codon optimized for optimal expression in *S. cerevisiae* using a program available from Integrated DNA Technologies (https://www.idtdna.com/CodonOpt). The resultant nucleotide sequences were then used to design gene fragments (GeneArt Strings). These gene fragments served as PCR templates to generate sequences that were then inserted in plasmids via Gibson Assembly (*90*), replacing the native Cox15 IDR in the full length Cox15 ORF, transformed into *E. coli* (DH5*α*), and then isolated by marker selection (Ampicillin). All plasmids were purified from bacteria and confirmed by Sanger sequencing (Eurofins Scientific SE). Positive clones were subsequently used for the yeast genomic integration. The transformed linear DNA fragments contained the native Cox15 promoter, the Cox15 ORF (containing the synthetic IDRs) fused to super-fold GFP (sfGFP), followed by the tADH1 terminator and an auxotrophic yeast marker (URA3) for selection. For the IDR knockout strain, a previously constructed strain containing an endogenous Cox15 2-45 deletion was used as a PCR template to generate the IDR deletion. Following transformation and cloning, positive clones were selected for yeast genomic integration, and were integrated into the same locus (HO) as all other strains via direct homologous recombination in yeast cells. This was a two fragment integration, with the second fragment providing the remaining Cox15 ORF fused to sfGFP and the tAdh1 terminator sequence, followed by the same auxotrophic marker used in the other strains. The HO locus was amplified from all yeast strains via colony PCR, and strain identity was confirmed by Sanger sequencing (Eurofins Scientific SE).

### Confocal microscopy and quantification of subcellular localization

Confocal images of yeast strains were obtained as follows. Control and mutant strains were inoculated into yeast synthetic minimal media (SD) and grown overnight at 30C. Prior to imaging, stationary cultures were diluted 1/10 in fresh media and grown at 30C for 4 hours to ensure log-phase growth and proper expression of mScarlet/GFP fusion products. Live cells were concentrated 50-fold by centrifugation (30s at 3,500 x g) and imaged using a Leica TCS SP8 confocal microscope with a 63X oil-immersion objective and a 10X eye-piece objective. mScarlet excitation was at 587nm and GFP excitation was at 488nm.

To quantify co-localization, confocal images were first segmented using the YeastSpotter segmentation model (*91*) applied on the brightfield image. All segmentation outputs under 2000 pixels were filtered to remove dust particles and small budding cells (which are typically out of focus). For each cell, co-localization was quantified by computing the Pearson correlation coefficient between the mScarlet channel (587nm excitation) and the GFP channel (488nm), for all pixels assigned to the cell by YeastSpotter.

### Motif scaffolding

*In-silico* scaffolding performance was evaluated on a recently published benchmark (*10*) of 25 scaffolding problems across 17 unique proteins.

In our scaffolding benchmark, each unique protein was treated as 1 example, for a total of 17 unique scaffolding examples. 100 samples were generated for each unique scaffolding example. For proteins 6E6R, 6EXZ, 7MRX, and 5TRV, which were the 4 examples evaluated at 3 different scaffolding lengths in RFdiffusion (*10*), the number of successes across these three different scaffolding lengths were averaged to facilitate comparisons between RFDiffusion and EvoDiff.

To generate a scaffold with EvoDiff-Seq, a scaffold length between 50-100 residues (exclusive of the motif) was sampled uniformly; the motif was placed randomly within the length; and scaffold residues were generated from EvoDiff-Seq conditioned on the provided motif residues. In this approach, on average, protein sequences generated by EvoDiff-Seq were longer (between 45 and 194 residues in length) than those inverse-folded from structures generated by RFdiffusion, which range from 30-152 residues in total length inclusive of the length of the motif.

For scaffolding with EvoDiff-MSA, MSAs for each sequence corresponding to the original PDB structure were generated using the tools from AlphaFold (*92*) and then subsampled to 64 sequences, and a maximum of 150 residues in length, where the original sequence obtained from the PDB crystal structure was assigned as the query sequence. In cases where the scaffolding examples were shorter than 150 residues, sequences were padded with a < `GAP` > token, to allow EvoDiff to generate longer-scaffolds. Sequences generated by EvoDiff-MSA were between 56 and 150 residues in length, inclusive of the motif and scafffold. For each scaffolding example, a common set of 100 subsampled MSAs, where 50 were randomly subsampled and 50 were subsampled via MaxHamming, was used commonly across EvoDiff-MSA (Max), EvoDiff-MSA (Random), and ESM-MSA. That is, an individual generation trial for each model corresponded to a unique MSA from the common set of 100 MSAs constructed for a scaffolding example. At inference time, all non-motif residues in the query sequence were masked, and new residues in these locations were generated by EvoDiff-MSA.

OmegaFold was used to predict structures corresponding to sequences generated by EvoDiff. A generation was counted as ‘successful’ if its predicted structure had a pLDDT >70 and a motifRMSD <1.0 *Å* relative to the original motif crystal structure. Note that these success criteria are cutoffs proposed by structure-based models (*10*) and adopted here to facilitate comparison. The motifRMSD was computed as the RMSD between the alpha-carbons of the motif in the original crystal structure and the predicted structure for the scaffolded motif.

### Design of experimentally-tested calmodulin scaffolds

EvoDiff-Seq, EvoDiff-MSA (Rand subsampling), and EvoDiff-MSA (Max subsampling) were used to design 100 calmodulin scaffolds each, resulting in 300 designs total. Generations from EvoDiff-Seq began by isolating the residues of the functional motif from the sequence of bovine calmodulin from the PDB entry 1PRW. The subsequence of residues 16-70 (inclusive), which contains the functional Ca^2+^-binding motif, was considered, and residues 35-50 (inclusive) were masked, such that only the residues of the functional motif remained. Then, for each EvoDiff-Seq generation, a total scaffold length between 10 and 100 residues was sampled and set to all mask tokens, with the start position of the motif positioned randomly within the scaffold. A sequence was generated from EvoDiff-Seq by randomly decoding at each position. For generation with EvoDiff-MSA, an alignment was constructed using the homology tools from AlphaFold. 63 sequences were subsampled from the alignment using Rand or Max subsampling, using a maximum sequence length of 150. All residues that did not belong to a motif were then masked, followed by decoding at each position randomly. To select designs for experimental validation, all 300 designs were compiled and filtered by the designability metrics of OmegaFold pLDDT > 70 and motifRMSD < 1.0Å. Designs that scored well by these metrics but seemed physically unfeasible (e.g., long unconstrained helices) were manually filtered out, and 13 designs were nominated for experimental validation (**Supplementary Table 1**).

### *E. coli* expression of calmodulin scaffolds

Except for the native bovine calmodulin (1PRW) and EvoDiff-Seq-1PRW-Design-5, all other calmodulin scaffolds successfully expressed in the scaled CHO expression system and were subsequently purified (see prior Methods section; **Supplementary Table 1**); these scaffolds also successfully expressed in *E. coli*, but expression was not scaled up, and thus, for these designs, purified samples derived from CHO expression were used in the downstream absorbance assay. Scaled expression of 1PRW and EvoDiff-Seq-1PRW-Design-5 occurred in *E. coli*.

For scaled expression of 1PRW and EvoDiff-Seq-1PRW-Design-5 in *E. coli*, assembly with NcoI and XhoI was used to insert constructs (Genewiz) into the pET28a vector containing 6xHis tags. Plasmids were transformed into *E. coli* BL21(DE3). Large-scale cultures were grown at 0.5L (EvoDiff-Seq-1PRW-Design-5) or 1L (1PRW) scale in auto-induction medium (*84*) for 2.5hrs at 37C pre-induction and 20hrs at 20C for induction.

Following growth, cultures were harvested via centrifugation at 8,000 x g for 20 min at 4C. Pellets were suspended at 5 mL / 1 g pellet in lysis buffer (for 1PRW and EvoDiff-Seq-1PRW-Design-5: 50 mM Tris, 150 mM NaCl, 1 mM TCEP, 10% v/v glycerol, with EDTA-free protease inhibitor cocktail, pH 8.0) and incubated at 4C for 1hr with stirring. High-pressure homogenization was used to lyse the cells (3x homogenization at 1000 bar), and the material was pelleted via centrifugation at 40,000 x g for 60 min at 4C. Following centrifugation, the clarified supernatant was collected.

### Purification of *E. coli*-expressed calmodulin scaffolds

Following centrifugal lysate clarification, proteins were purified by affinity chromatography at 4C. Clarified supernatant was equilibrated into equilibration buffer (for 1PRW, 1PRW 37: 20 mM Tris/HCl, 300 mM NaCl, 20 mM Imidazole, 0.5 mM TCEP, 10% v/v glycerol, pH 8.0), loaded onto Ni-NTA 3 mL affinity chromatography matrix (Ni-NTA Superflow, Qiagen), washed once with equilibration buffer, and eluted with Ni-NTA elution buffer (20 mM Tris/HCl, 300 mM NaCl, 250 mM Imidazole, 0.5 mM TCEP, 10% v/v glycerol, pH 8.0).

Following affinity chromatography, purification of *E. coli*-expressed 1PRW and EvoDiff-Seq-1PRW-Design-5 proceeded as follows. Throughout, intermediate samples were verified by SDS-PAGE. The purity and molecular weight of final samples were confirmed by liquid chromatography-mass spectrometry (LC-MS; Agilent).

For 1PRW, following affinity chromatography, the sample was subject to a digestion step with TEV protease (20:1 w/w ratio of 1PRW to TEV; TEV expressed and purified in-house by WuXi Biologics), in order to remove the His tag. Reverse Ni-NTA chromatography of the digested protein sample was subsequently performed, with the sample equilibriated into equilibriation buffer (20 mM Tris/HCl, 300 mM NaCl, 20 mM Imidazole, 0.5 mM TCEP, 10% v/v glycerol, pH 8.0), loaded into the column, washed once with equilibriation buffer, and then eluted with Ni-NTA elution buffer. The sample was then subject to size exclusion chromatography (SEC) (HiLoad Superdex 75 Prep Grade; Cytiva; 50mM Tris/HCl, 0.3 M NaCl, 0.5 mM TCEP, 10% v/v glycerol, pH 7.5). Following SEC, the 1PRW sample was subject to Q-HP anion exchange chromatography (HiTrap Q HP anion exchange chromatography column, Cytiva) at 4C. Samples were diluted 6x in dilution buffer (20 mM Tris/HCl, 0.5 mM TCEP, 10% v/v glycerol, pH 7.5), equilibrated and loaded onto the column (20 mM Tris/HCl, 50 mM NaCl, 0.5 mM TCEP, 10% v/v glycerol, pH 7.5), washed with equilibration buffer, and eluted (20 mM Tris/HCl, 1 M NaCl, 0.5 mM TCEP, 10% v/v glycerol, pH 7.5) using a concentration gradient of 0-60%, 60-100% for elution. 1PRW monomeric fractions were pooled, and samples were buffer exchanged overnight at 4C into a final buffer (50 mM Tris/HCl, 300 mM NaCl, 0.5 mM TCEP, pH 7.5) and stored.

For EvoDiff-Seq-1PRW-Design-5, following affinity chromatography, the sample was purified via SEC (HiLoad Superdex 75 Prep Grade; Cytiva; 50mM Tris/HCl, 0.3 M NaCl, 0.5 mM TCEP, 10% v/v glycerol, pH 7.5). Monomeric fractions were pooled, and samples were stored.

### Spectroscopic analysis of Ca^2+^ binding to calmodulin and designed scaffolds

Analysis of Ca^2+^ binding to recombinantly-expressed bovine calmodulin (1PRW) and EvoDiff-designed scaffolds was performed via a spectroscopic assay adapted from previous reports (*55, 56*). Purified proteins were incubated at 50 *μ*M final concentration with 32x molar excess CaCl_2_ (1.6mM; MCE) for 16 hrs at room temperature in assay buffer (20mM HEPES, 100mM KCl, pH 7.8). Following the 16h incubation, absorbance spectra were collected at wavelengths from 450nm to 650nm on a SpectraMax M2e Microplate Reader (Molecular Devices). Measurements were background-corrected using readings from the buffer-only controls. Successful binders showed characteristic absorbance peaks at 500-550 nm relative to the absorbance spectra for no-protein and buffer controls.

### Design of experimentally-tested p53 scaffolds

To design scaffolds for the transactivation domain of p53, 900 designs from EvoDiff-Seq, 200 from EvoDiff-MSA (Rand subsampling), and 200 from EvoDiff-MSA (Max subsampling) were generated, resulting in 1300 total designs to select from. For EvoDiff-Seq, the p53 transactivation domain, residues 5-15 (inclusive) from the PDB entry 1YCR, was scaffolded at variable scaffold lengths between 50 and 100 residues. Generations were decoded by sampling the amino acid probabilities at each position with various temperatures (*T*). 500 examples were generated at *T* = 1.0, 200 at *T* = 0.8, and another 200 at *T* = 0.7. For generation with EvoDiff-MSA, an alignment based on the full sequence of human p53 (Uniprot entry: P04637) was constructed using the homology tools from AlphaFold; the transactivation domain within 1YCR is from the human p53 sequence. 63 sequences containing the scaffolding motif were subsampled from the alignment using Rand or Max subsampling, using a maximum sequence length of 150. All residues that did not belong to the motif in the query sequence were then masked, followed by decoding at each position randomly without temperature sampling. The 900 generations from EvoDiff-Seq were rank ordered by the designability metrics of OmegaFold pLDDT and motifRMSD order and a set of 19 candidates was nominated (**Supplementary Table 1**). None of the designs from EvoDiff-MSA passed the *in-silico* pLDDT criterion (pLDDT >70), likely due to the native intrinsically disordered region within p53 (**Fig. 6G**). EvoDiff-MSA designs were thus filtered to those with motifRMSD <1.0 *Å*, and five sequences (3 Rand subsampling, 2 Max subsampling) were randomly selected such that three out of five sequences had shared homology with the native p53 sequence (**Supplementary Table 1**).

### Cell-free expression of p53 scaffolds

Designed sequences scaffolding the p53 transactivation domain were synthesized *in vitro* via a cell-free expression platform (Adaptyv Bio). DNA constructs encoding the ligands, each fused to a C-terminal assay tag, were designed by reverse-translating the target protein sequences. The sequences were optimized for manufacturability and yield using Adaptyv’s proprietary algorithms to maximize expression efficiency. The optimized DNA constructs were ordered as gene fragments from Twist Bioscience.

Assembly of the constructs into an appropriate expression vector was performed using the NEBuilder HiFi DNA Assembly Kit (New England Biolabs) in 3 *μ*L reactions. Protein expression was carried out in 8 *μ*L reactions using an optimized prokaryotic in-vitro translation system. Reactions were incubated at 37C for 12 hours to ensure efficient protein synthesis. Post-expression, protein concentration and yield were normalized using an affinity-based quantification assay. All tested synthetic designs and all positive controls expressed successfully.

### Affinity characterization using biolayer interferometry

Designed p53 scaffolds and positive controls (i.e., the ligands) were tested for their *in vitro* binding affinity to MDM2 (i.e., the antigen). The p53 transactivation domain peptide from 1YCR (p53 control) and two synthetic binders from RFDiffusion (*10*) (p53 RF 53 and p53 RF 89) were used as positive controls. Sequences of all tested designs as well as of the target antigen are provided in **Supplementary Table 1**.

Multi-cycle kinetic assays were conducted in triplicate to evaluate ligand binding to the antigen using Biolayer Interferometry (BLI) (Gator Bio). Ligands, tagged with a twin strep tag, were immobilized on streptavidin-coated probes via tag-specific interactions. Antigen solutions at multiple concentrations (500 nM, 158.1 nM, 50 nM, 15.8 nM, and 0 nM) of full-size, recombinant human GST-MDM2 (Bio-Techne E3-202-050) were flowed over the probes, with signal acquisition at a sampling rate of 5 Hz. The assay sequence began with a 120-second baseline phase, followed by 120 seconds of ligand loading onto the probes. After loading, a secondary 200-second baseline was established to ensure probe stability. The association phase, during which the antigen was introduced, lasted 180 seconds, followed by a 240-second dissociation phase to monitor antigen unbinding from the immobilized ligand.

All experiments were conducted in kinetic buffer (PBS-T supplemented with 0.02% w/v BSA) at 25°C. Negative controls, including buffer-only samples, were included for baseline subtraction. Data were corrected based on the baseline values and fitted globally to a 1:1 binding model across all concentrations. Kinetic rate constants and dissociation constants (*K*_*D*_) were derived from the fitted curves.

## Supplementary Information for

**Figure S1:**
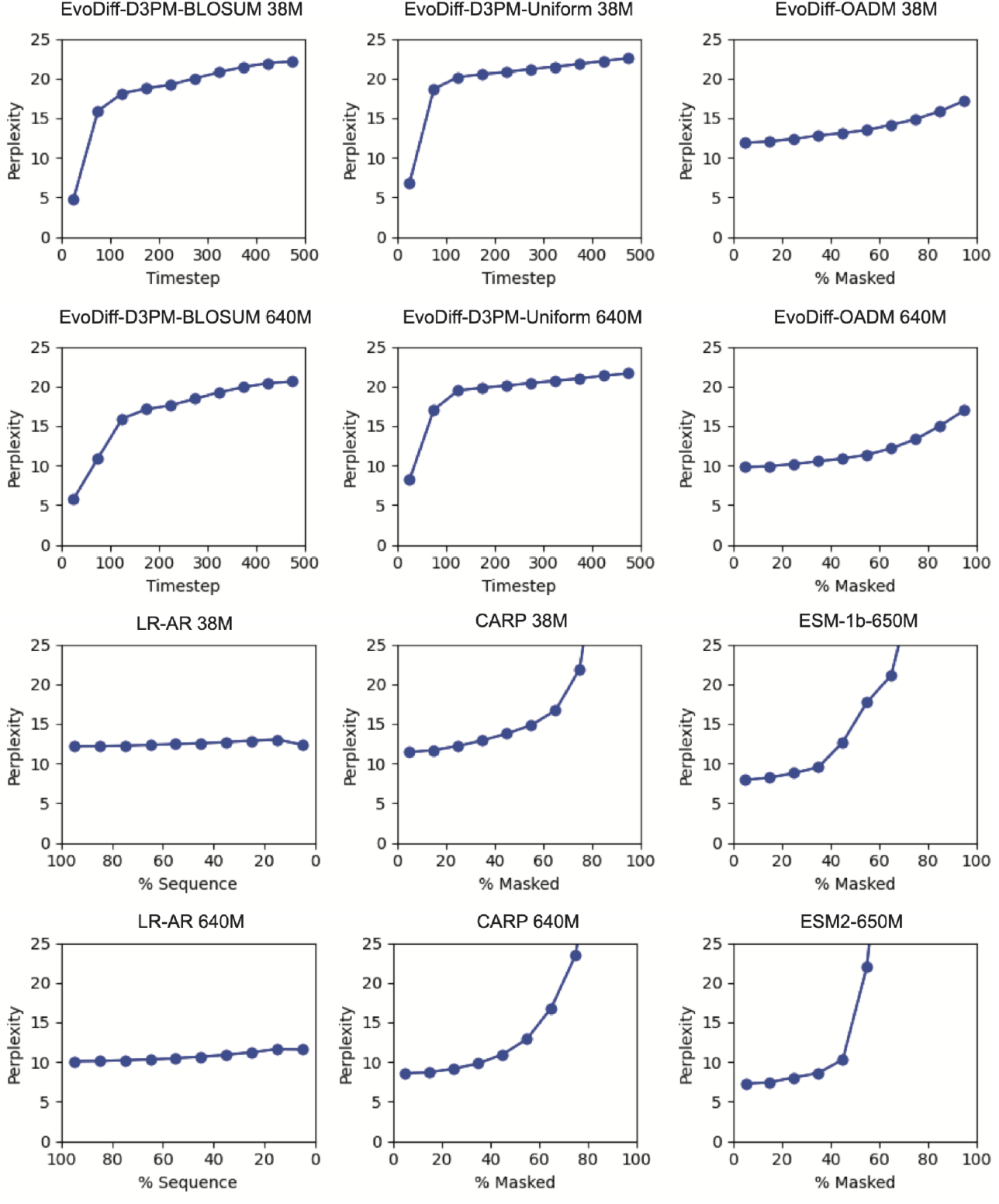
Perplexity as a function of corruption step for EvoDiff sequence models. Test-set perplexities at sampled intervals of the degree of corruption, specifically the diffusion timestep for D3PM models, the fraction of masked residues for OADM and masked language models, and the fraction of evaluated sequence for LRAR models. Intervals reflect evenly spaced windows of 50 timesteps for D3PM models or 10% masking for masked models.

**Figure S2:**
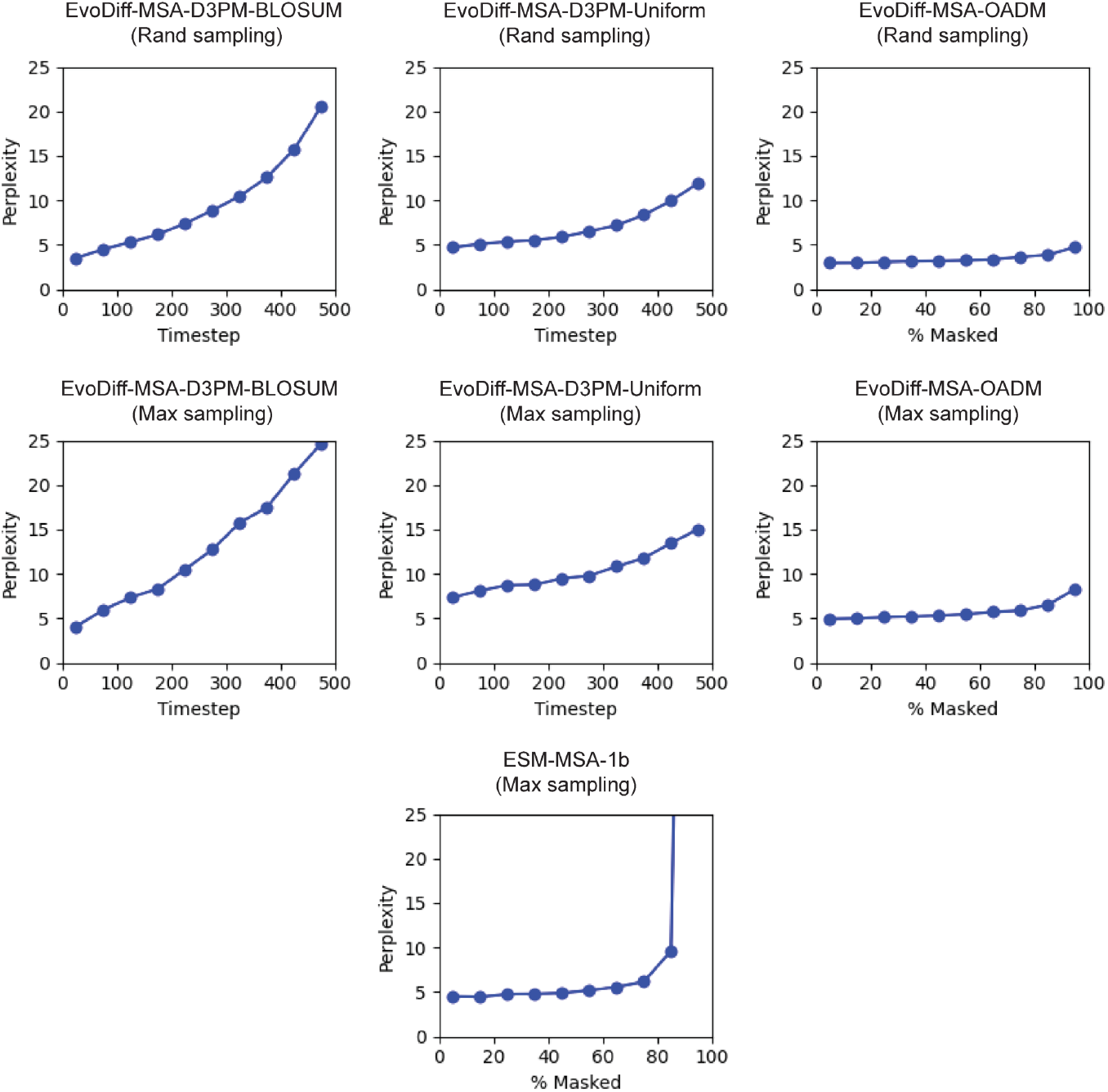
Perplexity as a function of corruption step for EvoDiff MSA models. Test-set MSA perplexities at sampled intervals of the degree of corruption, specifically the diffusion timestep for D3PM models and the fraction of masked residues for OADM and ESM models. The test-set evaluated for each model was sampled using the same sampling scheme assigned during training. Intervals reflect evenly spaced windows of 50 timesteps for D3PM models or 10% masking for masked models.

**Figure S3:**
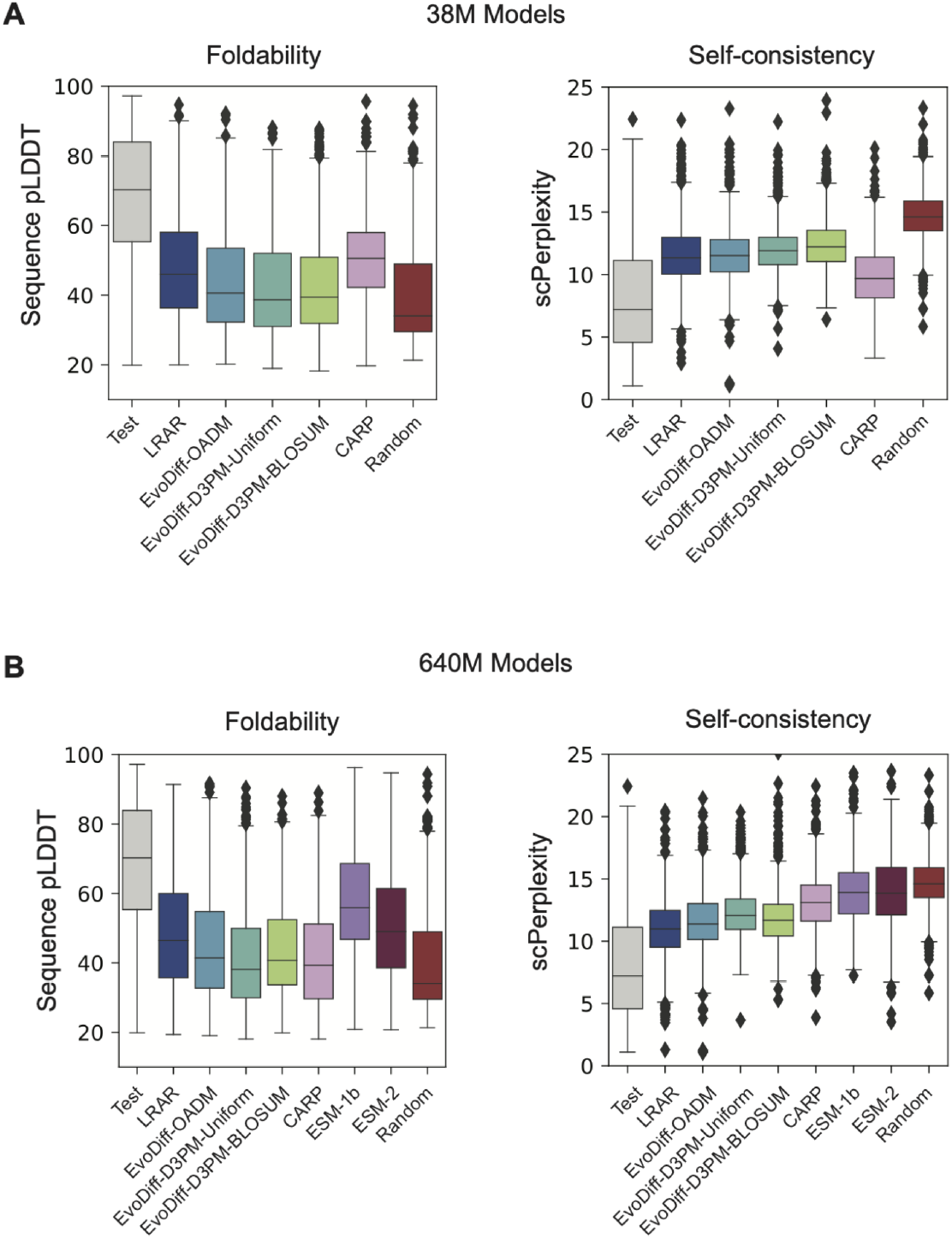
Summary statistics for structural plausibility metrics for sequence models. **(A-B)** Distribution of pLDDT and scPerplexity metrics for sequences from the test set, 38M parameter EvoDiff and baseline models (A), and 640M parameter EvoDiff and baseline models (B) (n=l000 sequences per model). Test and Random baselines are reproduced in (A) and (B) for reference.

**Figure S4:**
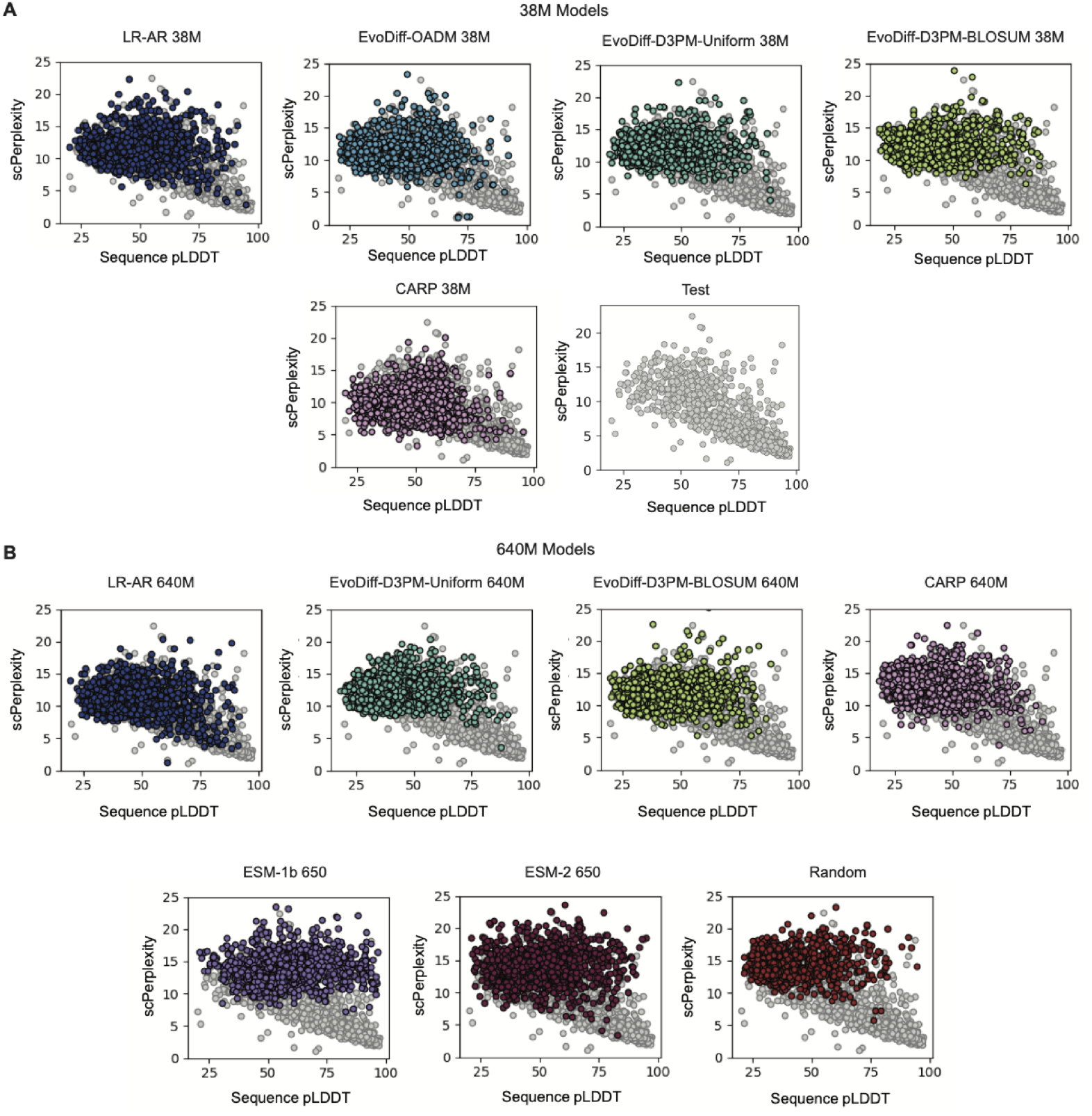
Sequence pLDDT versus self-consistency perplexity for EvoDiff sequence models. **(A-B)** Results for sequences from 38M parameter EvoDiff, baseline models, and test data plotted alone (A), and 640M parameter EvoDiff and baseline models (B) (various colors, *n=*1000), except for EvoDiff-OADM-640M (EvoDiff-Seq, shown in Fig. 2C), relative to sequences from the test set (grey, *n=*1000). The self-consistency perplexity (ESM-IF Perplexity) is computed using sequences inverse-folded by ESM-IF.

**Figure S5:**
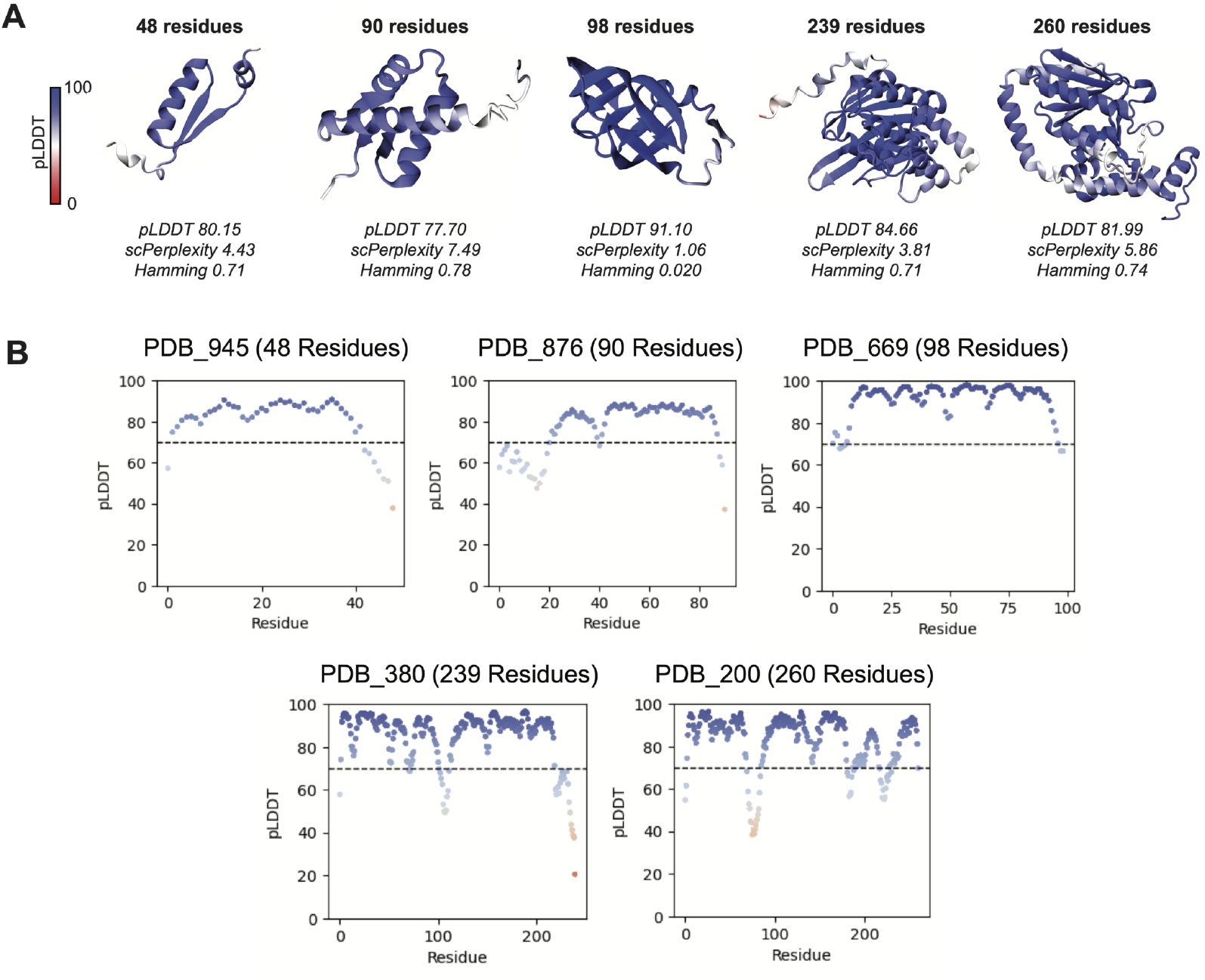
Representative proteins generated by EvoDiff-Seq. **(A)** Predicted structures and metrics for representative structurally plausible generations from EvoDiff-Seq, the 640M-parameter OADM model. **(B)** Corresponding per-residue pLDDT for pLDDT scores computed based on the OmegaFold predicted structures, for individual residues in representative high-fidelity generations from EvoDiff-Seq **(Fig. 2D)**. Points are colored by pLDDT (0-100, red to blue).

**Figure S6:**
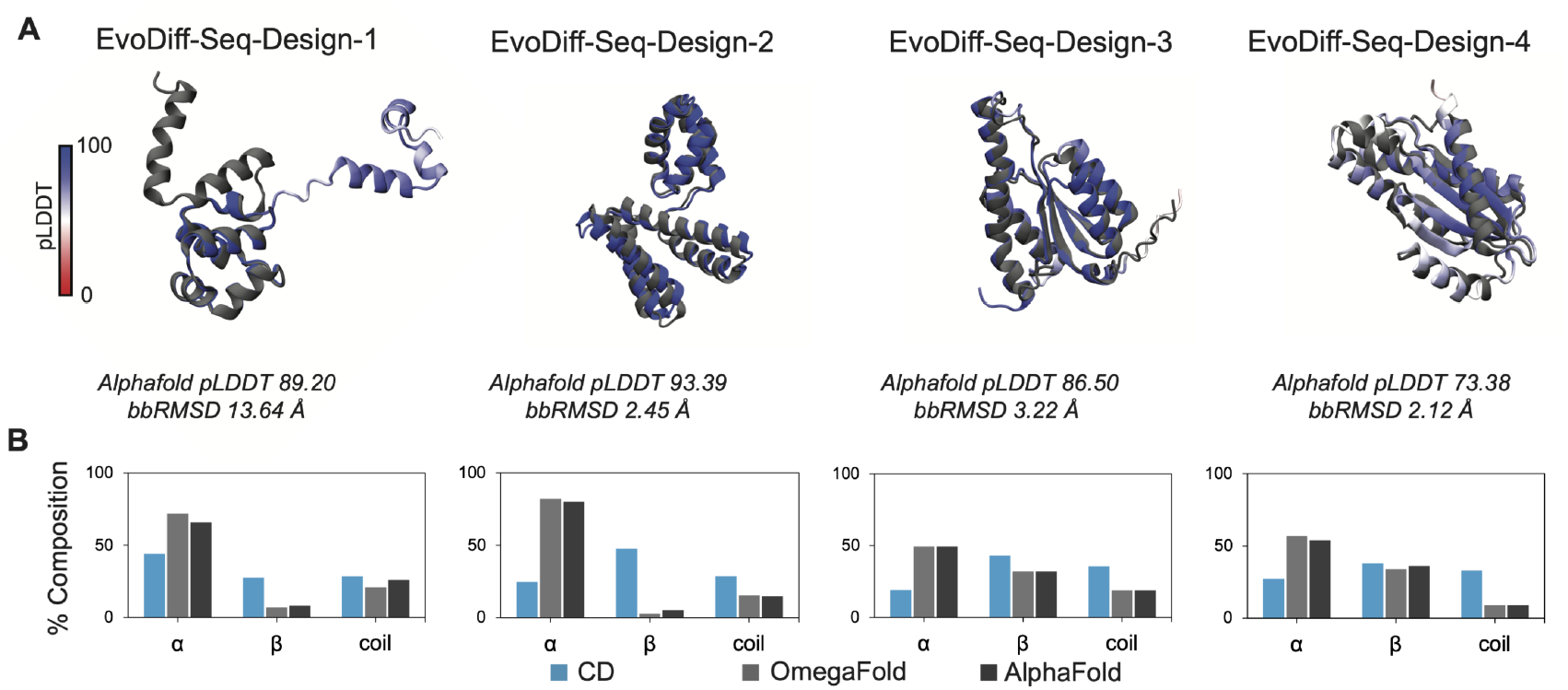
Comparison of structure-prediction methods for unconditional EvoDiff-Seq designs. **(A)** AlphaFold predictions (colored by pLDDT) overlaid on OmegaFold (grey). The AlphaFold pLDDT, and backbone RMSD (bbRMSD) taken between the two structures is denoted. **(B)** Percent composition extracted from CD spectra (blue) compared to structure prediction methods (AlphaFold and OmegaFold, grey).

**Figure S7:**
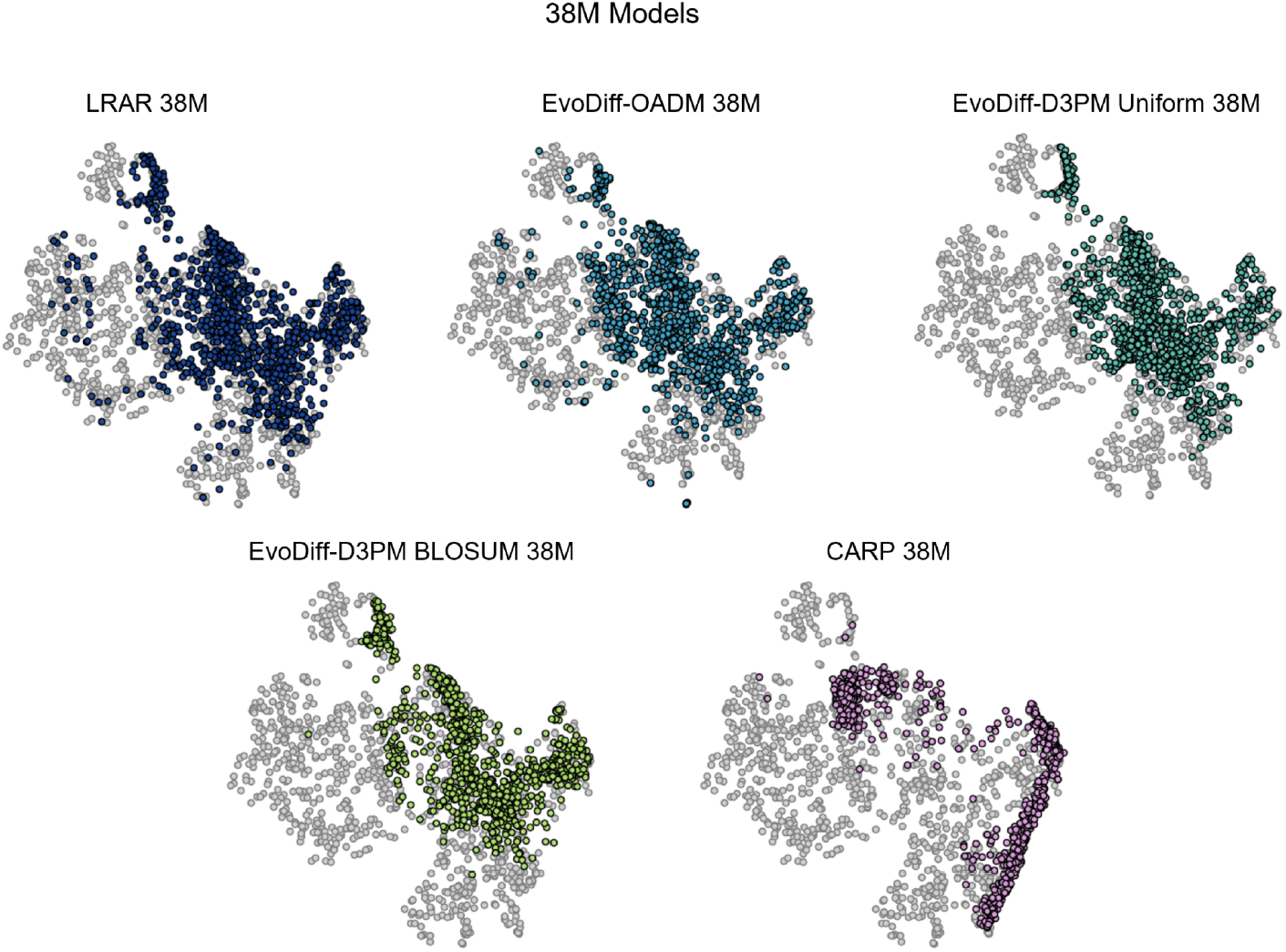
Coverage of sequence and functional space for generated distributions from 38M parameter EvoDiff sequence models and baselines. UMAP of ProtT5 embeddings, annotated with FPD, of natural sequences from test set (grey, *n=* l000 plotted) and of generated sequences from EvoDiff 38M parameter models and baselines (various colors, *n*=1000).

**Figure S8:**
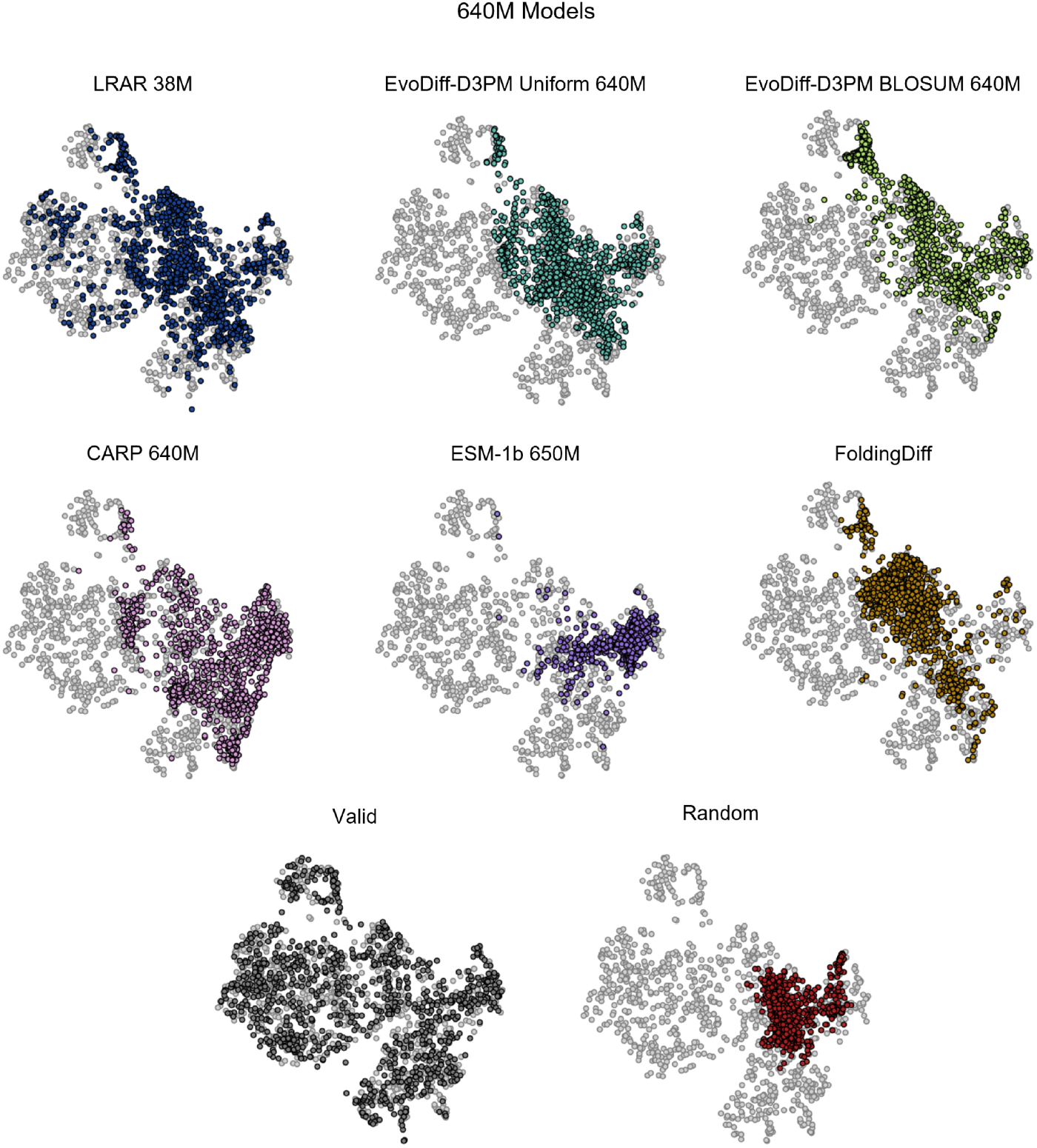
Coverage of sequence and functional space for generated distributions from 640M parameter EvoDiff sequence models and baselines. UMAP of ProtT5 embeddings, annotated with FPD, of natural sequences from test set (grey, *n=* 1000) and of generated sequences from EvoDiff 640M parameter models and baselines (various colors, *n*=l000). A visualization of sequences from the validation set (dark grey, *n=* 1000) is included for reference. The visualization for the 640M OADM model is excluded due to inclusion in Fig. 3A.

**Figure S9:**
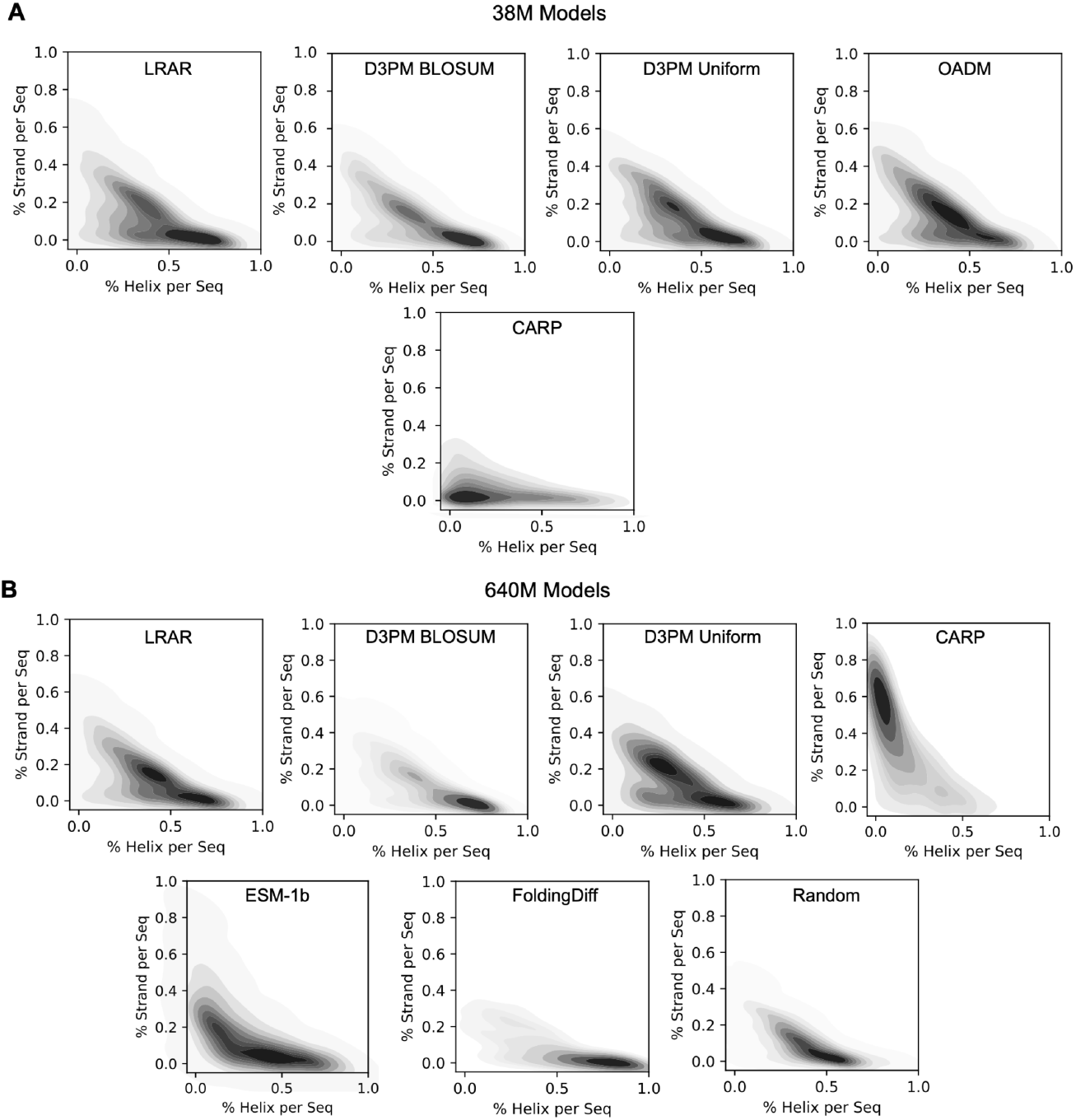
Structural features in generated sequences from all sequence models. **(A-B)** Multivariate distributions of helix and strand features in sequences from 38M (A) and 640M (B) parameter models, and baselines based on DSSP 3-state predictions and annotated with the KL divergence relative to the test set (*n=*1000 samples from each model). In (B), the distribution for the 640M OADM model is excluded (see Fig. 3B); the distribution for random sequences (*n=* 1000) is provided as reference.

**Figure S10:**
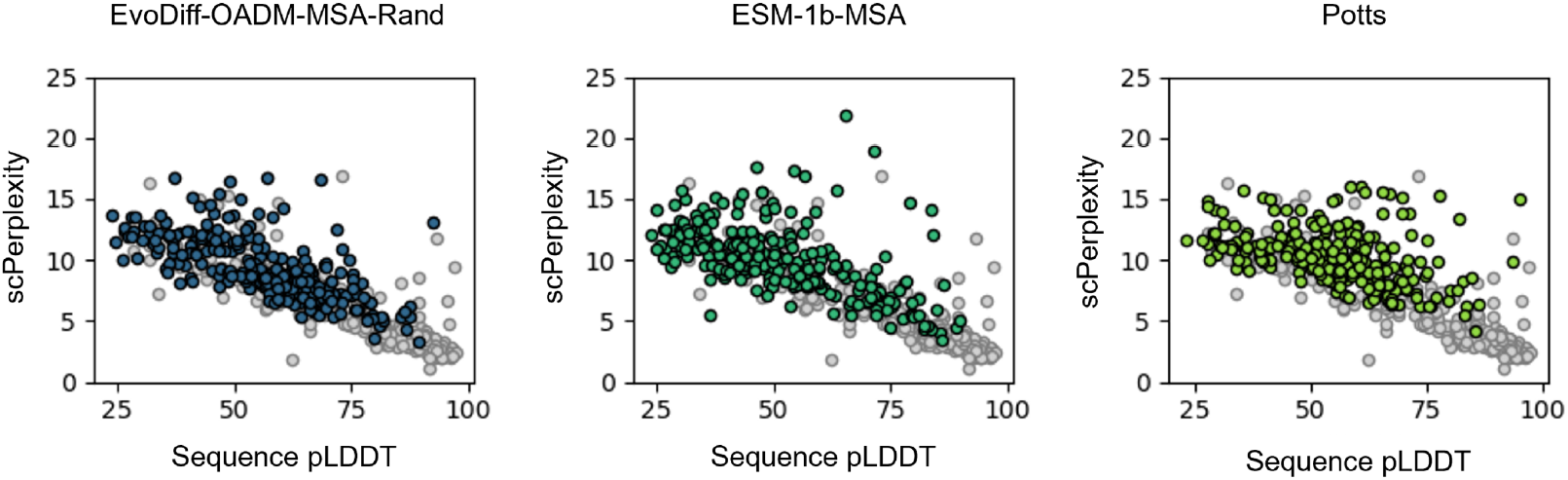
Sequence pLDDT versus scPerplexity for EvoDiff MSA models, for sequences from the validation set (grey, *n*=250) and evaluated MSA models (various colors, *n*=250), except for EvoDiff-OADM-MSA-Max (EvoDiff-MSA, shown in Fig. 4F).

**Figure S11:**
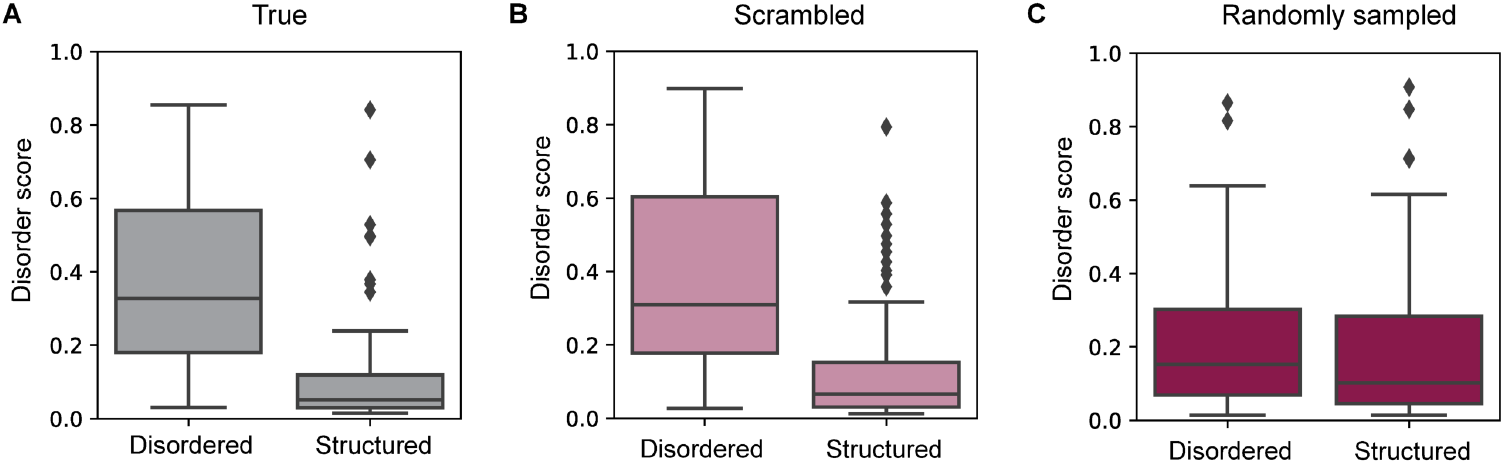
Baseline performance of DR-BERT evaluator. **(A-C)** Distributions of DR-BERT predicted disorder scores across disordered and structured regions for sequences with true (A), scrambled (B), and randomly sampled (C) IDRs (*n*=l00).

**Figure S12:**
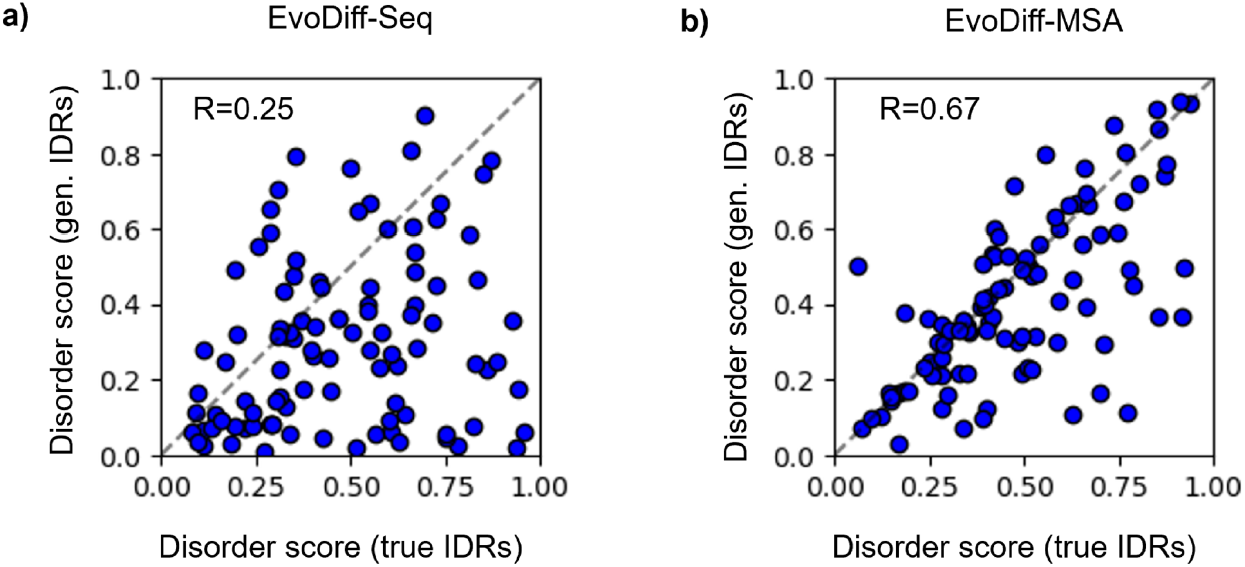
Performance of DR-BERT evaluator on disorder regions. **(A-B)** Disorder scores predicted by DR-BERT for true (x-axis) vs. generated (y-axis) IDRs for the same given sequence, for generations from EvoDiff-Seq (A) and EvoDiff-MSA (B). Each dot represents an individual IDR (*n*=l00). The Pearson R is given for each of EvoDiff-Seq and EvoDiff-MSA.

**Figure S13:**
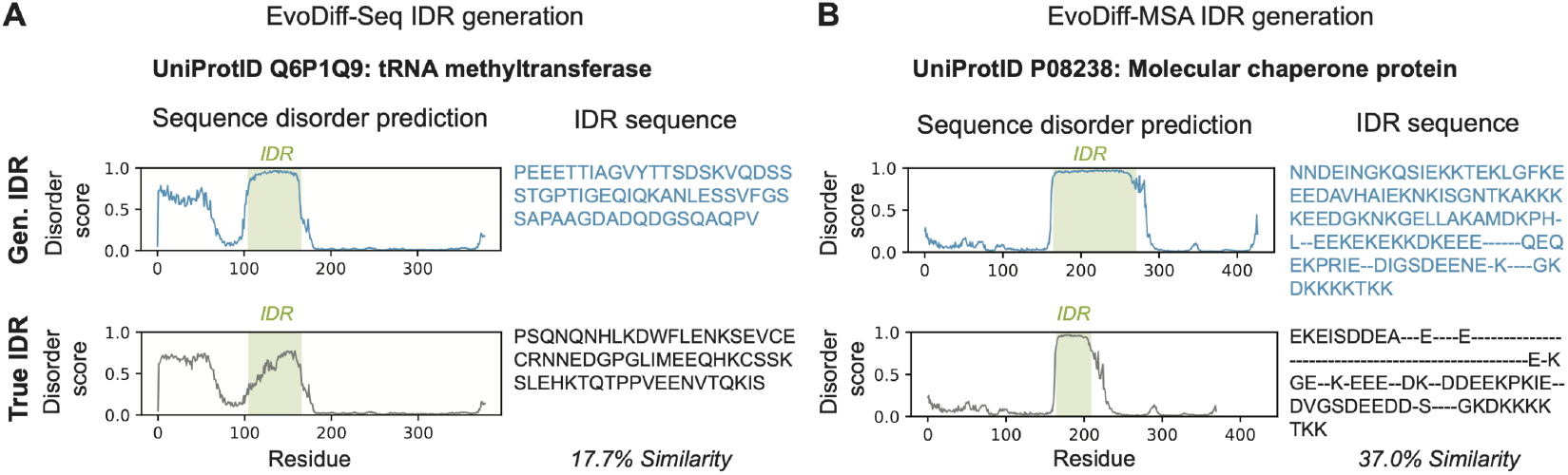
Examples of predicted disorder scores and corresponding sequences for representative generated (top row) and native (bottom row) IDRs from EvoDiff-Seq **(A)** and EvoDiff-MSA **(B)**.

**Figure S14:**
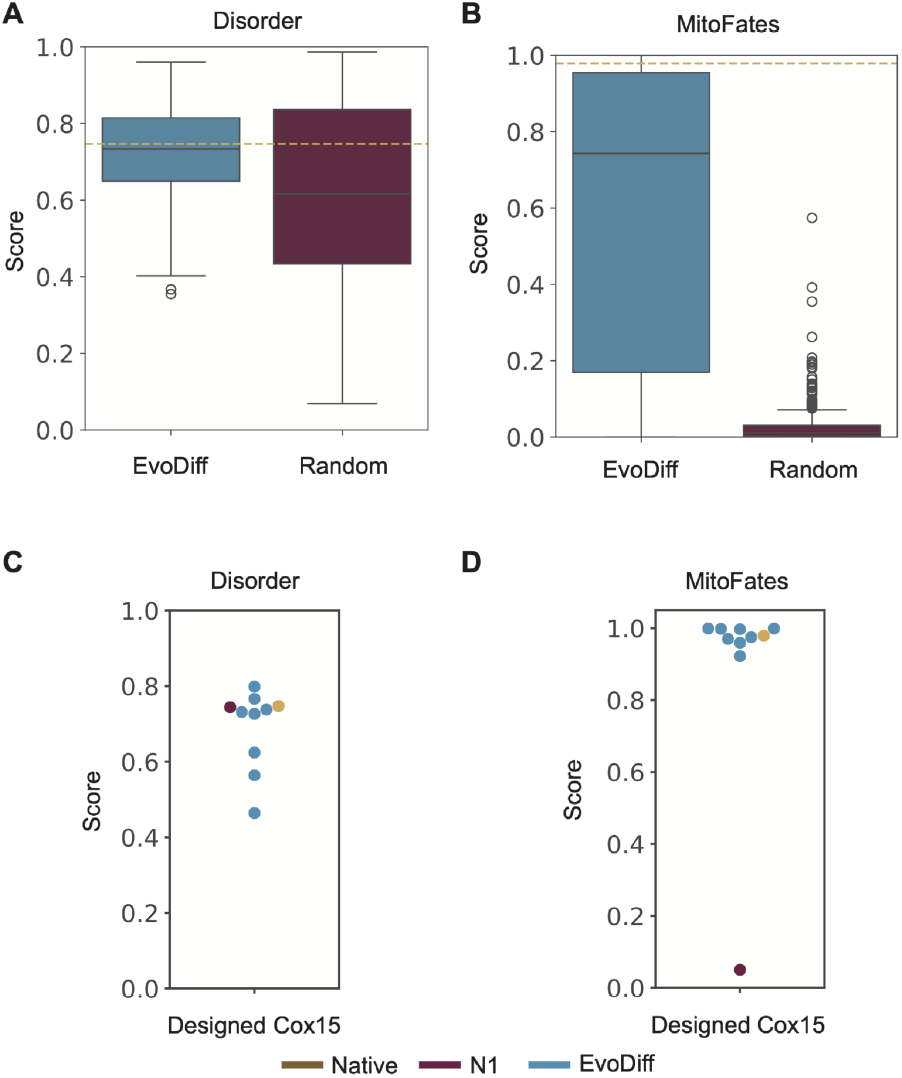
*In-silico* predictions for inpainted Cox15 designs. **(A)** Disorder scores predicted by DR-BERT and **(B)** MitoFates probability of presequences for 600 EvoDiff designs (blue) and 600 randomly sampled sequences (red). Scores for the native Cox15 IDR are denoted with a dashed yellow line. **(C)** Disorder and **(D**) Mitofates probability of presequence predictions for tested Cox15 IDRs, including EvoDiff designs (blue), native Cox15 IDR (yellow), and N1 (red).

**Figure S15:**
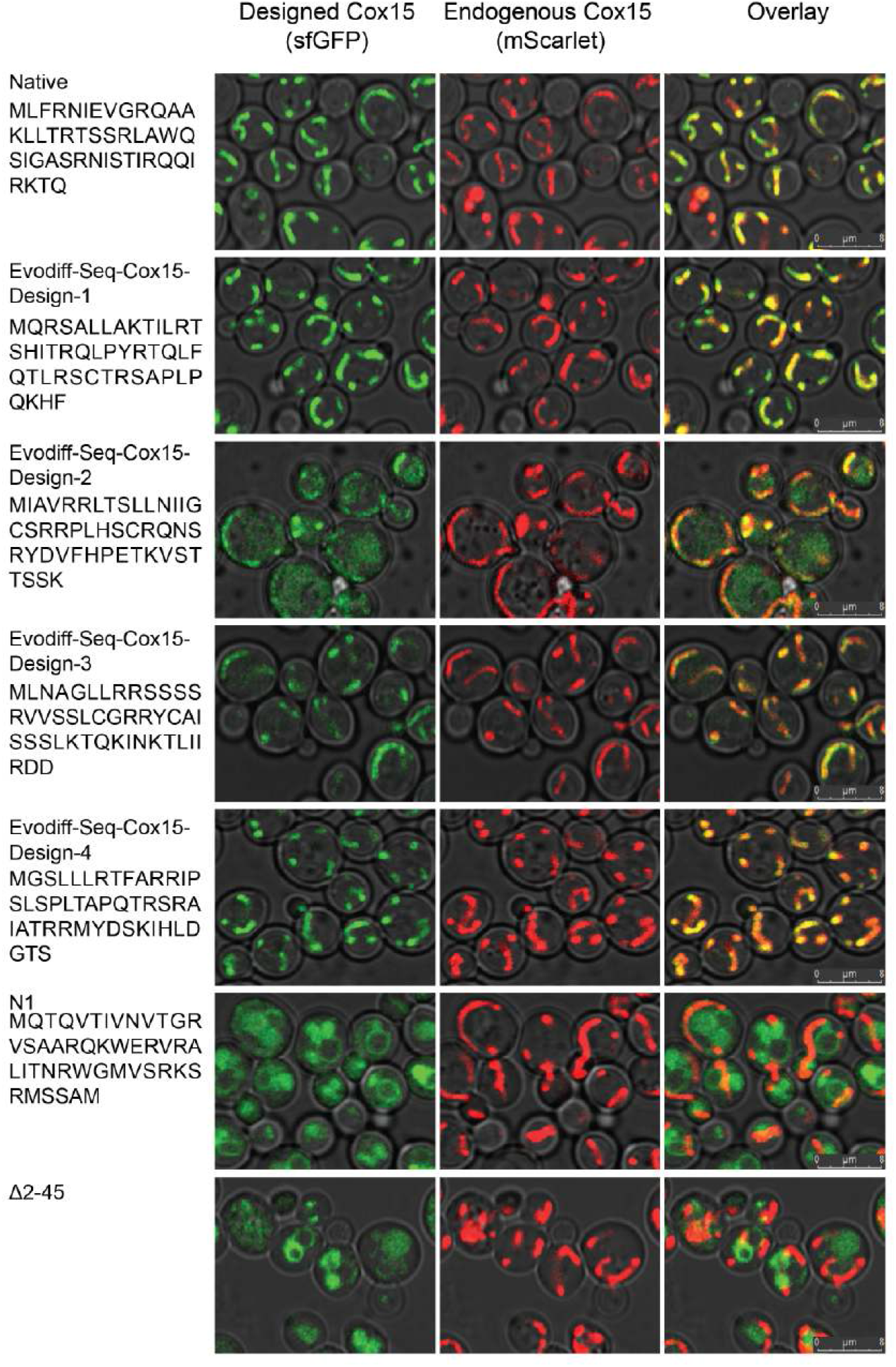
Fluorescence microscopy imaging of yeast cells for EvoDiff-Seq generated Designs. Columns from left to right show: the N-terminal IDR sequence used for a given Cox15-sfGFP design (rows); signal from the designed Cox15 (sfGFP. colored in green); signal from the endogenous Cox 15 (mScarlet, colored in red); overlaid images. All images show brightfield in grey, to visualize cell boundaries. Images for the native IDR, N1, and Δ2-45 are included for reference.

**Figure S16:**
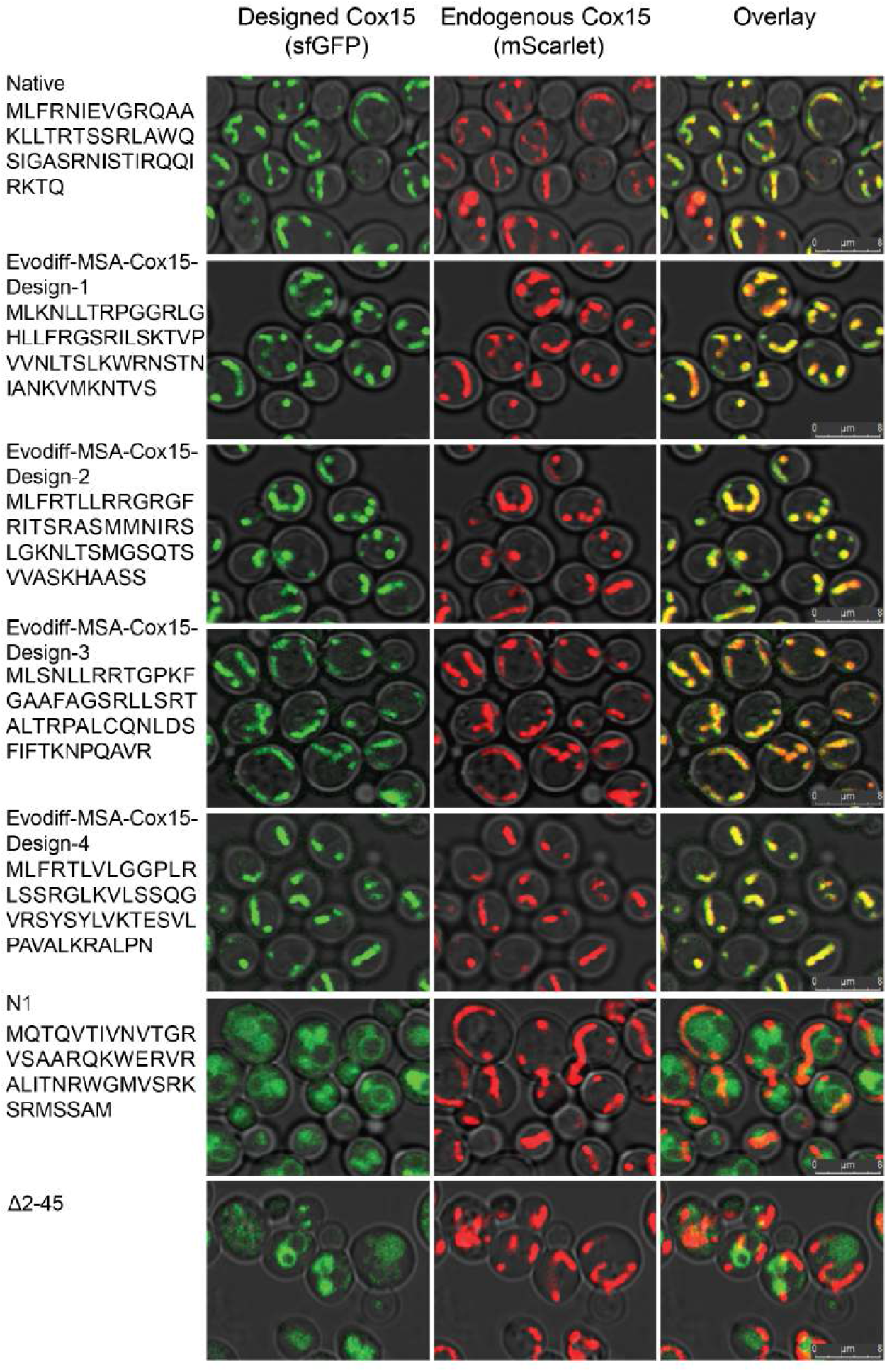
Fluorescence microscopy imaging of yeast cells for EvoDiff-MSA generated Designs. Columns from left to right show: the N-terminal IDR sequence used for a given Cox15-sfGFP design (rows); signal from the designed Cox15 (sfGFP, colored in green); signal from the endogenous Cox15 (mScarlet, colored in red); overlaid images. All images show brightfield in grey, to visualize cell boundaries. Images for the native IDR, N1, and Δ2-45 are included for reference.

**Figure S17:**
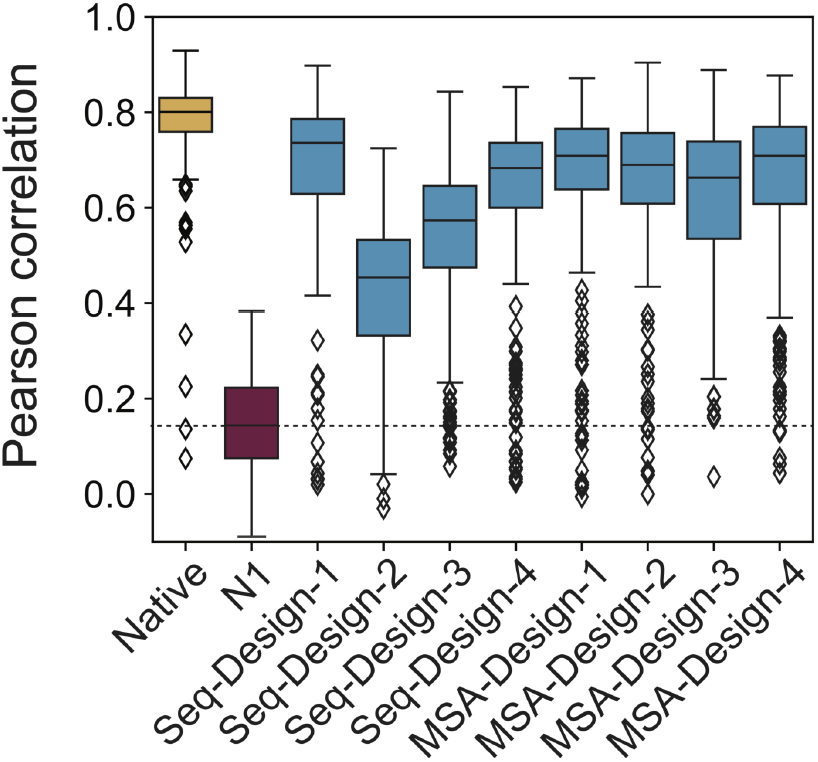
Per-cell pixel-wise Pearson correlation coefficients of fluorescence microscopy signal intensities between the mScarlet channel and the sfGFP channel. Independent replicates are measured over *n=*110-264 cells segmented by YeastSpotter for each design. The native IDR is shown in yellow, N1 in red, and EvoDiff designs in blue.

**Figure S18:**
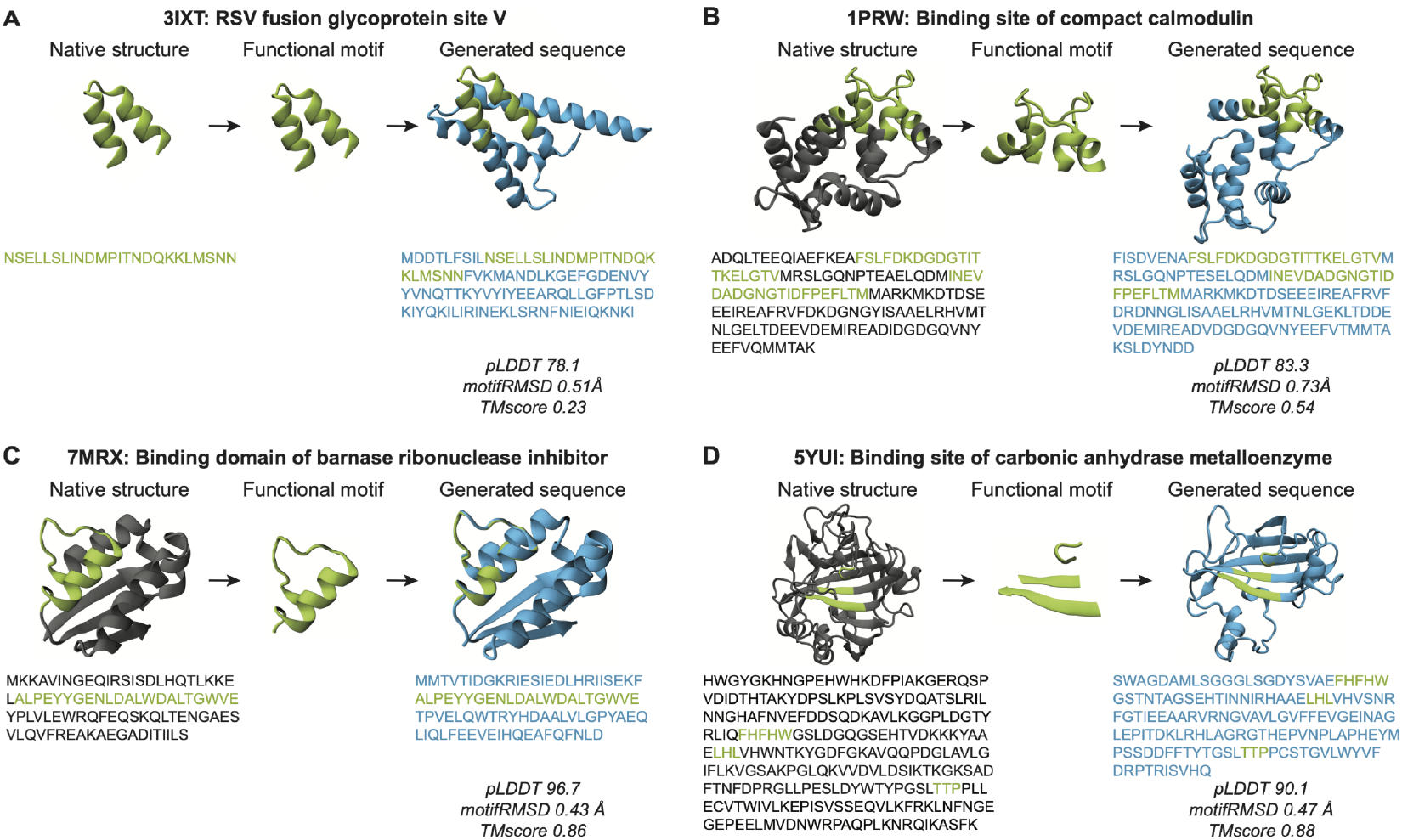
EvoDiff-Seq and EvoDiff-MSA scaffold binding domains in sequence space. **(A-D)** Generated sequences, predicted structures, and computed metrics for representative scaffolding examples from EvoDiff-Seq **(A/B)** and EvoDiff-MSA **(C/D)**. Motif is shown in green, original scaffold in gray, and generated scaffold in blue.

**Figure S19:**
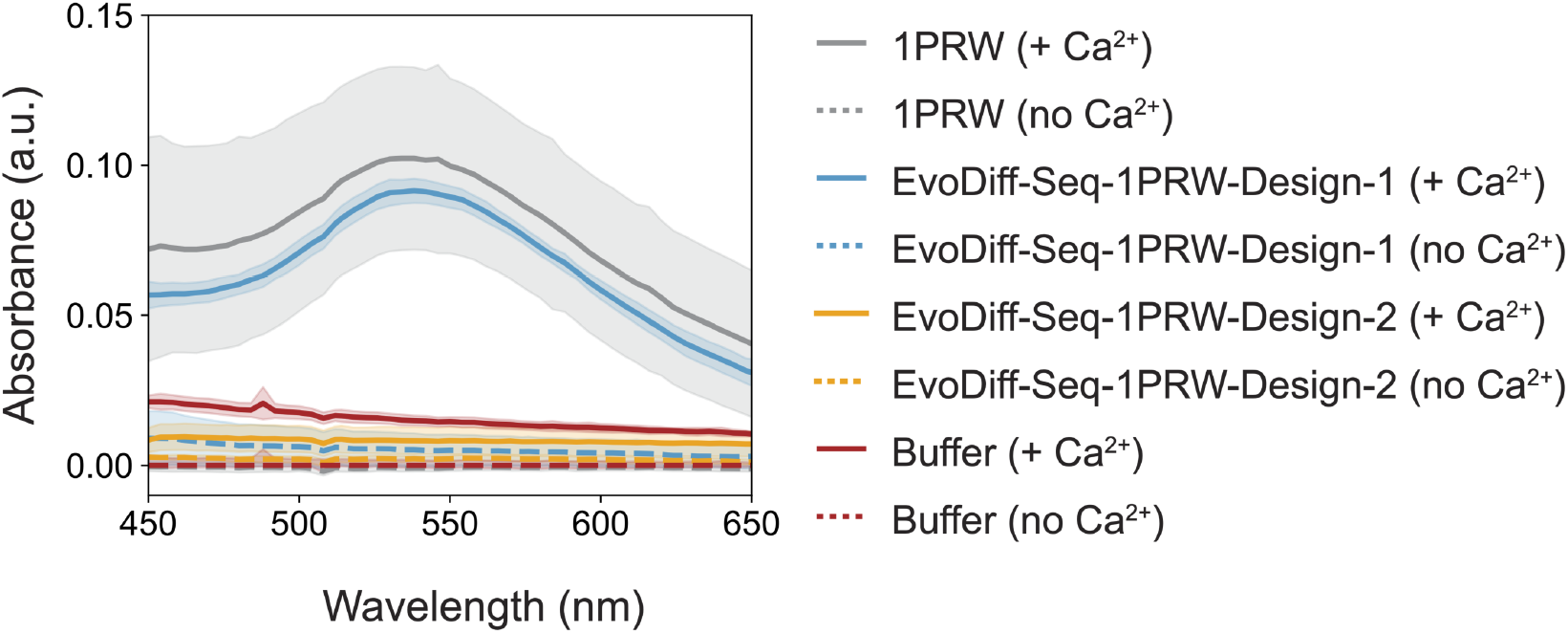
Calmodulin binding assay. Included is the native IPRW protein (grey), successful EvoDiff design (EvoDiff-Seq-lPRW-Design-l, blue), an unsuccessful EvoDiff design (EvoDiff-Seq-lPRW-Design-2, orange), and buffer (red), with (solid-line) and without (dashed-line) Ca^2^+. The mean value is reported, and error bars are represented by the std. dev taken from three independent trials

**Figure S20:**
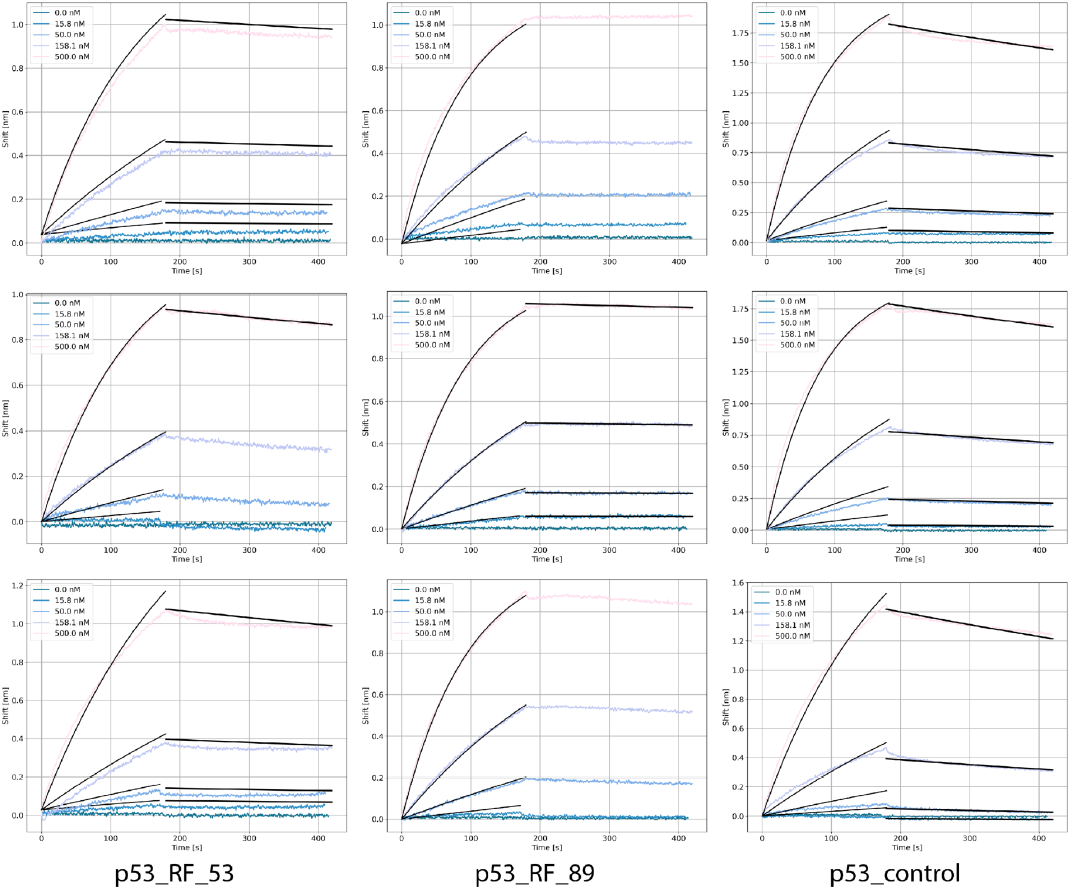
BLI results for positive controls.

**Figure S21:**
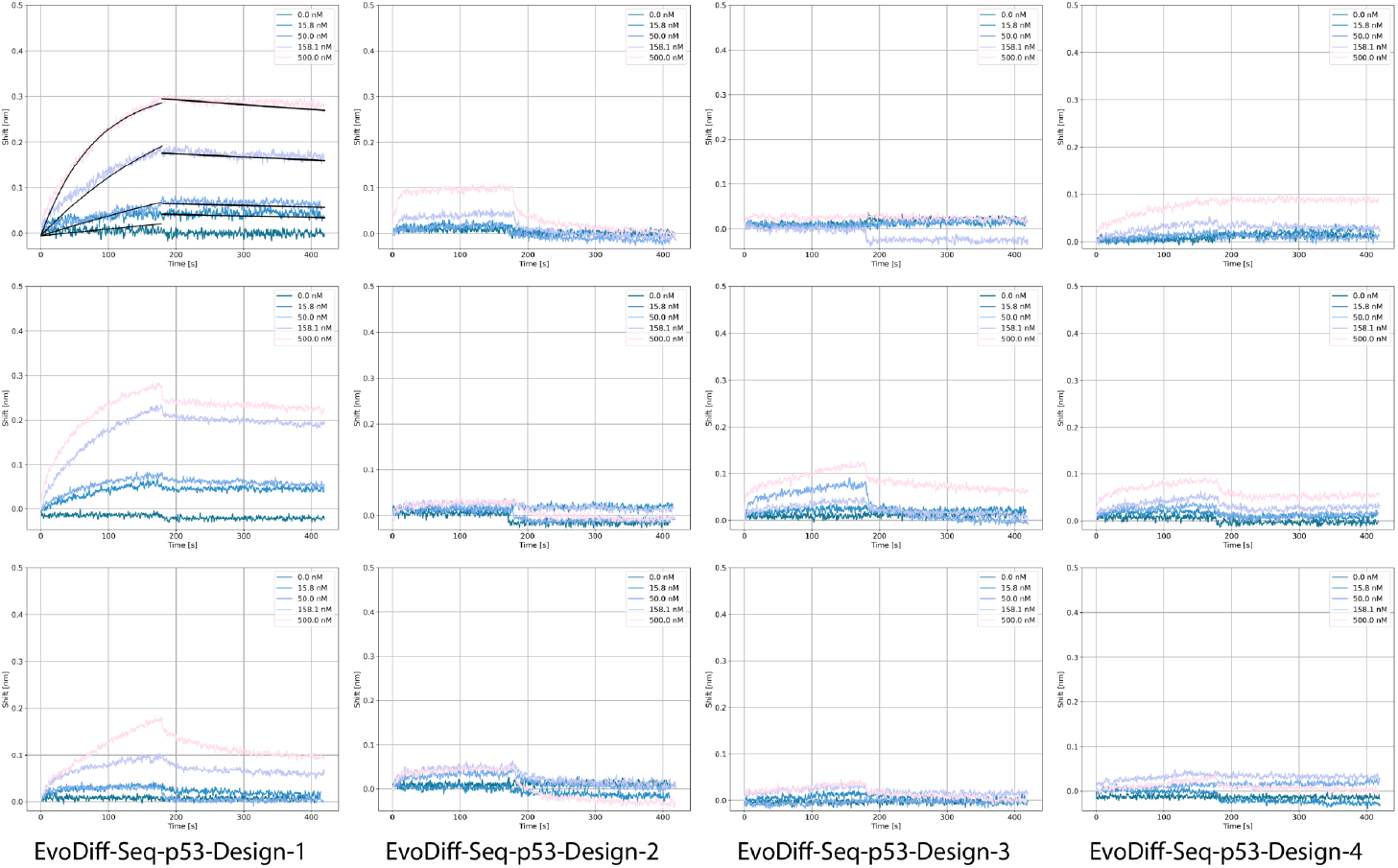
BLI results for successful MDM2 binders generated by EvoDiff-Seq.

**Figure S22:**
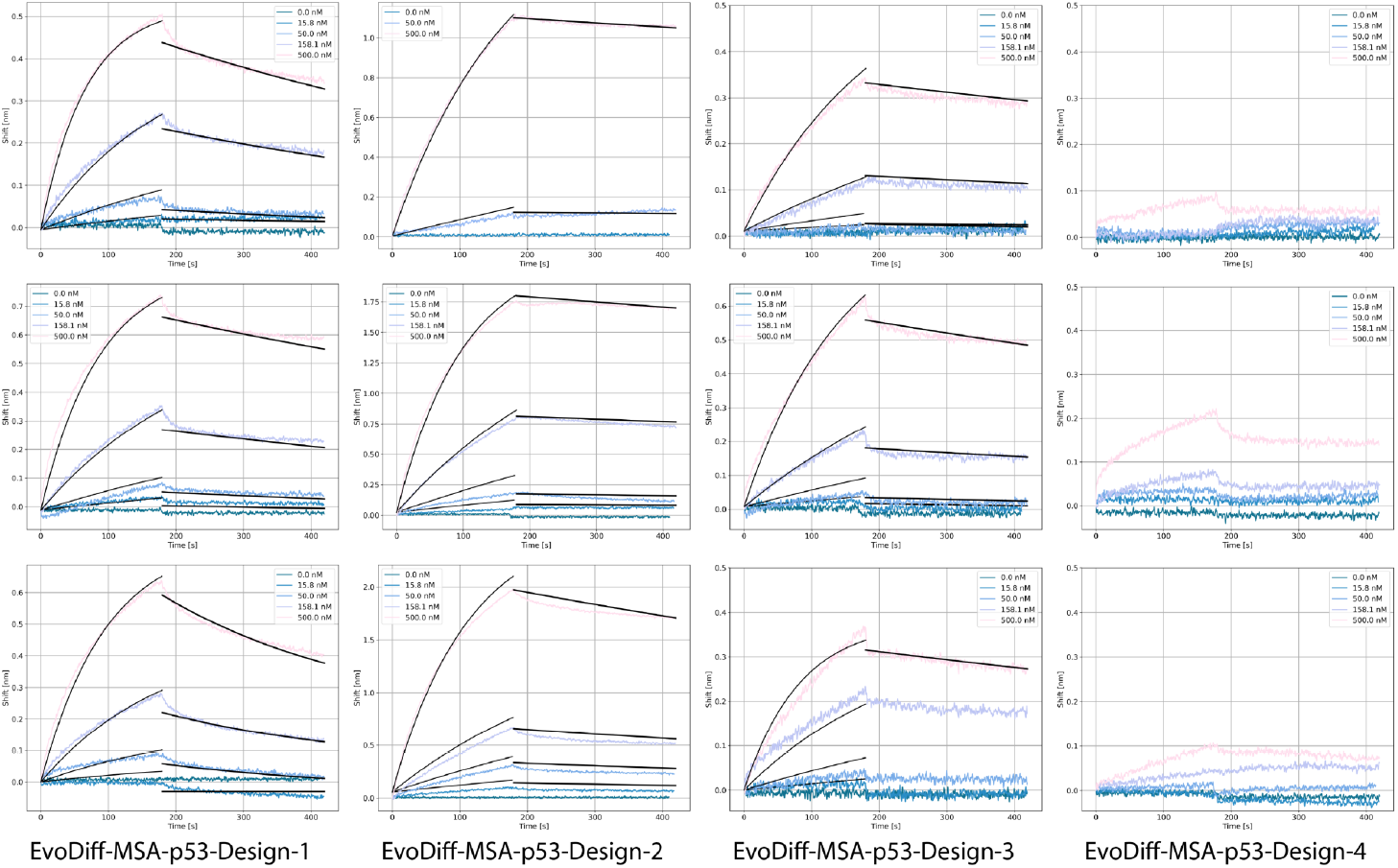
BLI results for successful MDM2 binders generated by EvoDiff-MSA.

**Figure S23:**
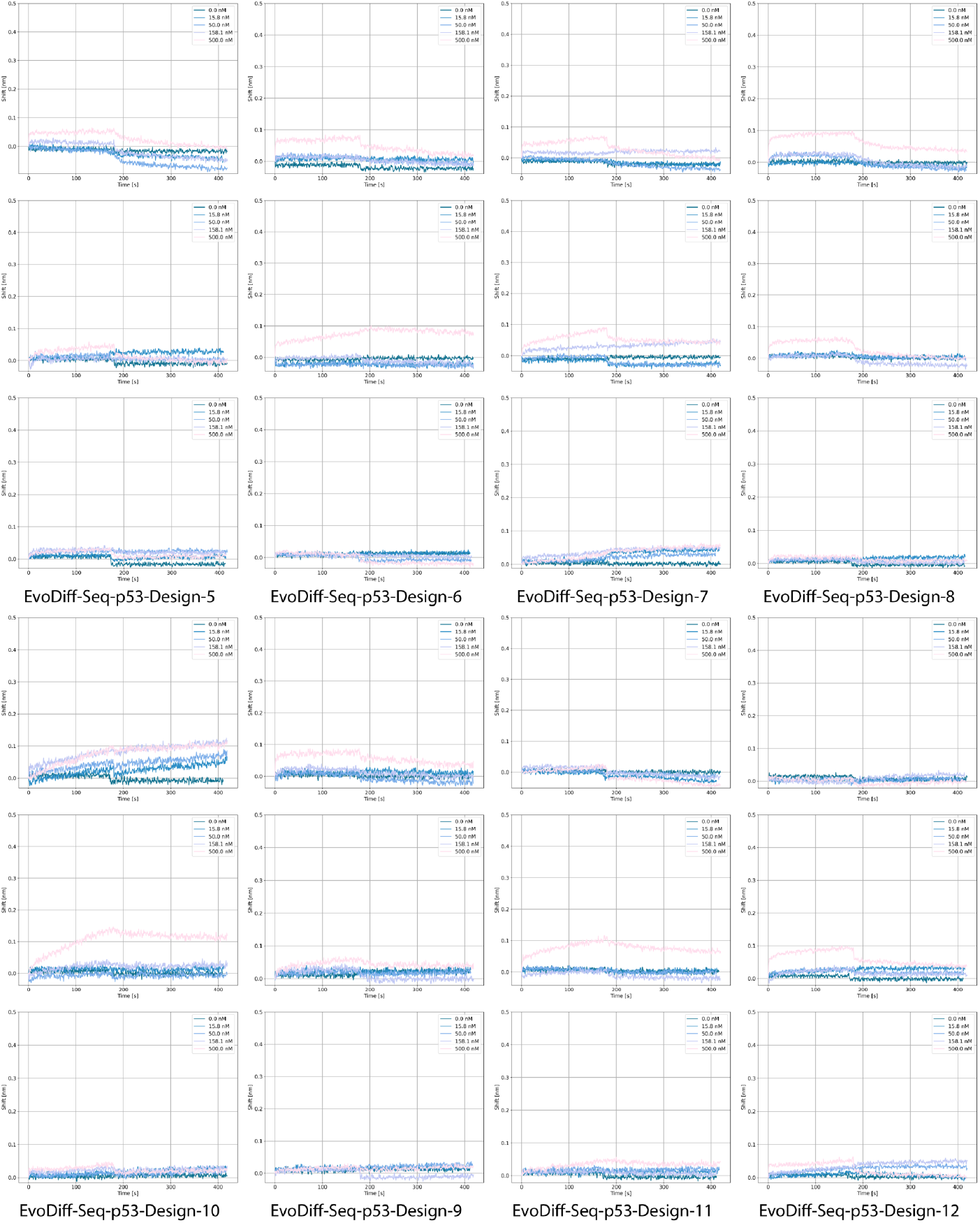
BLI results for unsuccessful MDM2 binders EvoDiff-Seq-p53-Design-5 through EvoDiff-Seq-p53-Design-12

**Figure S24:**
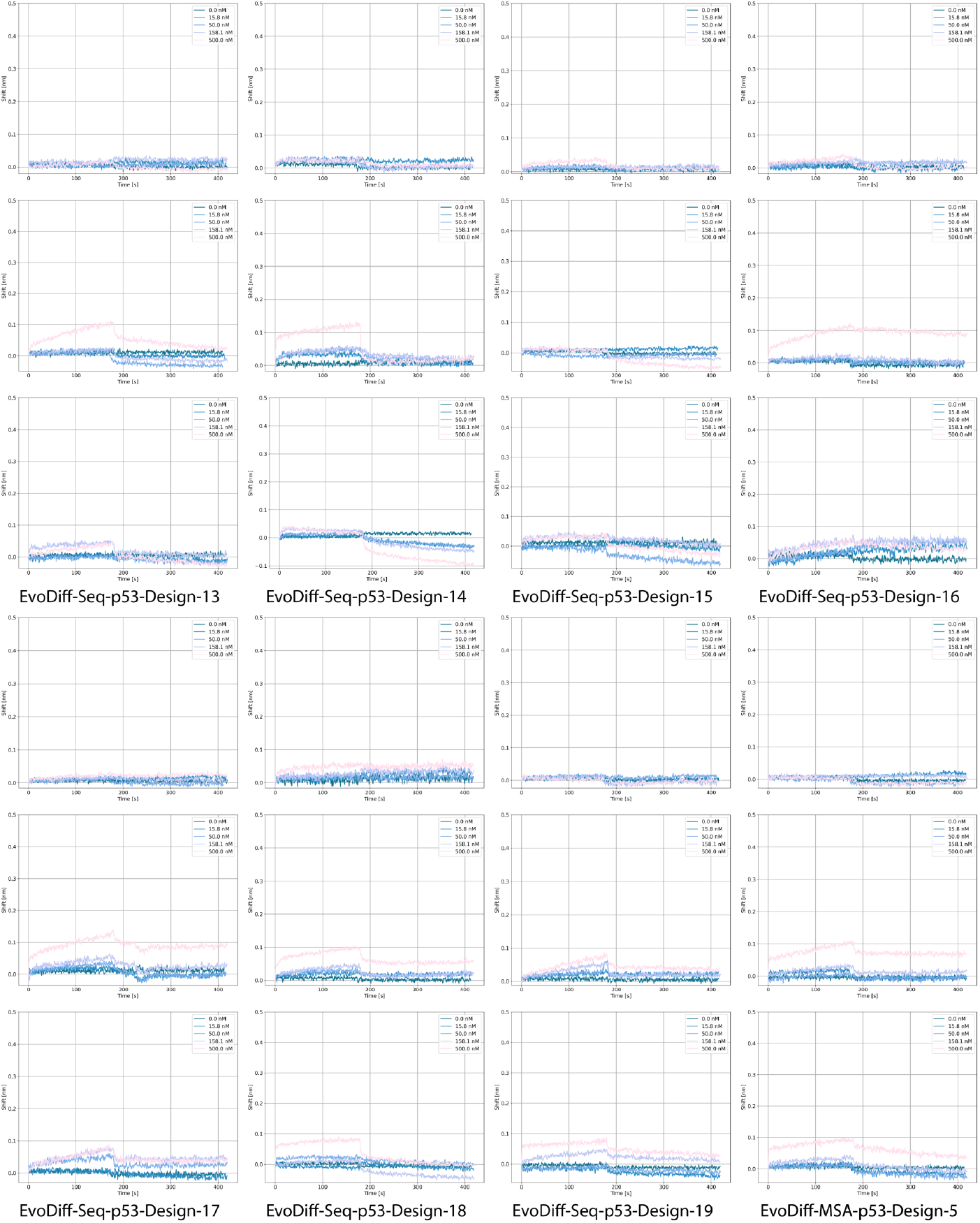
BLI results for unsuccessful MDM2 binders EvoDiff-Seq-p53-Design-13 through EvoDiff-MSA-p53-Design-5

**Figure S25:**
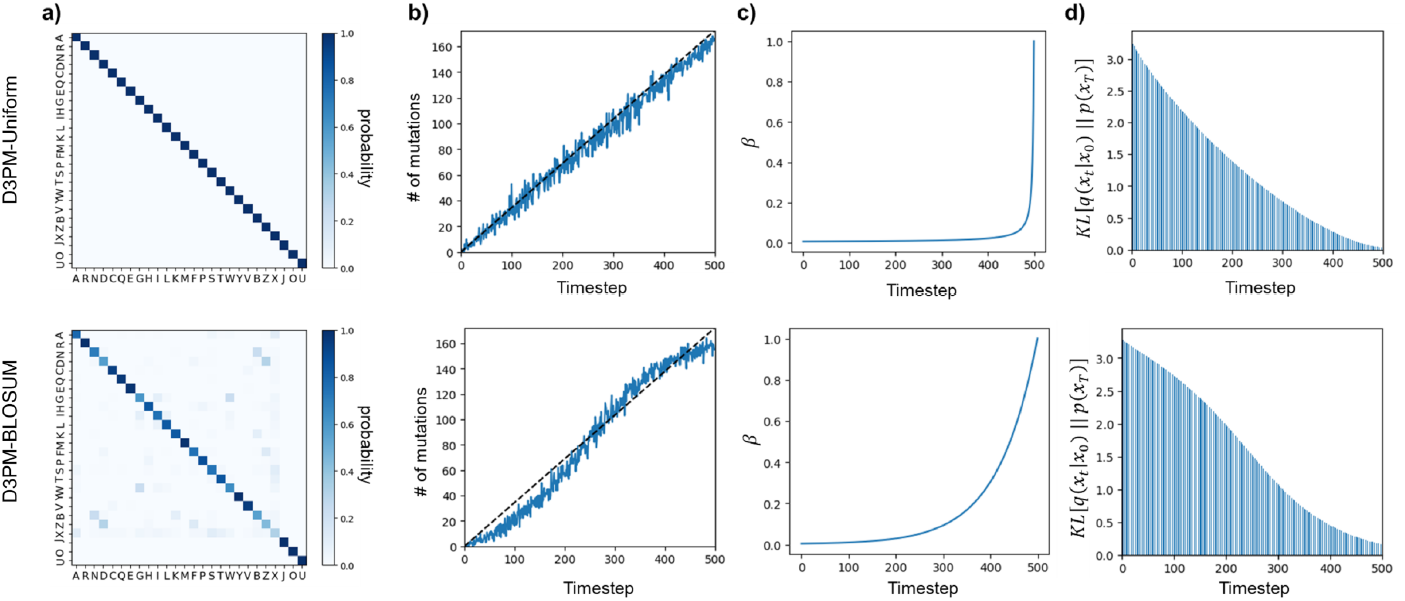
Details of EvoDiff-D3PM corruption schemes. The top and bottom rows correspond to EvoDiff-D3PM-Uniform and EvoDiff-D3PM-BLOSUM, respectively. **(A)** Visualization of EvoDiff-D3PM transition matrices. **(B)** Evolution of the number of mutations accrued as a function of the diffusion timestep *t* for a sample input. **(C)** Evolution of *β* as a function of the diffusion timestep *t*. **(D)** Evolution of *D*_*KL*_ [*q*(*x*_*t*_|*x*_0_)‖*p(x*_*T*_)] as a function of the diffusion timestep *t*, indicating convergence to a uniform stationary distribution at *t* = 500 as *D*_*KL*_ approaches zero.|

**Table S1:** Performance of EvoDiff sequence models. The reconstruction KL (Recon KL) was calculated between the distribution of amino acids in the test set and in generated samples (*n*=1000). The perplexity was computed on 25k samples from the test set. The minimum Hamming distance to any train sequence of the same length (Hamming) is reported for each model as the mean ± standard deviation over the generated samples. Fréchet ProtT5 distance (FPD) was calculated between the test set and generated samples. The secondary structure KL (SS KL) was calculated between the means of the predicted secondary structures of the test and generated samples.

| Model | parameters | Recon KL | perplexity | Hamming | FPD | SS KL |
| --- | --- | --- | --- | --- | --- | --- |
| Test | - | 9.92e-4<sup>1</sup> | - | 0.0039<sup>2</sup> | 0.10<sup>1</sup> | 1.37e-5<sup>1</sup> |
| D3PM BLOSUM | 38M | 1.77e-2 | 17.16 | 0.83 ± 0.05 | 1.42 | 3.30e-5 |
| D3PM Uniform | 38M | 1.48e-3 | 18.82 | 0.83 ± 0.05 | 1.31 | 3.73e-5 |
| OADM | 38M | 1.11e-3 | 14.61 | 0.83 ± 0.07 | 0.92 | 1.61e-4 |
| D3PM BLOSUM | 640M | 3.73e-2 | 15.74 | 0.83 ± 0.05 | 1.53 | 4.96e-4 |
| D3PM Uniform | 640M | 2.90e-3 | 18.47 | 0.83 ± 0.05 | 1.35 | 2.13e-4 |
| OADM | 640M | 1.26e-3 | 13.05 | 0.83 ± 0.08 | 0.88 | 1.48e-4 |
| LRAR | 38M | 7.90e-4 | 12.38 | 0.82 ± 0.06 | 0.86 | 1.61e-4 |
| CARP | 38M | 5.71e-1 | 25.13 | 0.74 ± 0.07 | 6.30 | 2.72e-3 |
| LRAR | 640M | 7.01e-4 | 10.41 | 0.83 ± 0.06 | 0.63 | 1.76e-5 |
| CARP | 640M | 3.56e-1 | 31.77 | 0.84 ± 0.05 | 1.78 | 5.03e-3 |
| ESM-1b <sup>3</sup> | 650M | 4.91e-1 | 53.49 | 0.83 ± 0.06 | 6.67 | 5.48e-4 |
| ESM-2 <sup>3</sup> | 650M | 5.00e-1 | 68.39 | 0.84 ± 0.06 | 6.79 | 3.05e-3 |
| FoldingDiff <sup>4</sup> | 14M | 5.49e-2 | - | - | 1.64 | 1.76e-3 |
| RFdiffusion <sup>5</sup> | 60M | 7.19e-2 | - | - | 1.96 | 5.98e-3 |
| Random | - | 1.65e-1 | 20 | 0.85 ± 0.04 | 3.16 | 1.90e-4 |

<sup>1</sup> Calculated between the test set and validation set.
<sup>2</sup> Reported value is the minimum Hamming distance between any two natural sequences of the same length in UniRef50.
<sup>3</sup> Due to model constraints, the maximum sequence length sampled was 1022.
<sup>4</sup> For the FoldingDiff baseline, 1000 structures generated by FoldingDiff were randomly selected, and the corresponding 1000 inferred sequences were inverse-folded using ESM-IF. These sequences are between lengths of 50 and 128 residues.
<sup>5</sup> For the RFdiffusion baseline, 1000 structures were generated corresponding to the UniRef train distribution length, and 1000 corresponding sequences were inverse-folded using ESM-IF.

**Table S2:** Validation-set perplexities for EvoDiff MSA models. The perplexity is calculated based on the ability of each model to reconstruct a subsampled MSA from the validation set. “Max Perplexity” and “Rand. Perplexity” indicate MaxHamming and Random subsampling, respectively, for construction of the validation MSA.

| Corruption | Subsampling | Params | Max Perplexity | Rand. Perplexity |
| --- | --- | --- | --- | --- |
| D3PM BLOSUM | Random | 100M | 11.35 | 8.31 |
| D3PM BLOSUM | Max | 100M | 10.98 | 7.61 |
| D3PM Uniform | Random | 100M | 10.14 | 6.77 |
| D3PM Uniform | Max | 100M | 10.06 | 6.66 |
| OADM | Random | 100M | 6.05 | 3.64 |
| OADM | Max | 100M | 6.14 | 3.60 |
| ESM-MSA-1b | Max | 100M | 11.20 | 5.89 |

**Table S3:**
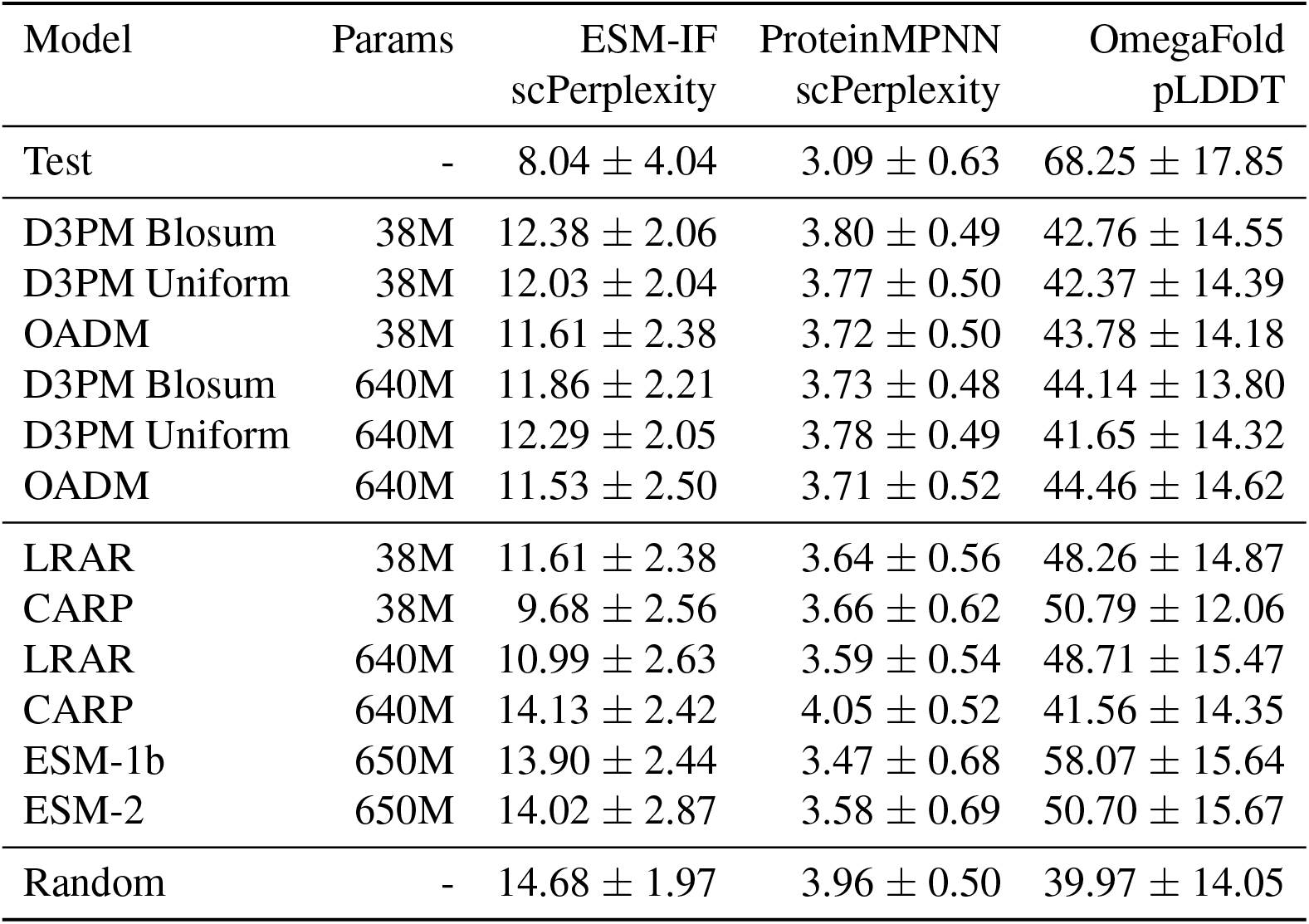
Structural plausibility metrics for EvoDiff sequence models and baselines. Metrics are reported as the mean ± standard deviation for 1000 generated samples for each model.

**Table S4:**
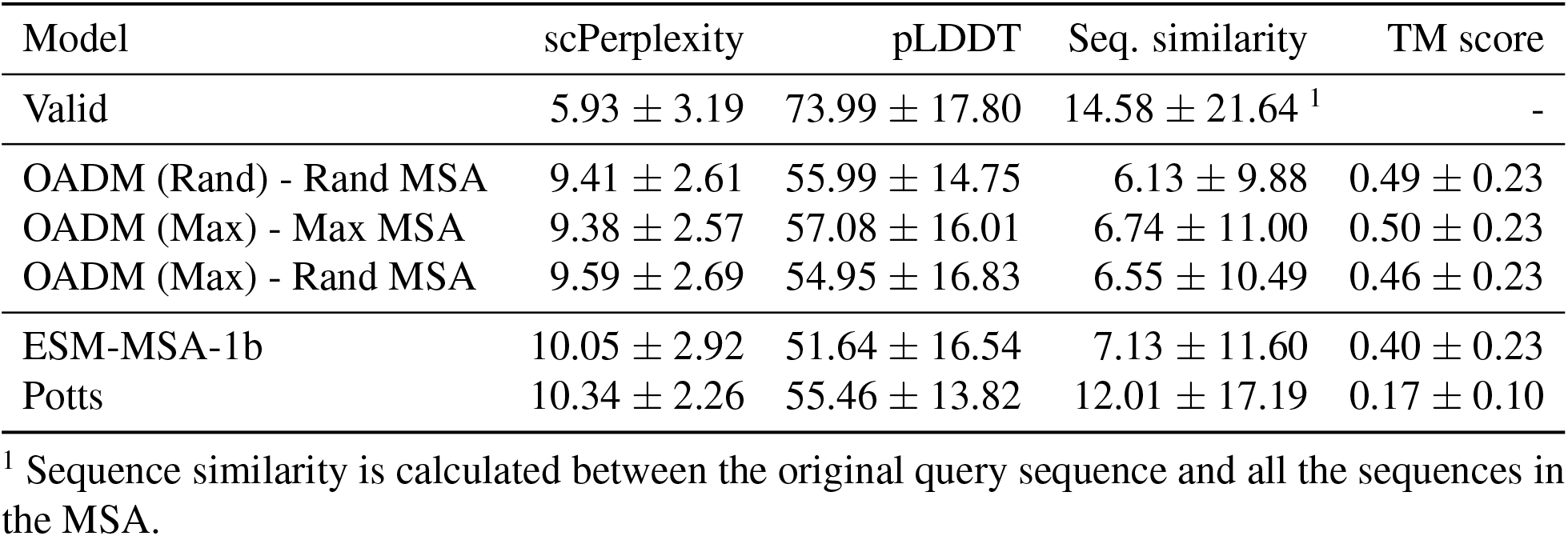
Performance of EvoDiff MSA models in generating query sequences conditioned on MSAs. Metrics are reported as the mean ± standard deviation over 250 generated samples for each model.

| Model | scPerplexity | pLDDT | Seq. similarity | TM score |
| --- | --- | --- | --- | --- |
| Valid | $5.93 \pm 3.19$ | $73.99 \pm 17.80$ | $14.58 \pm 21.64$ <sup>1</sup> | - |
| OADM (Rand) - Rand MSA | $9.41 \pm 2.61$ | $55.99 \pm 14.75$ | $6.13 \pm 9.88$ | $0.49 \pm 0.23$ |
| OADM (Max) - Max MSA | $9.38 \pm 2.57$ | $57.08 \pm 16.01$ | $6.74 \pm 11.00$ | $0.50 \pm 0.23$ |
| OADM (Max) - Rand MSA | $9.59 \pm 2.69$ | $54.95 \pm 16.83$ | $6.55 \pm 10.49$ | $0.46 \pm 0.23$ |
| ESM-MSA-1b | $10.05 \pm 2.92$ | $51.64 \pm 16.54$ | $7.13 \pm 11.60$ | $0.40 \pm 0.23$ |
| Potts | $10.34 \pm 2.26$ | $55.46 \pm 13.82$ | $12.01 \pm 17.19$ | $0.17 \pm 0.10$ |

<sup>1</sup> Sequence similarity is calculated between the original query sequence and all the sequences in the MSA.

**Table S5:**
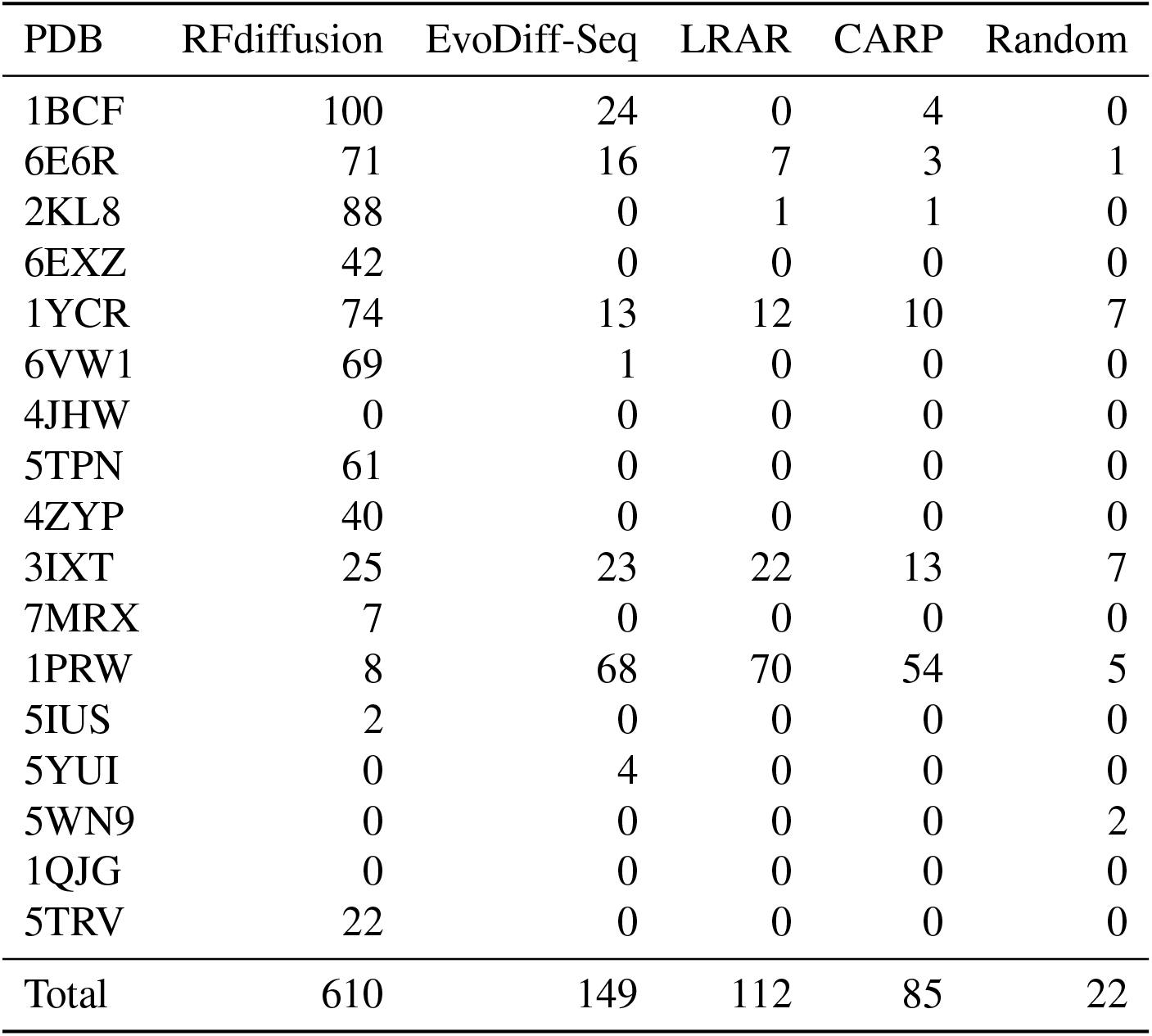
Scaffolding performance of EvoDiff-Seq. Number of scaffolding successes out of 100 generations for RFdiffusion, EvoDiff-Seq, the LRAR baseline, the CARP baseline, and randomly sampled scaffolds (Random), for each of 17 scaffolding problems. The bottom row contains the total number of successful scaffolds generated per model.

| PDB | RFdiffusion | EvoDiff-Seq | LRAR | CARP | Random |
| --- | --- | --- | --- | --- | --- |
| 1BCF | 100 | 24 | 0 | 4 | 0 |
| 6E6R | 71 | 16 | 7 | 3 | 1 |
| 2KL8 | 88 | 0 | 1 | 1 | 0 |
| 6EXZ | 42 | 0 | 0 | 0 | 0 |
| 1YCR | 74 | 13 | 12 | 10 | 7 |
| 6VW1 | 69 | 1 | 0 | 0 | 0 |
| 4JHW | 0 | 0 | 0 | 0 | 0 |
| 5TPN | 61 | 0 | 0 | 0 | 0 |
| 4ZYP | 40 | 0 | 0 | 0 | 0 |
| 3IXT | 25 | 23 | 22 | 13 | 7 |
| 7MRX | 7 | 0 | 0 | 0 | 0 |
| 1PRW | 8 | 68 | 70 | 54 | 5 |
| 5IUS | 2 | 0 | 0 | 0 | 0 |
| 5YUI | 0 | 4 | 0 | 0 | 0 |
| 5WN9 | 0 | 0 | 0 | 0 | 2 |
| 1QJG | 0 | 0 | 0 | 0 | 0 |
| 5TRV | 22 | 0 | 0 | 0 | 0 |
| Total | 610 | 149 | 112 | 85 | 22 |

**Table S6:** Scaffolding performance of EvoDiff-MSA. Number of scaffolding successes out of 100 generations for RFdiffusion, EvoDiff-MSA (Max), EvoDiff-MSA (Random), and the ESM-MSA baseline, for each of 17 scaffolding problems. The bottom row contains the total number of successful scaffolds generated per model.

| PDB | RFdiffusion | EvoDiff-MSA<br>(Max) | EvoDiff-MSA<br>(Random) | ESM-MSA |
| --- | --- | --- | --- | --- |
| 1BCF | 100 | 100 | 98 | 99 |
| 6E6R | 71 | 87 | 63 | 96 |
| 2KL8 | 88 | 11 | 31 | 42 |
| 6EXZ | 42 | 86 | 87 | 73 |
| 1YCR | 74 | 3 | 0 | 0 |
| 6VW1 | 69 | 4 | 3 | 4 |
| 4JHW | 0 | 0 | 0 | 0 |
| 5TPN | 61 | 0 | 0 | 0 |
| 4ZYP | 40 | 0 | 0 | 0 |
| 3IXT | 25 | 1 | 0 | 5 |
| 7MRX | 7 | 72 | 68 | 66 |
| 1PRW | 8 | 48 | 46 | 92 |
| 5IUS | 2 | 3 | 1 | 7 |
| 5YUI | 0 | 58 | 44 | 70 |
| 5WN9 | 0 | 0 | 0 | 0 |
| 1QJG | 0 | 34 | 22 | 38 |
| 5TRV | 22 | 15 | 12 | 12 |
| Total | 610 | 522 | 475 | 604 |

